# Microglial pruning of extinction-ensemble synapses preserves fear memory

**DOI:** 10.64898/2026.05.05.722833

**Authors:** Yong-Ling Wang, Yue Cao, Taohui Liu, Tian-Tian Shi, Yaoguang Zhao, Zhihao Jin, Xiaoyu Yang, Shuai-Yu Wang, Jia-Ze Ruan, Feng-Xia Zhang, Wei-Ke Li, Yang He, Sheng-Jie Lin, Wenying Xu, Xin Yi, Yan-Jiao Wu, Hengyue Shi, Jie Wang, Yuli Jiang, Ying-Xiao Liu, Xin-Ni Li, Tian-Lin Cheng, Tian-Le Xu, Bo Li, Peng Yuan, Bo Peng, Wei-Guang Li

## Abstract

Fear extinction suppresses learned fear without erasing the original memory, yet how competing fear and extinction ensembles are selectively updated remains unknown. Here, we show that microglia preserve fear memory by selectively editing extinction-ensemble synapses during retrieval. In medial prefrontal cortex, microglial processes, but not somata, expand, ramify, and tighten their engagement with extinction-ensemble dendrites and spines, where they preferentially engulf excitatory postsynaptic material and bias spine remodeling toward elimination. This selectivity is instructed by local find-me, eat-me, and don’t-eat-me cues: purinergic signaling recruits microglial processes, phosphatidylserine exposure licenses engulfment of extinction-ensemble synapses, and CD47-SIRPα protects fear ensembles from removal. Weakening microglial recruitment or engulfment, or removing this protection, accelerates extinction without impairing fear acquisition. These findings identify microglial processes as active gatekeepers of ensemble competition and reveal a neuroimmune mechanism that preserves fear memory by limiting extinction.

**In Brief:** During memory updating, microglial processes selectively engage extinction-ensemble synapses rather than globally pruning active circuits. Local recruitment, engulfment, and protection cues determine this choice, allowing microglia to preserve fear memory by limiting extinction.

**Highlights:**

1. Extinction retrieval selectively recruits microglial processes to extinction ensembles.
2. Microglia preferentially engulf excitatory postsynaptic material from extinction ensembles.
3. Local “find-me,” “eat-me,” and “don’t-eat-me” cues determine synapse selection.
4. Disrupting microglial pruning facilitates extinction without impairing fear learning.

## Introduction

Memories of threat are essential for survival and therefore tend to be robust and persistent. In modern human contexts, however, this adaptive property can become maladaptive, allowing traumatic memories to endure long after danger has passed. Fear extinction is distinct from forgetting: extinction is new learning that suppresses expression of the original fear memory, whereas forgetting reflects passive decay of an existing trace. Long-term memories are thought to reside in sparsely distributed neuronal ensembles that are reactivated during recall and modified by activity-dependent plasticity.^1-4^ In aversive learning, fear conditioning and extinction provide a tractable framework for studying how competing memory traces are formed, stabilized, and updated.^5-7^ Although fear and extinction are encoded by partially overlapping yet functionally distinct ensembles, the mechanisms that coordinately and differentially remodel these populations in the adult brain remain poorly understood.^8-12^ Understanding how these ensembles are regulated during fear learning and extinction is therefore important for improving interventions for post-traumatic stress disorder (PTSD).^13,14^

Microglia are the resident parenchymal macrophages of the central nervous system (CNS).^15,16^ They continuously survey local tissue and shape synaptic structure and function in an experience-dependent manner.^17-23^ Prior work suggests that microglia can either support or weaken memory, including through spine pruning, but findings across paradigms remain mixed.^22-27^ Although microglia eliminate synapses during development and in pathological states,^28-32^ how they engage specific neuronal ensembles during physiological memory updating remains unclear.^23,33^ A central unresolved question is whether microglia can bias competition between fear and extinction by selectively remodeling the synaptic substrates of one ensemble while sparing the other.

At neuron–microglia interfaces, local molecular cues can instruct when and where microglia engage synapses. These cues are often framed as “find-me” signals that recruit processes, “eat-me” signals that license engulfment, and “don’t-eat-me” signals that inhibit phagocytosis.^34,35^ This framework suggests that distinct neuronal ensembles may carry different molecular signatures that are read out by microglia to direct targeted synaptic remodeling. Under this view, microglia would not indiscriminately remove synapses but would instead interpret experience-dependent cues to bias pruning toward one ensemble while sparing the other, thereby shifting the balance between fear expression and extinction.

Here, using auditory fear conditioning in the medial prefrontal cortex (mPFC), a key node in both fear acquisition and extinction,^10,36-38^ we show that microglia preserve fear memory by selectively remodeling extinction ensembles during retrieval while largely sparing fear ensembles. This editing is carried out by microglial processes at soma-proximal, neuritic, and synaptic compartments and is instructed by distinct “find-me,” “eat-me,” and “don’t-eat-me” cues. These findings identify microglia as active gatekeepers of memory updating in the adult brain and suggest that locally targeted manipulation of microglial signaling may provide a route to enhance extinction in PTSD.

## Results

### Extinction retrieval selectively strengthens microglial process interfaces with neuronal ensembles

To determine whether microglia differentially engage memory-associated neuronal ensembles in the mPFC, we generated FosTRAP2::Ai14;;CX3CR1-GFP (Fos^wt/iCreER^::Ai14^wt/mut^;;CX3CR1^wt/GFP^) mice by crossing CX3CR1^wt/GFP^,^39^ FosTRAP2 (Fos2A-iCreER),^40^ and the Ai14 reporter mice.^41^ In this model, microglia are GFP-labeled and neurons activated within a defined 4-hydroxytamoxifen (4-OHT) window are permanently labeled with tdTomato. To avoid the potential influence from the CX3CR1^wt/GFP^ haplodeficiency,^42^ we verified some key conclusions in FosTRAP2::Ai14 (Fos^wt/iCreER^::Ai14^wt/mut^) mice by crossing FosTRAP2 and Ai14 mice. To label ensembles associated with fear learning, mice underwent auditory fear conditioning (conditioned stimulus–unconditioned stimulus pairings) and received 4-OHT immediately after training, followed by fear retrieval on day 7; CS-only controls received identical tone presentations without shock and showed minimal freezing (Figures 1A, 1B, and S1A). To tag extinction-associated ensembles, a separate cohort underwent extinction training in a distinct context, received 4-OHT after the final extinction session, and was tested at extinction retrieval on day 10 (Figures 1A, 1B, and S1B), consistent with incomplete suppression of conditioned fear.

**Figure 1.**
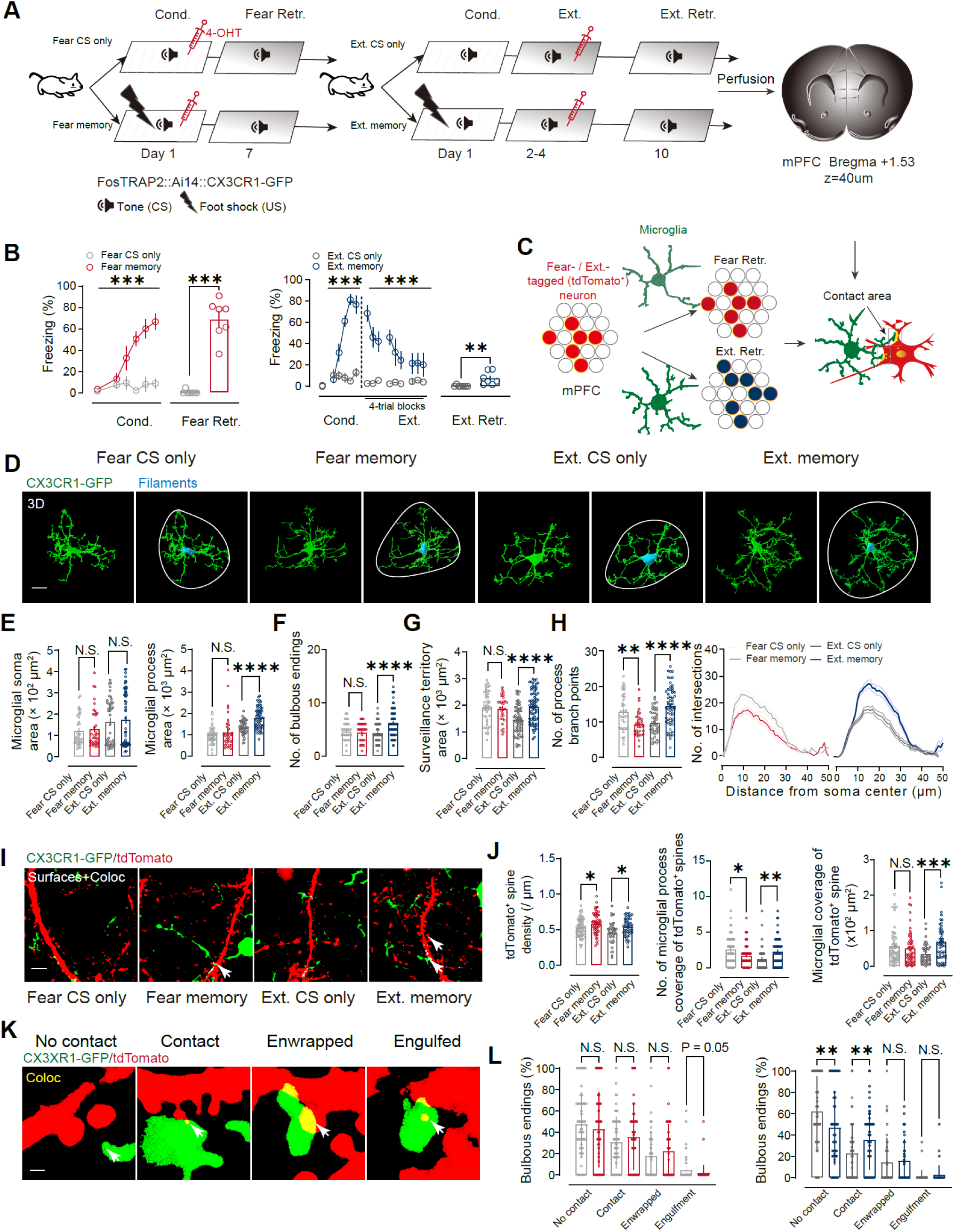
Extinction retrieval selectively strengthens compartment-specific microglia–ensemble interfaces, particularly at dendritic spines. (A) Ensemble-tagging strategy in the mPFC (coronal sections centered near Bregma +1.53) of CX3CR1FosTRAP2::Ai14;;CX3CR1-GFP mice. For fear ensembles, 4-OHT was administered immediately after auditory fear conditioning (Cond.; CS–US pairings) to TRAP-label fear-learning–activated neurons (tdTomato^+^), followed by a retrieval test on day 7 (Fear Retr.). For extinction ensembles, a separate cohort underwent extinction training (Ext.; days 2–4) and received 4-OHT immediately after the third extinction session to TRAP-label extinction-learning–activated neurons, followed by an extinction retrieval test on day 10 (Ext. Retr.). Brains were collected after the retrieval/CS-only session for imaging. (B) Freezing during CS presentations across conditioning and retrieval sessions (left) or conditioning, extinction training (4-trial blocks), and extinction retrieval (right). CS-only control groups received the same CS exposure without shock. Data are mean ± s.e.m. Fear CS only, *n* = 11 mice; Fear memory, *n* = 7 mice; Ext. CS only, *n* = 11 mice; Ext. memory, *n* = 11 mice. (C) Schematic of compartment-resolved quantification of microglia–ensemble interfaces (soma, neurites, and spines) in mPFC. (D) Representative 3D reconstructions of CX3CR1-GFP^+^ microglia (green) and corresponding Imaris Filament reconstructions (cyan). White outline denotes the surveillance territory defined by the outermost process tips. Scale bar, 10 μm. (E to H) Quantification of microglial morphology from 3D reconstructions: soma area and total process area (E), number of bulbous endings (F), surveillance territory area (G), and arbor complexity by Sholl analysis (branch points and intersections) (H). Extinction retrieval increased process elaboration (process area, bulbous endings, surveillance territory, and branching) without changing soma size, whereas fear retrieval did not induce comparable process expansion and reduced branching complexity. Each dot represents one microglial cell; *n* = 29–72 cells from 7 mice per group (exact *n* varies by metric). (I) Leica Lightning super-resolution confocal z-stacks (effective lateral resolution ∼120 nm) showing tdTomato^+^ dendrites/spines (tdTomato, red) and CX3CR1-GFP^+^ microglial processes (green); arrows indicate microglia–spine appositions. Scale bar, 5 μm. (J) Spine-level engagement quantified from 3D reconstructions: tdTomato^+^ spine density, number of microglial processes apposed to tdTomato^+^ spines, and total microglia–spine contact area. Fear retrieval reduced microglial process apposition to tdTomato+ dendrites/spines without increasing contact area, whereas extinction retrieval increased both process apposition and contact area. (K) Operational classification of microglia–spine interactions based on 3D coverage: no contact, contact (<50% surface coverage), enwrapped (≥50% coverage), or engulfed. Scale bar, 0.5 μm. (L) Distribution of tdTomato^+^ spines across interaction categories for fear (left) and extinction (right) group. Extinction retrieval shifted tdTomato^+^ spines from the “no contact” to the “contact” category, with no detectable change in enwrapped or engulfed fractions; fear retrieval did not measurably redistribute spine categories. *n* = 42–58 dendritic segments (or equivalent analyzed units) from 7 mice per group (Fear CS-only, 58; Fear memory, 57; Ext. CS-only, 42; Ext. memory, 55). N.S., not significant; \**P* < 0.05, \*\**P* < 0.01, \*\*\**P* < 0.001. Statistics: two-way repeated-measures ANOVA for conditioning/extinction time courses (B) and two-tailed unpaired Student’s *t* tests for pairwise comparisons versus the corresponding CS-only controls unless otherwise indicated.

We first examined whether memory retrieval is accompanied by remodeling of microglial arbors. Using three-dimensional (3D) reconstructions of CX3CR1-GFP microglia, we quantified soma and process features across conditions (Figures 1C and 1D). Microglial soma area remained stable (Figure 1E), arguing against a gross amoeboid transformation. In contrast, extinction retrieval induced a pronounced process-expanded, highly ramified state, with increased total process area, more distal bulbous endings, and enlarged surveillance territory relative to extinction CS-only controls (Figures 1F–1H). Fear retrieval did not produce comparable process expansion and was associated with reduced arbor complexity by Sholl analysis (Figure 1H). These observations indicate that extinction retrieval preferentially engages a process-rich microglial configuration.

We next asked whether this altered morphology translates into stronger physical engagement with tagged neuronal ensembles. In FosTRAP2::Ai14 tissue immunostained for IBA1, we quantified the shortest 3D Euclidean distance from each tdTomato^+^ soma to the nearest microglial soma and measured microglial apposition onto the tdTomato^+^ soma surface. Retrieval of either fear or extinction memory reduced the distance between tdTomato^+^ neuronal bodies and the nearest microglia relative to the corresponding CS-only controls (Figures S1C–S1E), indicating that both memory states recruit microglia into closer proximity to the engaged ensembles. However, strengthening of the soma-level interface was memory selective: extinction retrieval increased microglial coverage of tdTomato^+^ neuronal bodies, whereas fear retrieval did not measurably increase somatic coverage; tdTomato^+^ body size was unchanged across groups (Figures S1F–S1I). We then quantified neuronal body- and neurite-level interfaces directly in the same FosTRAP2::Ai14;;CX3CR1-GFP sections using 3D surface rendering and voxel-based colocalization (Figure S2). tdTomato^+^ body size again remained unchanged (Figure S2C, left), but microglial coverage of tdTomato^+^ neuronal bodies—measured as interface area and fractional coverage—was reduced in the fear-memory group and increased in the extinction-memory group relative to their corresponding CS-only controls (Figure S2C). Neurite coverage showed the same directional divergence, decreasing with fear retrieval and increasing with extinction retrieval (Figure S2D). Consistent with these measurements, the number of microglial processes contacting each tdTomato^+^ body was lower after fear retrieval but higher after extinction retrieval (Figure S2E). Together, these data show that proximity alone is not sufficient to predict interface strength; rather, extinction retrieval selectively deepens microglial engagement at somatic and neuritic compartments.

To verify that the confocal-defined interfaces reflect bona fide physical contacts rather than optical overlap, we performed correlative light–electron microscopy (CLEM) in FosTRAP2::Ai14;;CX3CR1-GFP mice. Two-photon imaging localized candidate microglia–ensemble appositions, near-infrared branding enabled relocation, and the registered regions were processed for serial scanning electron microscopy (SEM) (Figure S3A). In both fear and extinction conditions, aligned EM micrographs revealed closely juxtaposed plasma membranes between microglial processes and ensemble bodies or neurites, with elongated non-synaptic interfaces conforming to neuronal contours (Figures S3B and S3C). These ultrastructural data validate the structural basis of the interface measurements.

We then resolved microglia–ensemble interactions at synaptic compartments using Leica Lightning super-resolution confocal imaging (effective lateral resolution ∼120 nm) to analyze tdTomato^+^ dendrites and spines together with GFP^+^ microglial processes (Figure 1I). Compared to CS-only controls, both fear and extinction conditions showed increased tdTomato^+^ spine density (Figure 1J, left), consistent with structural remodeling within the tagged ensembles. Microglial engagement, however, diverged sharply by memory type: fear retrieval reduced the number of microglial processes associated with tdTomato^+^ dendrites without changing total process–spine contact area, whereas extinction retrieval increased both the number of microglial processes engaging tdTomato^+^ dendrites and the total contact area at tdTomato^+^ spines (Figure 1J, middle and right). Categorizing tdTomato^+^ spines by the extent of microglial coverage (no contact, contact < 50% surface coverage, enwrapped ≥ 50%, engulfed) revealed no redistribution in fear memory, whereas extinction memory selectively shifted spines from “no contact” into “contact,” with enwrapped and engulfed fractions largely unchanged (Figures 1K and 1L). Thus, extinction retrieval strengthens microglial apposition at extinction-ensemble synaptic compartments, without a global increase in full spine envelopment or overt engulfment at this stage.

We next asked whether these extinction-associated microglial process states and interfaces are evident in the intact brain and can be followed longitudinally. To this end, we performed *in vivo* two-photon imaging in mPFC through a micro-prism.^43,44^ In CX3CR1-GFP mice, we repeatedly imaged the same microglia across baseline, re-extinction training, and extinction retrieval (Figures S4A and S4B). Here, day 2 (re-extinction) served as a renewed extinction-learning session after spontaneous recovery, whereas day 3 tested retrieval of the updated extinction trace. Across this time course, microglial moving area (a proxy for gross process motility) remained stable, whereas surveillance territory increased selectively at extinction retrieval and bulbous endings were elevated during both re-extinction and retrieval relative to baseline (Figures S4C–S4E). We then combined this longitudinal imaging design with TRAP labeling to directly quantify microglia–ensemble coupling *in vivo* (Figures 2A and 2B). In FosTRAP2::Ai14;;CX3CR1-GFP mice with extinction ensembles labeled before imaging, microglial soma area again remained stable, whereas process architecture progressively shifted toward a highly ramified configuration and peaked at extinction retrieval (Figures 2C–2G). Critically, microglial coverage of tdTomato^+^ neurites increased during re-extinction and further at retrieval (Figure 2H), demonstrating that extinction retrieval *in vivo* tightens microglia–ensemble engagement at neurite compartments in the same fields where microglial process expansion is observed.

**Figure 2.**
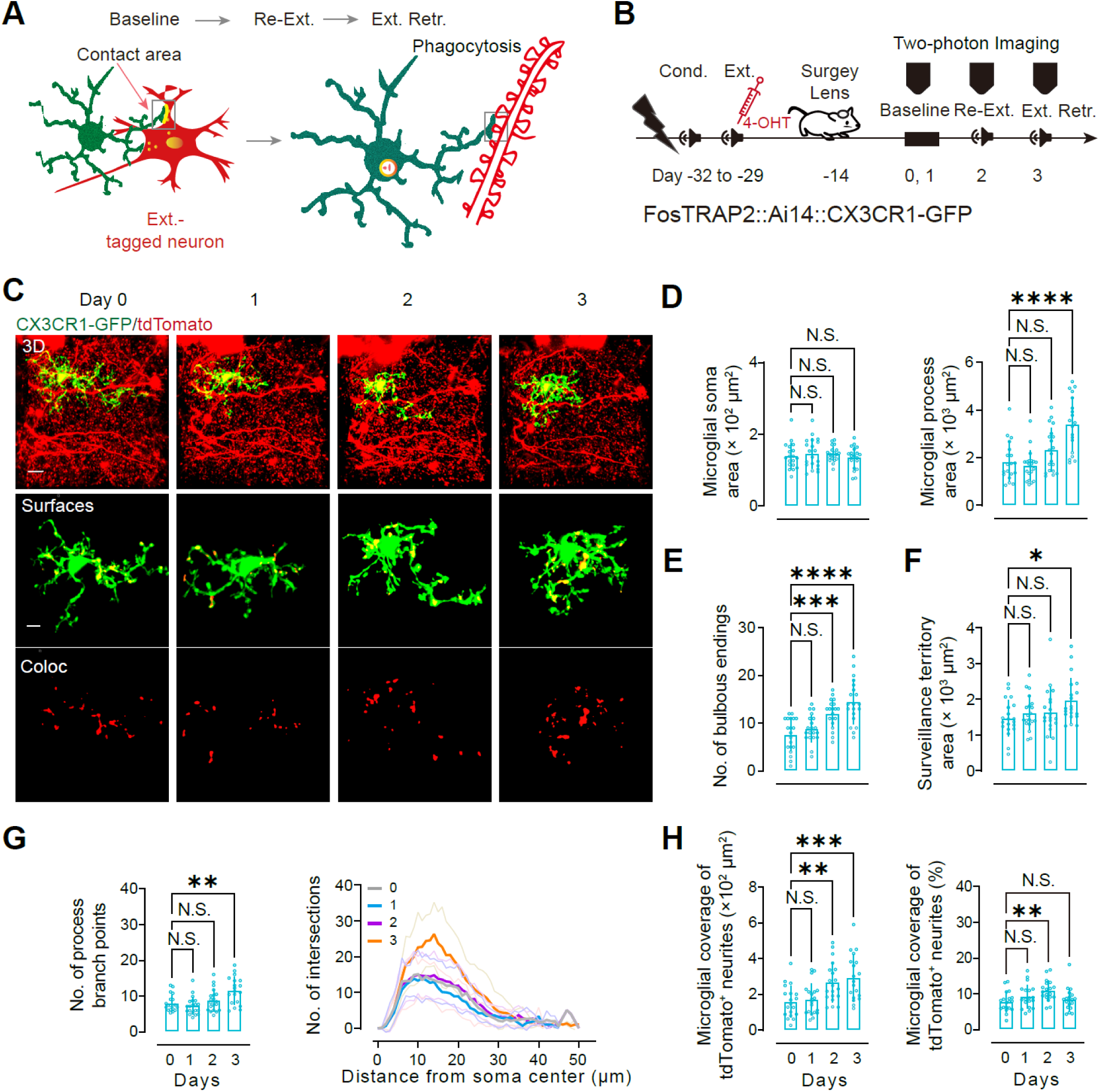
Longitudinal *in vivo* imaging reveals progressive microglial process remodeling and increased coupling to extinction ensembles. (A) Conceptual schematic illustrating that, across baseline, re-extinction (Re-Ext.), and extinction retrieval (Ext. Retr.), microglial processes can expand their contact interfaces with extinction-tagged neurites and engage phagocytic programs. (B) Experimental timeline for longitudinal two-photon imaging through a GRIN lens in FosTRAP2::Ai14;;CX3CR1-GFP mice. Mice underwent fear conditioning (Cond.) followed extinction training (Ext.); 4-OHT was administered after the third extinction session to TRAP-label extinction ensembles (tdTomato^+^). After GRIN-lens implantation over mPFC and recovery (∼4 weeks; allowing robust tdTomato expression and spontaneous recovery of extinguished fear), the same fields were imaged at baseline (Days 0–1), during re-extinction (Day 2), and at extinction retrieval (Day 3). (C) Representative z-projections showing CX3CR1-GFP^+^ microglia (green) and tdTomato^+^ extinction-ensemble neurites (red) across days (top row). Middle row, 3D surface renderings of the same microglia. Bottom row, voxels assigned to the microglia–tdTomato^+^ neurite interface (Coloc). Scale bar, 5 μm. (D to G) Longitudinal quantification of microglial morphology from the tracked cells, soma area remained stable (D, left), whereas process area (D, right), the number of bulbous endings (E), surveillance territory (F), and branching complexity (G; branch points and Sholl intersections) increased across sessions and peaked at extinction retrieval. (H) Microglial engagement of extinction ensembles quantified as microglial coverage of tdTomato^+^ neurites (left, absolute covered area; right, percent coverage). Coverage increased during re-extinction and was further elevated at extinction retrieval. *n* = 19 microglia (2 mice). Data are mean ± s.e.m. N.S., not significant; \**P* < 0.05, \*\**P* < 0.01, \*\*\**P* < 0.001, \*\*\*\**P* < 0.0001 (one-way ANOVA with Dunnett’s multiple-comparisons test versus Day 0).

Together, these results identify extinction retrieval as a state that reconfigures microglial process architecture and strengthens microglia–ensemble interfaces across neuronal bodies, neurites, and spines. In contrast, fear retrieval brings microglia into closer proximity to fear-tagged ensembles but does not comparably reinforce—and can reduce—microglial engagement at dendritic and synaptic compartments, strongly suggesting that microglia preferentially modulate extinction ensembles.

### Microglia preferentially engulf excitatory postsynaptic material from extinction ensembles

Having found that extinction retrieval selectively strengthens microglial interfaces with extinction-tagged somata, neurites, and spines and enriches distal bulbous endings, we next asked whether this engaged microglial state is accompanied by increased engulfment of ensemble-derived material. As an independent test that did not rely on a microglial reporter, we first analyzed TRAP2::Ai14 tissue in which microglia were identified by IBA1 immunostaining and ensemble-derived material by tdTomato. Microglial density in mPFC was comparable between fear and extinction conditions and their matched CS-only controls (Figures S5A and S5B), indicating that differential uptake could not be explained by differences in microglial abundance. Microglial area was unchanged during fear retrieval but increased selectively during extinction retrieval (Figure S5B), consistent with the extinction-associated expansion of microglial arbors described above. Importantly, 3D distance measurements revealed tdTomato^+^ puncta positioned within the microglial soma volume, as indicated by negative shortest distances relative to the microglial surface (Figure S5C, left), supporting bona fide internalization rather than perisomatic overlap. Quantification of internal tdTomato^+^ material in microglial somata uncovered opposite memory-type signatures: internalized tdTomato^+^ puncta decreased after fear retrieval but increased strongly after extinction retrieval relative to the corresponding CS-only controls (Figure S5C, middle and right).

We next defined this cargo accumulates and whether it is linked to a lysosomal engulfment program. In FosTRAP2::Ai14;;CX3CR1-GFP sections, we quantified CD68 area as a phagolysosomal burden index and measured internalized tdTomato^+^ puncta in microglial somata and processes (Figures 3A and 3B). Extinction retrieval increased CD68 burden (absolute and fractional area) relative to extinction CS-only controls, whereas fear retrieval did not (Figure 3C). In parallel, extinction retrieval elevated the total number of tdTomato⁺ puncta within microglia—including puncta in somata and in processes—whereas fear retrieval produced little change (Figure 3D). Restricting analysis to CD68^+^ microglia sharpened this contrast: tdTomato^+^ puncta decreased after fear retrieval but increased after extinction retrieval (Figure 3E), indicating selective phagocytic loading during extinction.

**Figure 3.**
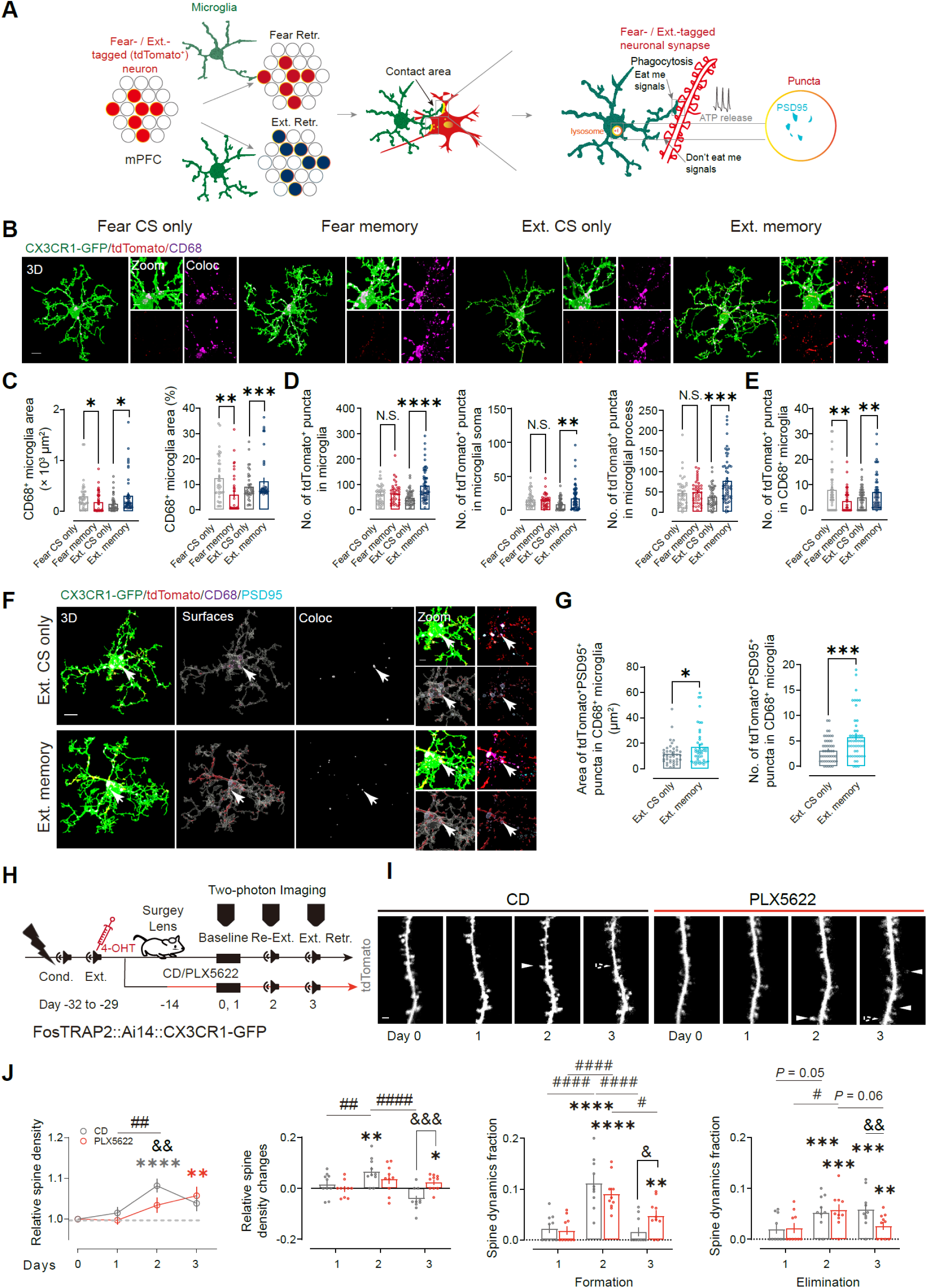
Extinction retrieval loads microglial lysosomes with excitatory postsynaptic cargo and limits stabilization of extinction-ensemble spines *in vivo*. (A) Analysis framework. Microglia contact fear- or extinction-tagged (tdTomato^+^) neuronal ensembles in mPFC and internalize ensemble-derived puncta into CD68^+^ phagolysosomes; engulfed excitatory postsynaptic cargo was identified by PSD-95. (B) Representative 3D confocal reconstructions from mPFC of FosTRAP2::Ai14;;CX3CR1-GFP mice showing CX3CR1-GFP^+^ microglia (green), TRAP-labeled ensemble material (tdTomato, red), and CD68 (magenta) in Fear CS-only, Fear memory, Ext. CS-only, and Ext. memory groups. Insets show higher-magnification views and colocalization masks (Coloc) used to score internalized tdTomato^+^ puncta. Scale bar, 10 μm. (C) CD68^+^ compartment size per microglia, expressed as absolute area (left) and as fraction of microglial area (right). Fear retrieval reduced CD68 burden relative to Fear CS-only controls, whereas extinction retrieval increased CD68 burden relative to Ext. CS-only controls. Fear CS only, *n* = 36 cells (7 mice); Fear memory, *n* = 38 cells (7 mice); Extinction CS only, *n* = 38 cells (7 mice); Extinction memory, *n* = 46 cells (7 mice). (D) Ensemble-derived cargo quantified as tdTomato^+^ puncta enclosed within microglial surfaces, shown as total puncta per microglia (left) and partitioned to soma (middle) or processes (right). Extinction retrieval increased tdTomato^+^ puncta in both somatic and process compartments relative to Ext. CS-only, whereas fear retrieval did not increase tdTomato^+^ puncta relative to Fear CS-only. Fear CS only, *n* = 40 cells (7 mice); Fear memory, *n* = 40 cells (7 mice); Extinction CS only, *n* = 46 cells (7 mice); Extinction memory, *n* = 59 cells (7 mice). (E) tdTomato^+^ puncta within CD68^+^ microglial compartments. tdTomato^+^ cargo within CD68⁺ volumes decreased after fear retrieval and increased after extinction retrieval relative to matched CS-only controls. Fear CS only, *n* = 40 cells (7 mice); Fear memory, *n* = 40 cells (7 mice); Extinction CS only, *n* = 38 cells (7 mice); Extinction memory, *n* = 49 cells (7 mice). (F) Representative reconstructions showing tdTomato^+^PSD-95^+^ puncta (cyan) within CD68^+^ microglial compartments (magenta) in Ext. CS-only (top) and Ext. memory (bottom). Arrows indicate tdTomato^+^PSD-95^+^ puncta contained within CD68^+^ volumes. Scale bars, 10 μm (main) and 2 μm (zoom). (G) Quantification of tdTomato^+^PSD-95^+^ puncta within CD68^+^ microglia in Ext. CS-only versus Ext. memory groups: summed puncta area per microglia (left) and puncta number per microglia (right). Extinction retrieval increased both measures, consistent with preferential uptake of extinction-ensemble excitatory postsynaptic cargo. Ext. CS only, *n* = 34 cells (5 mice); Ext. memory, *n* = 49 cells (5 mice). (H) Timeline for longitudinal two-photon imaging of extinction-ensemble spines in FosTRAP2::Ai14;;CX3CR1-GFP mice. Mice underwent conditioning (Cond.) and extinction training (Ext.); extinction ensembles were labeled by 4-hydroxytamoxifen (4-OHT) after the final extinction session. After surgery, the same dendritic fields were imaged at baseline (Days 0–1), during re-extinction (Re-Ext., Day 2), and at extinction retrieval (Ext. Retr., Day 3) in mice maintained on control diet (CD) or PLX5622 chow to deplete microglia. (I) Representative longitudinal images of tdTomato-labeled extinction-ensemble dendritic segments across Days 0–3 in CD and PLX5622 groups. Filled arrowheads indicate newly formed spines; open arrowheads indicate eliminated spines. Scale bar, 2 μm. (J) Spine density and turnover on extinction-ensemble dendrites with or without microglia. Relative spine density was normalized to Day 0; daily changes, formation fraction, and elimination fraction were quantified across imaging intervals. During re-extinction (Day 2), spine density increased in CD mice and showed a similar upward trend in PLX5622-treated mice. At extinction retrieval (Day 3), spine density declined in CD controls due to increased elimination with little additional formation, whereas it remained elevated in PLX5622-treated mice because formation persisted and elimination was reduced. CD, *n* = 10 dendritic segments (2 mice); PLX5622, *n* = 10 dendritic segments (3 mice). Data are mean ± s.e.m. N.S., not significant. \**P*< 0.05, \*\**P*< 0.01, \*\*\**P*< 0.001, \*\*\*\**P*< 0.0001, two-tailed unpaired Student’s *t* tests comparing each memory group with its matched CS-only control (C–E) or Ext. memory versus Ext. CS-only (G). Statistical significance is indicated in the panels (tests as described in Methods); in (J), * indicates comparisons versus Day 0 within a diet group, ^#^ indicates within-group comparisons across days, and ^&^ indicates CD versus PLX5622 at the same time point.

We then asked what synaptic material is preferentially targeted. Triple labeling of microglia (CX3CR1-GFP), ensembles (tdTomato), and synaptic markers showed that extinction retrieval selectively increased microglial contact with tdTomato^+^PSD-95^+^ puncta, whereas contact with tdTomato^+^vGLUT1^+^ and tdTomato^+^Gephyrin^+^ puncta was unchanged and contact with tdTomato^+^vGAT^+^ puncta decreased (Figure S6). Within microglia, engulfed tdTomato^+^PSD-95^+^ puncta increased, whereas tdTomato^+^vGLUT1^+^ and tdTomato^+^Gephyrin^+^ puncta unchanged and tdTomato^+^vGAT^+^ puncta declined (Figure S6). Consistent with this selectivity, CD68^+^ lysosomes in microglia contained more tdTomato^+^PSD-95^+^ puncta, and these puncta occupied a greater total area, after extinction retrieval than in extinction CS-only controls (Figures 3F and 3G). Thus, extinction retrieval preferentially drives microglial engulfment of excitatory postsynaptic material from extinction ensembles.

To test whether attenuating this engulfment program facilitates extinction, we treated TRAP2::Ai14;;CX3CR1-GFP mice with minocycline to suppress microglial activation.^23,45^ Minocycline accelerated extinction and reduced freezing at extinction retrieval without affecting acquisition (Figures S7A and S7B). Whole-cell recordings after extinction retrieval revealed an increased NMDAR/AMPAR ratio in extinction-tagged neurons (Figure S7C), consistent with a shift in excitatory postsynaptic signaling that paralleled the reduced microglial uptake of tdTomato^+^PSD-95^+^ puncta. In parallel, minocycline reduced microglial coverage of extinction-tagged neuronal bodies and neurites and decreased the number of contacting processes (Figures S7D–S7F), while lowering microglial process area and CD68 burden without changing soma size (Figures S7G–S7I). Concordantly, fewer tdTomato^+^ puncta—including tdTomato^+^PSD-95^+^ puncta—were detected within CD68^+^ microglial lysosomes, and their total area was reduced (Figures S7H and S7O). Together, these results show that reducing microglial engulfment of extinction-ensemble excitatory postsynaptic material can effectively enhances fear extinction.

### Microglial synapse editing limits extinction-associated spine remodeling *in vivo*

If microglia actively prune dendritic spines within extinction ensembles, then their heightened engagement during extinction retrieval should be reflected in spine remodeling dynamics *in vivo*. To test this prediction, we combined FosTRAP labeling of extinction ensembles with longitudinal *in vivo* two-photon imaging of the same tdTomato^+^ dendritic segments in mPFC through an implanted micro-prism (Figures 3H and 3I). Extinction ensembles were labeled by 4-OHT induction before imaging (Figure 3H). To determine whether microglia are required for the retrieval-linked spine dynamics observed in these ensembles, mice were maintained on either control diet (CD) or a diet containing the brain-penetrant CSF1R inhibitor PLX5622, which depletes microglia.^46-52^ Across the same imaging volumes, we quantified spine density and turnover on tdTomato^+^ extinction-ensemble dendrites during baseline (Day 0→1), re-extinction (Day 2), and extinction retrieval (Day 3) (Figure 3J). Baseline turnover was comparable between groups. During re-extinction (Day 2), which models renewed extinction learning after spontaneous recovery, spine density increased in CD mice and showed a similar upward trend in PLX5622-treated mice, consistent with net spine formation during this relearning phase (Figure 3J). By extinction retrieval (Day 3), which tests expression of the updated extinction trace, the trajectories diverged. In CD mice, spine density fell from the Day 2 peak, coincident with increased spine elimination and little additional formation (Figure 3J). In PLX5622-treated mice, by contrast, spine density continued to show an upward trend from re-extinction to retrieval (*P* = 0.06), driven by persistently elevated formation and reduced elimination relative to controls; accordingly, the retrieval-associated change in spine density (Day 3 vs Day 2) was greater in PLX5622-treated mice than in CD mice (Figure 3J). Thus, these live-imaging data indicate that microglia normally bias extinction ensembles toward spine loss at the time the updated extinction memory is expressed, while limiting further spine addition. Removing microglia shifts the same ensembles toward spine gain and retention through the retrieval phase.

### Disrupting microglial function accelerates extinction and elevates retrieval-linked activation of tagged ensembles

We next asked whether constraining microglial function is sufficient to improve fear extinction and alter retrieval-linked activation of tagged neuronal ensembles. Pharmacological depletion with PLX5622 beginning two weeks before conditioning did not affect fear acquisition, yet accelerated extinction learning and improved extinction retrieval (Figures 4A and 4B). This facilitation was reproduced when depletion was initiated after conditioning and restricted to the extinction phase (Figures S8A–S8H), and it disappeared after PLX5622 withdrawal (Figures S8I–S8P), which allowed microglial repopulation.^51,52^ To confirm that this effect did not depend on CSF1R inhibition alone, we used CX3CR1-CreER::ROSA26iDTR (CX3CR1^wt/CreER^::R26-CAG-LSL-tdTomato-2A-DTR) mice, in which tamoxifen-inducible Cre enables temporally controlled ablation of adult CX3CR1^+^ CNS-resident myeloid cells, including parenchymal microglia.^53,54^ This independent depletion strategy produced the same behavioral effect (Figure S9), supporting the conclusion that microglia act as a brake on extinction.

**Figure 4.**
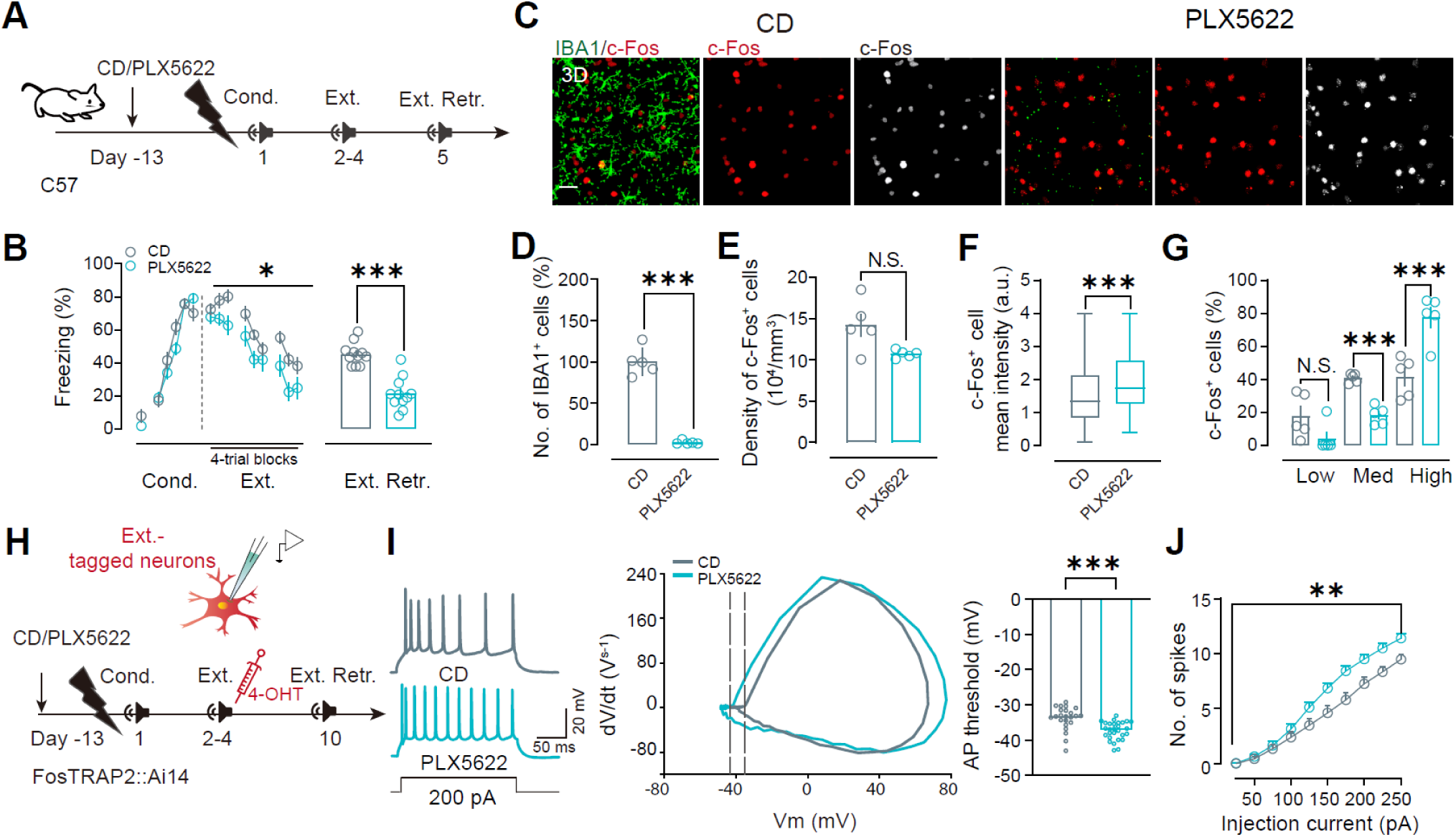
Microglial depletion elevates extinction-retrieval–locked neuronal activation and increases the excitability of extinction-tagged mPFC neurons. (A) Treatment schedule in C57BL/6J mice fed CD or PLX5622 beginning before conditioning. (B) Freezing across conditioning, extinction training, and extinction retrieval in CD and PLX5622-treated mice. CD, *n* = 11 mice; PLX5622, *n* = 11 mice. (C) Representative mPFC sections immunolabeled for microglia (IBA1, green) and neuronal activity (c-Fos, red) from CD and PLX5622 groups. Rightmost panels show segmented c-Fos signal used for 3D quantification. Scale bar, 50 μm. (D to G) Quantification of depletion efficacy and retrieval-evoked activation. (D) IBA1^+^ microglia abundance (normalized to CD) confirms robust depletion by PLX5622. (D) Density of c-Fos^+^ cells is unchanged. (E) Mean c-Fos fluorescence intensity is increased by PLX5622. (F) Distribution of c-Fos^+^ cells across intensity bins (Low, 500–1500 a.u.; Med, 1500–2500 a.u.; High, ≥2500 a.u.) shows a shift toward the Fos-high population with PLX5622. CD, *n* = 8099 cells; PLX5622, *n* = 7398 cells; pooled from 5 mice per group. (G) Summary of the fraction of c-Fos+ cells in each intensity bin. (H) Timeline for intrinsic excitability recordings from extinction-tagged neurons. FosTRAP2::Ai14 mice were fed CD or PLX5622 (day −13), conditioned (day 1), extinguished (days 2–4; 4-OHT after the third extinction session to label extinction ensembles), and tested at extinction retrieval (day 10); whole-cell recordings were performed in acute mPFC slices from tdTomato^+^ neurons. (I and J) PLX5622 increases the intrinsic excitability of extinction-tagged neurons. (I) Example spike trains evoked by a 200-pA current step (left), representative phase plots (middle), and action potential (AP) threshold summary (right). (J) Frequency–current relationship showing increased spike output with PLX5622. CD, n = 23 neurons (9 mice); PLX5622, n = 28 neurons (11 mice). Data are mean ± s.e.m. N.S., not significant; \**P* < 0.05, \*\**P* < 0.01, \*\*\**P* < 0.001, unpaired Student’s *t* test (between groups) or two-way repeated-measures ANOVA with post hoc tests (as appropriate).

We then tested how microglia influence activation of tagged ensembles at retrieval. Using the same TRAP strategy described above, we identified tdTomato^+^c-Fos^+^ neurons, that is, neurons active both during the original TRAP-labeling window and during the retrieval test (Figure S10). During fear retrieval, the density of tdTomato^+^ neurons was unchanged relative to Fear CS-only controls, but the distribution of tdTomato^+^c-Fos^+^ neurons shifted toward medium- and high-intensity c-Fos labeling, with a corresponding reduction in the low-intensity fraction (Figures S10D–S10H). During extinction retrieval, both tdTomato⁺ and c-Fos⁺ neuronal densities increased, whereas the fraction of tdTomato^+^c-Fos^+^ neurons remained comparable to Extinction CS-only controls; importantly, these double-positive neurons again shifted toward higher c-Fos intensity categories (Figures S10K–S10O). Thus, in both memory conditions, retrieval preferentially engages the more strongly activated subset of the tagged ensemble.

Consistent with this framework, microglial depletion amplified retrieval-linked neuronal activation. At extinction retrieval, depletion—whether instituted before conditioning or restricted to the extinction phase—increased c-Fos intensity per labeled neuron and increased the proportion of Fos-high neurons (Figures 4C–4G, and S8C–S8H); both measures returned toward baseline after microglial repopulation (Figures S8K–S8P). During fear retrieval, depletion similarly elevated c-Fos intensity and increased the Fos-high fraction despite unchanged freezing behavior (Figures S11A–S11H). These observations indicate that microglia normally constrain the upper range of retrieval-time neuronal activation.

To determine whether this activity shift is accompanied by altered intrinsic excitability, we performed whole-cell current-clamp recordings from tdTomato^+^ neurons in acute mPFC slices prepared after retrieval. In extinction-tagged neurons, microglial depletion increased action-potential output across depolarizing current steps after extinction retrieval (Figures 4H–4J). In contrast, fear-tagged neurons recorded after fear retrieval did not show a comparable increase in spike output (Figures S11I–S11K). These data indicate that microglia preferentially restrain the excitability of extinction-tagged ensembles at the time extinction is expressed.

We next asked whether elevating activity in fear-tagged neurons is sufficient to bias subsequent extinction. Using TRAP-driven hM3Dq, we chemogenetically increased excitability in fear ensembles and administered clozapine *N*-oxide (CNO) immediately after each extinction session (Figure S12). This manipulation accelerated extinction learning and reduced freezing at extinction retrieval, indicating that increased activity in fear-tagged neurons is sufficient to promote subsequent extinction.

Together, these convergent behavioral, activity-mapping, electrophysiological, and chemogenetic data indicate that microglia hinder fear extinction by limiting retrieval-linked activation of tagged ensembles, with a particularly strong constraint on extinction-ensemble excitability during extinction retrieval.

### Fear-ensemble purinergic signaling promotes microglial process engagement through P2Y12

We next asked what activity-linked cues recruit microglial processes to specific ensemble compartments during retrieval. Because P2Y12 directs microglial process responses to extracellular nucleotides, and K_V_2.1-enriched somatic junctions have been implicated as sites of neuron–microglia communication,^20^ we examined whether fear and extinction ensembles present distinct purinergic and junction-associated microdomains at microglia-contacted surfaces. To do this, we immunolabeled vNUT and K_V_2.1 in TRAP2::Ai14;;CX3CR1-GFP mice and quantified, on tdTomato^+^ neuronal bodies, the marker-positive surface apposed by microglia using 3D reconstructions (Figure 5A).

**Figure 5.**
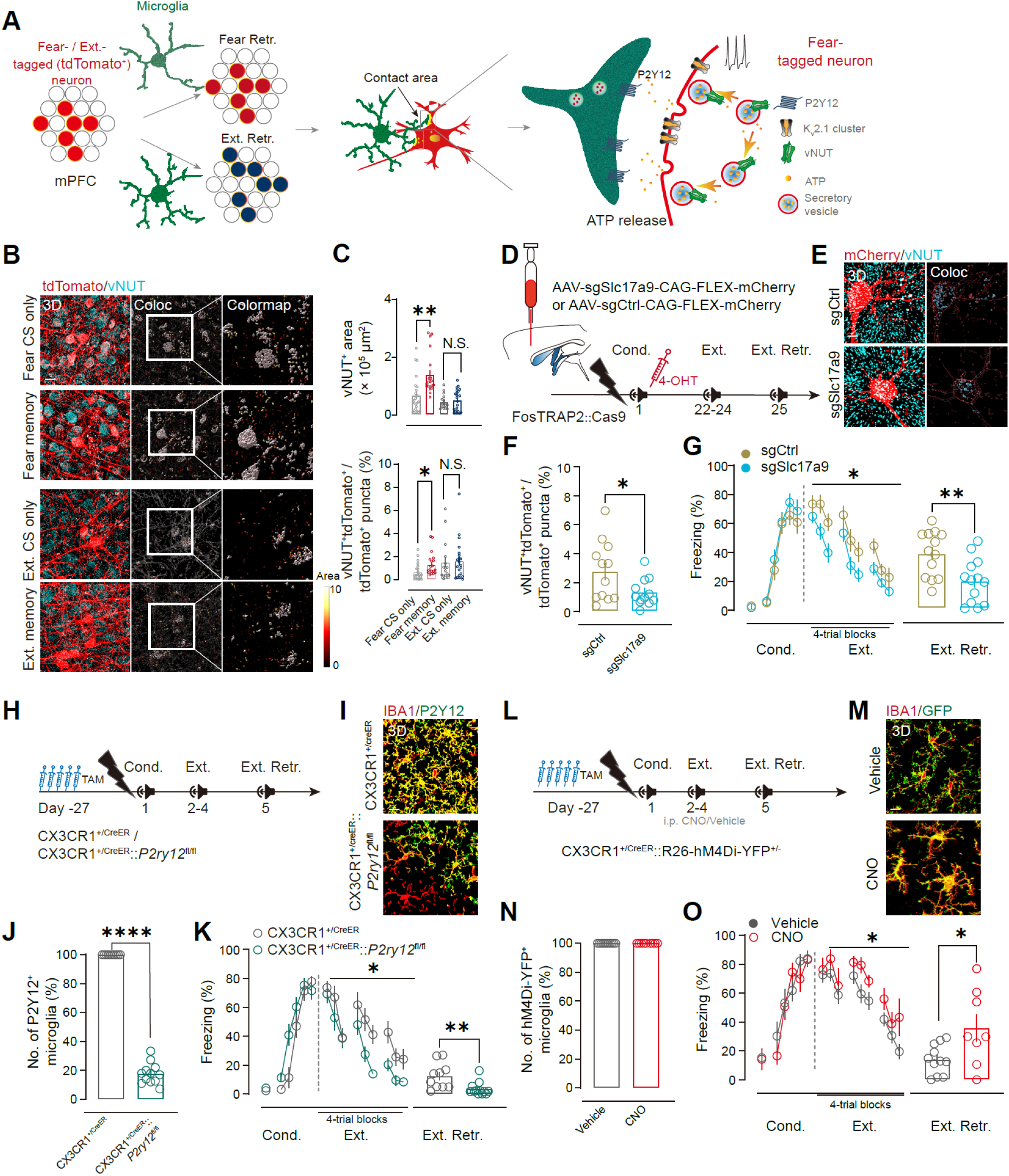
Ensemble-derived ATP signals through microglial P2Y12–Gi to restrain fear extinction. (A) Working model. During retrieval, fear-ensemble neurons enrich the vesicular nucleotide transporter vNUT (encoded by *Slc17a9*) at somatic/neuritic microdomains, promoting vesicular ATP release that is sensed by P2Y12 receptors on nearby microglial processes to shape microglial engagement and phagocytic output at ensemble compartments. (B) Representative 3D confocal reconstructions showing vNUT immunoreactivity (cyan) in tdTomato^+^ ensemble neurons (red) in mPFC after fear retrieval (Fear memory) or extinction retrieval (Ext. memory), or their respective CS-only controls. Surfaces, 3D surface rendering; Coloc, vNUT signal colocalized with TRAP labeling (pseudocolor). Scale bar, 10 μm. (C) Quantification of vNUT enrichment in tdTomato^+^ neurons. Top, vNUT^+^ area associated with tdTomato^+^ labeling. Bottom, fraction of tdTomato^+^ puncta that are vNUT^+^. Fear retrieval increased vNUT in fear ensembles, whereas extinction retrieval did not measurably change vNUT in extinction ensembles. Fear CS only, *n* = 23 sections (6 mice); Fear memory, *n* = 20 sections (6 mice); Ext. CS only, *n* = 16 sections (6 mice); Ext. memory, *n* = 22 sections (6 mice). (D) Ensemble-restricted disruption of vesicular ATP loading. FosTRAP2::Cas9 mice received intra-mPFC microinjection of AAV encoding sgSlc17a9 (or sgCtrl) in conjunction with mCherry in a Cre-dependent (FLEX) cassette. 4-OHT was administered after conditioning to restrict Cas9 expression/edition to fear ensemble neurons; extinction training and extinction retrieval were performed subsequently. (E) Representative images illustrating reduced vNUT signal in mCherry-labeled fear ensembles after sgSlc17a9 compared with SgCtrl. Scale bar, 7 μm. (F) Editing/knockdown efficiency, quantified as the vNUT^+^tdTomato^+^/tdTomato^+^ puncta ratio in mCherry^+^ neurons. sgCtrl, *n* = 12 sections; sgSlc17a9, *n* = 11 sections. (G) Behavioral consequence of fear-ensemble *Slc17a9* disruption. sgSlc17a9 accelerated extinction learning and reduced freezing at extinction retrieval relative to sgCtrl. sgCtrl, *n* = 13 mice; sgSlc17a9, *n* = 13 mice. (H) Timeline for tamoxifen-induced microglia-specific deletion of *P2ry12* (CX3CR1^+/CreER^::*P2ry12^fl/fl^*). (I) Representative mPFC images showing IBA1 (red) and P2Y12 (green) immunostaining in controls and microglial *P2ry12* conditional knockouts. Scale bar, 30 μm. (J) Fraction of P2Y12^+^ microglia (among IBA1^+^ cells), confirming efficient deletion. *n* = 10 mice (control) or 12 mice (cKO), one section per mouse. (K) Behavioral effect of microglial *P2ry12* deletion. *P2ry12* loss facilitated extinction and reduced freezing at extinction retrieval without disrupting acquisition. CX3CR1^+/CreER^, *n* = 10 mice; CX3CR1^+/CreER^::*P2ry12^fl/fl^*, *n* = 12 mice. (L) Chemogenetic engagement of microglial Gi signaling. CX3CR1^+/CreER^::*R26-^hM4Di-YFP/+^*mice received tamoxifen before conditioning; CNO or vehicle was administered intraperitoneally during extinction training. (M) Representative images showing IBA1 (red) and hM4Di-YFP (green) in mPFC after vehicle or CNO. Scale bar, 10 μm. (N) Fraction of microglia expressing hM4Di-YFP, showing that CNO does not alter expression. Vehicle, *n* = 11 sections; CNO, *n* = 8 sections. (O) Behavioral effect of microglial Gi activation. CNO impaired extinction learning and increased freezing at extinction retrieval relative to vehicle. Vehicle, *n* = 11 mice; CNO, *n* = 8 mice. Data are mean ± s.e.m. N.S., not significant; \**P* < 0.05, \*\**P* < 0.01, \*\*\*\**P* < 0.0001, unpaired two-sided Student’s *t* test for two-group comparisons; two-way repeated-measures ANOVA for time courses with appropriate post hoc tests.

Fear retrieval selectively increased microglial apposition to vNUT^+^ microdomains on fear-ensemble surfaces, as reflected by increases in both covered area and fractional coverage relative to fear CS-only controls (Figures 5B, 5C, S13A, and S13B). Thus, fear retrieval did not globally strengthen the fear-ensemble interface, but instead redistributed microglial contact toward vNUT-rich microdomains. By contrast, extinction retrieval did not measurably alter vNUT microdomains at the extinction-ensemble interface (Figures 5B and 5C). K_V_2.1 showed the opposite, memory-type–specific pattern: extinction retrieval increased microglial apposition to K_V_2.1^+^ domains on extinction-ensemble bodies, whereas fear retrieval reduced K_V_2.1 engagement at fear-ensemble bodies (Figures S13C and S13D). These findings indicate that fear and extinction ensembles present distinct molecular signatures at microglia-contacted surfaces.

To test whether VNUT-dependent purinergic signaling from fear ensembles contributes causally to extinction suppression, we disrupted vesicular ATP loading selectively in fear ensembles by targeting *Slc17a9*, which encodes vNUT, in TRAP2::Cas9 mice (Figure S14), with 4-OHT delivered after conditioning to restrict editing to fear-tagged neurons (Figure 5D). vNUT was reduced in tdTomato^+^ neurons (Figures 5E and 5F), and its disruption accelerated extinction learning and reduced freezing at extinction retrieval (Figure 5G), indicating that vNUT-dependent signaling from fear ensembles restrains extinction. Because ATP-derived purinergic signaling activates microglial P2Y12 and downstream Gi signaling,^18,48,55,56^ we next tested this microglial pathway directly. Microglia-specific knockout of P2Y12 (which is encoded by *P2ry12*) facilitated extinction (Figures 5H–5K), and chronic intracerebroventricular infusion of the P2Y12 antagonist clopidogrel produced a similar behavioral effect (Figure S15A). Notably, clopidogrel did not measurably alter gross microglia–ensemble body apposition or the P2Y12-positive interface area (Figures S15B–S15F), but it reduced microglial process territory while preserving soma size, decreased CD68 burden, and lowered internalized tdTomato^+^ puncta—most prominently within microglial processes (Figures S15G–S15K). Conversely, chemogenetic activation of Gi signaling in microglia with hM4Di impaired extinction (Figures 5L–5O). Together, these results support a model in which vNUT-dependent purinergic signaling from fear ensembles engages microglial P2Y12 signaling to promote a process-rich, uptake-competent state that restrains extinction.

### Extinction and fear ensembles oppositely gate engulfment through PS exposure and CD47 protection

Because microglia preferentially removed excitatory postsynaptic material from extinction ensembles, we next asked what molecular cues mark one ensemble for engulfment while shielding the other. Microglial uptake of neuronal material is often initiated when phosphatidylserine (PS) becomes exposed on the outer leaflet of the plasma membrane and is bridged by Gas6 to the microglial phagocytic receptor MER tyrosine kinase (MERTK).^57-59^ To assess extracellularly exposed PS (ePS) *in vivo*, we delivered the membrane-impermeant PS-binding probe PSVue into the lateral ventricle^60^ and quantified PSVue signal in mPFC during retrieval in TRAP2::Ai14;;CX3CR1-GFP mice (Figures 6A–6C). Total PSVue⁺ area did not differ across memory groups and their matched CS-only controls (Figure 6D, left), arguing against a global increase in ePS during retrieval. Instead, extinction retrieval selectively increased the fraction of PSVue puncta associated with tdTomato^+^ extinction ensembles, whereas fear retrieval did not increase PSVue association with tdTomato^+^ fear ensembles (Figure 6D, right). Because PSVue binds externalized PS before internalization, PSVue signal detected inside microglia most directly reports uptake of PS-bearing material. By this measure, fear retrieval reduced PSVue^+^ puncta within microglia without changing ensemble-associated PSVue, whereas extinction retrieval increased ensemble-associated PSVue without increasing total PSVue^+^ puncta within microglia (Figure 6E). This pattern indicates that retrieval does not globally increase PS-tagged cargo uptake, but instead redistributes PS exposure toward extinction-ensemble compartments. retrieval does not globally increase PS-tagged cargo uptake, but instead redistributes PS exposure toward extinction-ensemble compartments. In parallel, Gas6 immunoreactivity decreased after fear retrieval but increased after extinction retrieval, while the fraction overlapping tdTomato labeling remained stable (Figures S16A and S16B), consistent with retrieval-dependent tuning of PS–Gas6 signaling.

**Figure 6.**
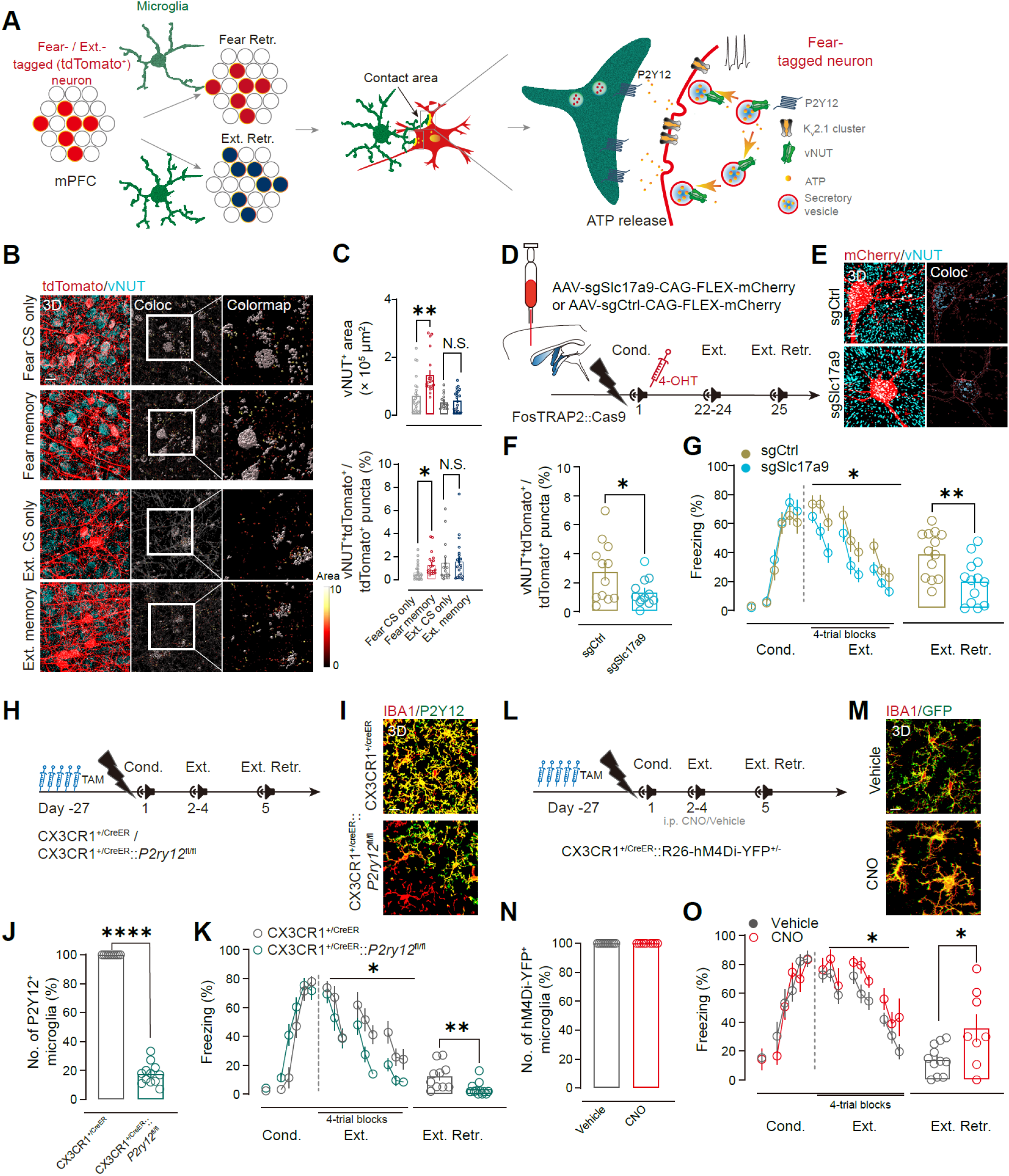
Extinction ensembles expose phosphatidylserine and engage microglial MERTK signaling to restrain extinction. (A) Schematic of an “eat-me” pathway in which extracellularly exposed phosphatidylserine (PS) on ensemble membranes is bridged by Gas6 to the microglial phagocytic receptor MERTK, promoting engulfment. (B) Timeline for labeling extracellular PS with PSVue-647 (a membrane-impermeant PS probe delivered intracerebroventricularly through an implanted cannula) and assessing fear or extinction retrieval in FosTRAP2::Ai14;;CX3CR1-GFP mice (ensemble tagging as in Figure 1). (C) Representative mPFC confocal images showing microglia (CX3CR1-GFP, green), TRAP-labeled ensemble neurons (tdTomato, red), and PSVue signal (cyan). Right, “Coloc” indicates PSVue puncta segmented within microglial surfaces (i.e., PSVue-labeled PS-bearing material internalized by microglia). Scale bar, 10 μm. (D) Total PSVue signal per field (left) and fraction of PSVue signal associated with TRAP structures (right). Fear CS only, *n* = 12 sections (4 mice); Fear memory, *n* = 22 sections (6 mice); Ext. CS only, *n* = 29 sections (7 mice); Ext. memory, *n* = 26 sections (7 mice). N.S., not significant; \*\*\**P* < 0.001, unpaired Student’s *t* test. (E) Quantification of PSVue^+^ puncta detected within microglia (left) and TRAP⁺PSVue⁺ puncta within microglia (right). Fear CS only, *n* = 8 sections (3 mice); Fear memory, *n* = 16 sections (6 mice); Ext. CS only, *n* = 29 sections (7 mice); Ext. memory, *n* = 26 sections (7 mice). (F) Ensemble-targeted CRISPR–Cas9 strategy to disrupt Tmem16f selectively in extinction-tagged neurons in FosTRAP2::Cas9 mice using AAV-sgTmem16f-CAG-FLEX-mCherry or AAV-sgCtrl-CAG-FLEX-mCherry, with 4-OHT administered after extinction training to restrict editing to extinction ensembles. (G) Representative mPFC images showing mCherry-labeled extinction ensemble neurons (red) and TMEM16F immunoreactivity (cyan); “Coloc” indicates TMEM16F signal within mCherry⁺ surfaces. Scale bar, 5 μm. (H) TMEM16F knockdown efficiency in tdTomato^+^ neurons (fraction of tdTomato^+^ puncta positive for TMEM16F). sgCtrl, *n* = 20 sections (5 mice); sgTmem16f, *n* = 20 sections (5 mice). (I) Freezing during conditioning, extinction, re-extinction, and extinction retrieval in sgCtrl and sgTmem16f groups. sgCtrl, *n* = 12 mice; sgTmem16f, *n* = 14 mice. (J) Scheme for tamoxifen-induced microglia-specific deletion of Mertk (CX3CR1^+/CreER^::*Mertk^fl/fl^*). (K) Representative mPFC images showing microglia (IBA1, red) and MERTK (green); “Coloc” indicates MERTK signal within IBA1^+^ microglia. Scale bar, 10 μm. (L) Fraction of IBA1^+^ microglia that are MERTK^+^. (M) Freezing during conditioning, extinction, and extinction retrieval in control and microglia-specific Mertk conditional knockout mice. CX3CR1^+/CreER^, *n* = 12 mice; and CX3CR1^+/CreER^::*Mertk^fl/fl^*, *n* = 11 mice. Data are mean ± s.e.m. N.S., not significant; \**P* < 0.05, \*\**P* < 0.01, \*\*\**P* < 0.001, \*\*\*\**P* < 0.0001, unpaired Student’s *t* test for two-group comparisons; two-way repeated-measures ANOVA for behavioral time courses, with post hoc tests as appropriate.

We next asked how extinction ensembles generate ePS. TMEM16F is a Ca^2+^-activated scramblase/ion channel that can promote PS externalization.^61^ TMEM16F abundance in mPFC was unchanged after fear or extinction retrieval (Figures S16C and S16D), suggesting that retrieval-linked PS exposure is more likely to reflect activity-dependent regulation than bulk changes in protein expression. To test whether TMEM16F within extinction ensembles contributes functionally to extinction restraint, we reduced *Tmem16f* selectively in extinction-tagged neurons using TRAP-targeted CRISPR–Cas9. This manipulation enhanced extinction performance (Figures 6F–6I and S17), supporting the idea that PS exposure from extinction ensembles contributes to microglia-dependent suppression of extinction.

On the microglial side, both fear and extinction retrieval increased MERTK immunoreactivity and the fraction of MERTK^+^ microglia (Figures S16E and S16F), indicating that retrieval generally places microglia in a state capable of reading PS-related cues. To test whether this receptor is required for the behavioral effect, we conditionally deleted *Mertk* in microglia using CX3CR1-CreER::MERTK^fl/fl^ mice. Microglial *Mertk* deletion improved fear extinction without altering acquisition (Figures 6J–6M), consistent with MERTK-dependent engulfment acting as a brake on extinction. By contrast, knockout of triggering receptor expressed on myeloid cells 2 (TREM2), another microglial receptor associated with phagocytic activation,^62^ did not alter extinction learning or retrieval (Figure S18), arguing that the effect is not a generic consequence of disrupting any phagocytosis-related receptor.

The preferential pruning of extinction, but not fear, ensembles further suggested that fear ensembles might be actively protected from microglial engulfment. We therefore examined the canonical “don’t-eat-me” signal in which neuronal CD47 engages microglial SIRPα to inhibit phagocytosis.^35,63^ Fear retrieval increased overall CD47 signal and elevated the fraction of CD47^+^ puncta colocalized with tdTomato^+^ fear-ensemble membranes, whereas extinction retrieval left total CD47 largely unchanged but reduced tdTomato-associated CD47 (Figure 7A and B). Similarly, SIRPα signal and SIRPα^+^ area per microglial cell increased after fear retrieval but were unchanged during extinction retrieval (Figures 7C and 7D). Functionally, disrupting CD47 selectively in fear ensembles increased tdTomato^+^ puncta within microglia and accelerated extinction (Figures 7E–7H and S19). Continuous intracerebroventricular infusion of a SIRPα-blocking antibody similarly increased microglial uptake of fear-ensemble material and facilitated extinction (Figures 7I–7M). Thus, fear retrieval is accompanied by active molecular protection of fear ensembles from microglial engulfment.

**Figure 7.**
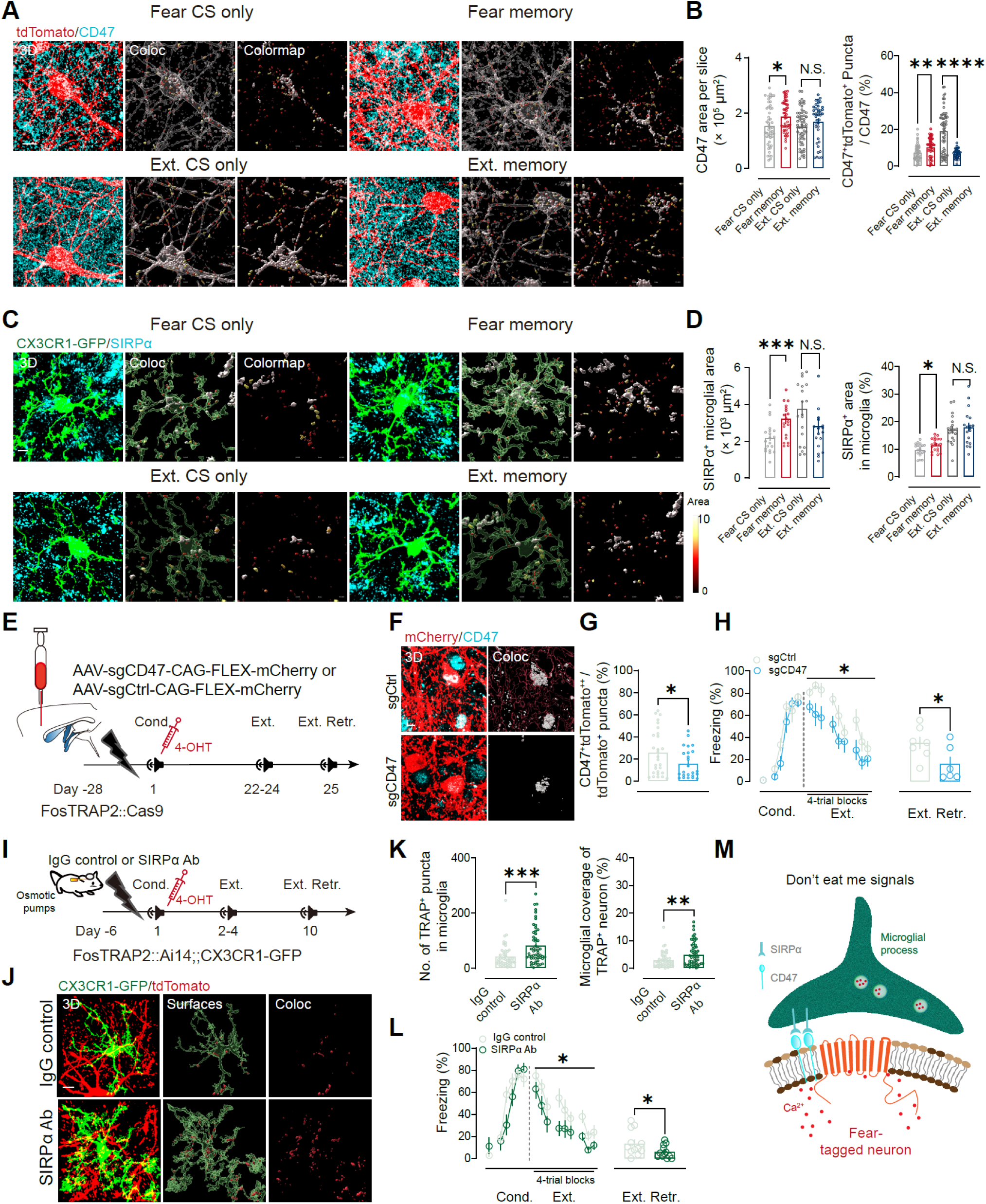
Fear ensembles deploy a CD47–SIRPα “don’t-eat-me” gate that limits microglial uptake and preserve fear memory. (A) Representative confocal images showing TRAP-labeled ensemble neurons (tdTomato, red) and CD47 immunoreactivity (cyan) in Fear CS-only, Fear memory (fear retrieval), Ext. CS-only, and Ext. memory (extinction retrieval) groups. Surfaces, 3D reconstructions of tdTomato^+^ neuronal membranes; Coloc, CD47 signal associated with TRAP⁺ membranes. Scale bar, 10 µm. (B) Quantification of CD47 signal on tdTomato^+^ membranes (left, CD47^+^ area per section) and CD47 enrichment on ensemble membranes (right; percentage of tdTomato^+^ puncta that are CD47^+^). Fear CS only, *n* = 50 sections (5 mice); fear memory, *n* = 50 sections (5 mice); Ext. CS only, *n* = 60 sections (5 mice); Ext. memory, *n* = 40 sections (5 mice). (C) Representative images of microglia (CX3CR1-GFP, green) and SIRPα (cyan) in the four behavioral groups. Surfaces, 3D reconstructions of individual microglia; Coloc, SIRPα signal within microglial volumes. Scale bar, 5 µm. (D) Quantification of microglial SIRPα (left; SIRPα^+^ area within microglia per section; right, SIRPα^+^ area normalized to microglial area). Fear CS only, *n* = 19 sections (5 mice); fear memory, *n* = 20 sections (mice); Ext. CS only, *n* = 20 sections (5 mice); Ext. memory, *n* = 18 sections (5 mice). (E) Strategy for fear-ensemble–restricted Cd47 disruption. FosTRAP2::Cas9 mice received intra-mPFC microinjection of AAV-sgCD47-CAG-FLEX-mCherry or AAV-sgCtrl-CAG-FLEX-mCherry; 4-OHT after conditioning restricted editing to fear-activated (tdTomato^+^) neurons. (F) Representative mPFC images after extinction retrieval showing reduced CD47 signal in mCherry-labeled fear ensembles in sgCd47 mice. Coloc, CD47 signal associated with tdTomato^+^ neuronal volumes. Scale bar, 10 µm. (G) Cd47 disruption efficiency, quantified as the percentage of tdTomato^+^ puncta that are CD47^+^. sgCtrl, *n* = 28 sections (7 mice); sgCd47, *n* = 24 sections (6 mice). \**P* < 0.05, unpaired Student’s *t* test. (H) Behavioral effect of fear-ensemble CD47 disruption: accelerated extinction training and reduced freezing at extinction retrieval. sgCtrl, *n* = 7 mice; sgCd47, *n* = 6 mice. (I) Timeline for continuous intra-mPFC microinjection of infusion of a SIRPα-blocking antibody or IgG control (osmotic pump) in FosTRAP2::Ai14;;CX3CR1-GFP mice across conditioning, extinction training, and extinction retrieval. (J) Representative images after extinction retrieval showing microglia (CX3CR1-GFP, green) and fear ensembles (tdTomato, red) under IgG control or anti-SIRPα treatment. *Surfaces*, microglial reconstructions; *Coloc*, tdTomato^+^ puncta detected within microglial volumes. Scale bar, 10 µm. (K) SIRPα blockade increased microglial engagement of fear ensembles (left, tdTomato^+^ puncta per microglia; right, percent microglial coverage of tdTomato^+^ neuronal bodies). IgG, *n* = 48 microglia (6 mice); anti-SIRPα, n = 55 microglia (6 mice). (L) Behavioral effect of SIRPα blockade: facilitated extinction and reduced freezing at extinction retrieval. IgG, *n* = 13 mice; anti-SIRPα, *n* = 13 mice. (M) Model: fear ensembles up-regulate CD47, engaging microglial SIRPα to inhibit phagocytosis (“don’t-eat-me” signaling). Data are mean ± s.e.m. N.S., not significant; \**P* < 0.05, \*\**P* < 0.01, \*\*\**P* < 0.001, \*\*\*\**P* < 0.0001, two-sided unpaired Student’s *t* test for single comparisons; two-way repeated-measures ANOVA with appropriate post hoc tests for learning curves, and unpaired Student’s *t* test for retrieval comparisons, as applicable.

Together, these results show that the microglia-facing surfaces of fear and extinction ensembles are specified in opposite ways during retrieval. Extinction ensembles become more permissive to microglial engulfment, whereas fear ensembles become more resistant, enabling microglial processes to selectively prune extinction-ensemble synapses and thereby preserve fear memory (Figure 8).

**Figure 8.**
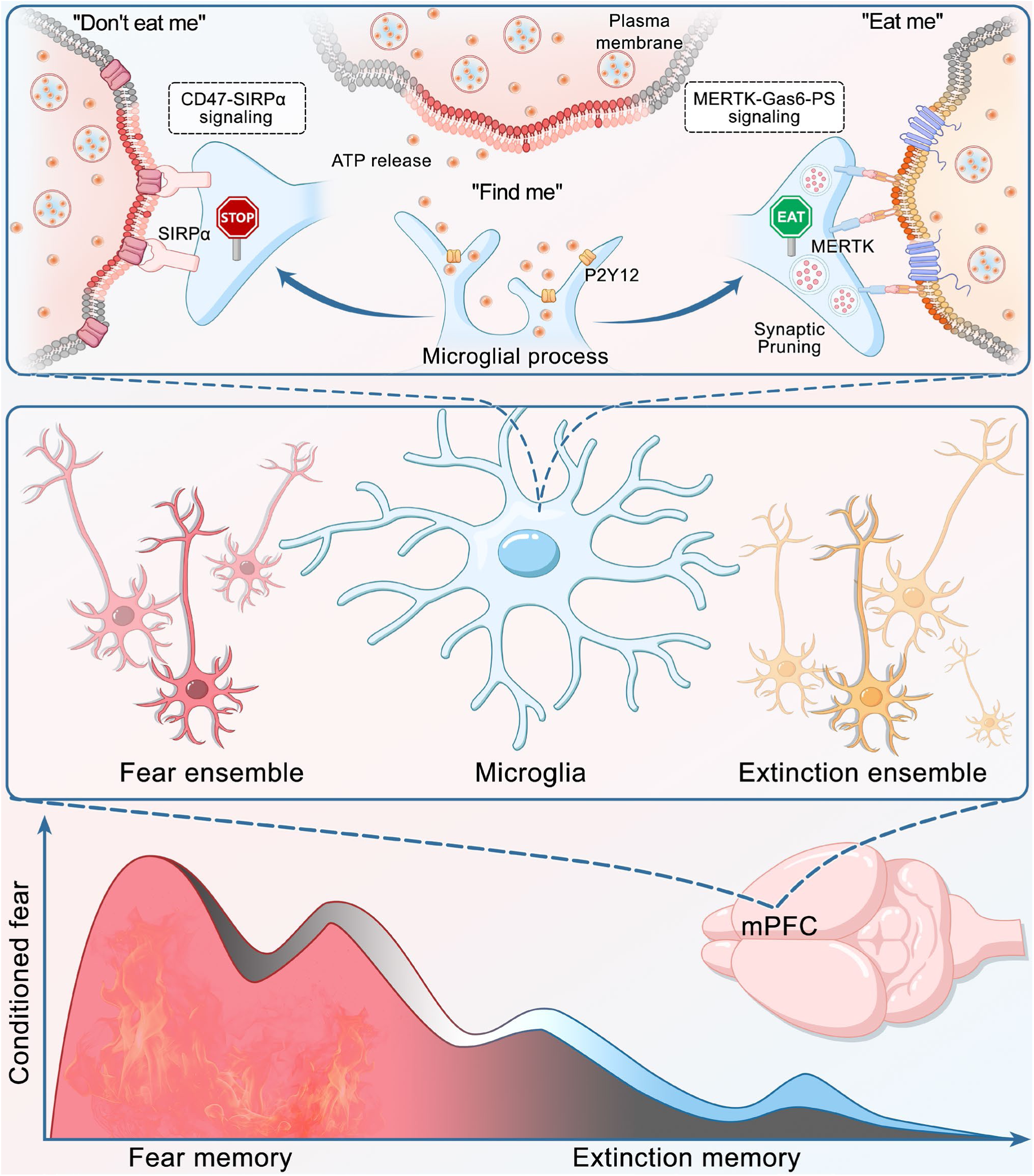
Working model for how microglial processes use local “find-me,” “eat-me,” and “don’t-eat-me” cues to preserve fear memory. Top, microglial processes survey neuronal membranes and integrate three classes of local signals that determine where they accumulate and whether synaptic material is removed. “Don’t-eat-me” signaling: fear-tagged neuronal surfaces are enriched for CD47, which engages microglial SIRPα and locally suppresses phagocytic output, thereby protecting fear-ensemble synapses from pruning. “Find-me” signaling: retrieval-linked neuronal activity promotes VNUT-dependent vesicular ATP release, which is sensed by P2Y12 receptors on nearby microglial processes and biases their recruitment toward active compartments. “Eat-me” signaling: at extinction-tagged synaptic compartments, activity-linked exposure of phosphatidylserine (ePS), together with Gas6-mediated engagement of microglial MERTK, licenses engulfment and lysosomal degradation of tagged postsynaptic material. Middle, conceptual organization in mPFC: microglia are positioned between fear and extinction ensembles and are differentially instructed by their local surface cues. Bottom, consequence for behavior: preferential pruning of extinction-tagged synapses, together with protection of fear-tagged synapses, biases the balance between competing memory traces toward persistence of conditioned fear and limits the decline in fear expression across extinction. Reweighting these cues toward weaker recruitment/engulfment or reduced protection is predicted to favor extinction. Abbreviations: mPFC, medial prefrontal cortex; vNUT, vesicular nucleotide transporter; ePS, externalized phosphatidylserine; MERTK, MER tyrosine kinase; SIRPα, signal regulatory protein alpha.

## Discussion

Our data position microglia as selective editors of adult memory circuits rather than passive responders to neuronal activity. In the mPFC, extinction retrieval shifts microglia into a hyper-ramified, non-amoeboid process state and strengthens process-mediated interfaces with extinction ensembles across neuronal body, neuritic, and synaptic compartments. At these interfaces, microglia preferentially internalize excitatory postsynaptic (PSD-95^+^) material from extinction ensembles and bias extinction-ensemble spine remodeling toward loss at retrieval, consistent with a microglial brake on extinction. Conversely, weakening microglial recruitment or engulfment—or removing the protection afforded to fear ensembles—accelerates extinction without disrupting fear acquisition. Together, these findings indicate that microglial processes use local ensemble-specific signals during retrieval to tune synaptic remodeling in a way that preserves fear memory while limiting extinction.

A central theme that emerges is that ATP signaling links ensemble activity to microglial recruitment and state. The soma–microglia junction^20^ provides one plausible decoding site: fear ensembles show selective enrichment of vNUT within microglia-apposed somatic microdomains and along proximal neuritic compartments, whereas extinction retrieval preferentially strengthens microglial engagement at K_V_2.1-associated junctional domains at the same interfaces. Importantly, these patterns argue for microdomain specialization rather than a simple scaling of total contact: even when overall apposition to fear ensemble bodies and neurites is reduced, the remaining microglia-apposed surface becomes selectively enriched for vNUT^+^ domains, consistent with a “find-me” signal embedded within a smaller interface. This division of labor supports a model in which vNUT-dependent ATP release from active fear-ensemble neuronal bodies and dendrites biases microglial process recruitment, while K_V_2.1-associated somatic junctions on extinction ensembles may help stabilize expanded contact zones seen during extinction retrieval.

Pharmacological and genetic evidence converges on microglial processes as the main effectors of this signaling. Chronic P2Y12 antagonism with clopidogrel mimics the pro-extinction phenotype of *P2ry12* deletion and reduces CD68 load and ensemble-derived cargo inside microglia, yet leaves gross somatic apposition to extinction ensembles and P2Y12 coverage at the neuron–microglia interface largely unchanged. This dissociation suggests that although P2Y12 is critical for orchestrating process dynamics and phagocytic loading, stabilization of somatic junctions likely depends on additional adhesion or recognition mechanisms that cooperate with purinergic signaling. In this view, ATP/P2Y12 is not the sole anchor, but a dynamic recruiter that biases process elaboration and process-associated uptake of ensemble-derived material. More broadly, our data are consistent with a model in which retrieval-locked activation of ensemble neurons—reinforced in part by NMDAR-dependent synaptic plasticity that stabilizes ensemble connectivity^10,64^—evokes vNUT-mediated ATP release from soma-proximal and dendritic compartments. ATP gradients are sensed by P2Y12 receptors on nearby microglial processes, triggering Gi-coupled changes in process surveillance and compartment engagement.^65,66^ At the same time, the preserved somatic interface after P2Y12 blockade indicates that ATP is unlikely to act alone; additional activity-dependent cues almost certainly cooperate with vNUT-derived ATP to stabilize microglia at ensemble somata and tune downstream process–synapse engagement *in vivo*.

The synaptic specificity of this editing further supports an active, instructive role for microglia. Within extinction ensembles, engulfment is biased toward excitatory postsynaptic sites, with little uptake of vGluT1^+^ presynaptic or gephyrin^+^ or vGAT^+^ inhibitory elements, consistent with a postsynaptic locus of extinction plasticity in mPFC.^67-69^ In parallel, extinction retrieval can increase microglial contact with inhibitory synapses outside ensemble-derived excitatory postsynaptic cargo without detectable ingestion—reminiscent of process-mediated “shielding” of GABAergic inputs after anesthesia^70^ and distinct from the preferential removal of inhibitory synapses reported in epilepsy models.^32,71^ These patterns argue against indiscriminate scavenging and instead point to a layered program in which microglia cull specific excitatory postsynaptic structures from extinction ensembles while interacting with inhibitory synapses in a more protective, non-engulfing mode. The tight coupling of engulfment to retrieval, its enrichment in CD68^+^ lysosomes, and its dependence on ensemble PS exposure (TMEM16F) and microglial MERTK are more consistent with receptor-guided culling than with passive collection of neuron-shed material, in line with recent conceptual distinctions between microglia as hunters versus gatherers of synapses.^72^

Our data further highlight the PS–Gas6–MERTK signaling as an operative phagocytic mechanism in this physiological setting. Complement-dependent pruning is typically linked to developmental refinement and inflammatory disease contexts,^28-32,73-75^ whereas MERTK can respond directly to activity-dependent PS externalization. We find that extinction ensembles preferentially expose PS, in part via TMEM16F, and that microglial MERTK is required for the behavioral and structural manifestations of extinction-linked pruning. Moreover, disrupting TREM2, another microglial receptor associated with phagocytic activation,^62^ does not measurably affect extinction, supporting pathway specificity in this context. Targeting the PS–MERTK signaling may therefore provide a way to modulate extinction-related synaptic remodeling without broadly engaging complement or systemic immune pathways.

A complementary conclusion is that fear memory is actively protected by a “don’t-eat-me” gate. Fear retrieval elevates CD47 on fear ensembles and increases microglial SIRPα; releasing this brake—via SIRPα blockade or ensemble-restricted CD47 deletion—heightens microglial engulfment of fear-ensemble material and promotes extinction. Developmental studies show that loss of CD47 or SIRPα augments microglial pruning and reduces synapse number, with CD47 preferentially enriched at highly active synapses.^35,63^ Conversely, reduced microglial SIRPα in neurodegenerative models associates with excessive pruning and cognitive decline.^76,77^ Our results extend this mechanism to memory circuits, suggesting that CD47–SIRPα normally preserves fear-ensemble connectivity during extinction but, if exaggerated, could contribute to extinction-resistant fear as in PTSD.^78^ Because CD47–SIRPα also operates in peripheral immunity, coordinated regulation across brain and periphery is plausible and raises the possibility that peripheral measures of this pathway might report central ensemble states.

In summary, our work supports a simple organizing principle for how microglia influence competing memory traces in the adult brain. By combining activity-dependent ensemble tagging with compartment-resolved imaging and targeted perturbations, we show that microglial processes use local “find-me,” “eat-me,” and “don’t-eat-me” cues to bias synapse editing toward extinction ensembles while sparing fear ensembles. In this way, microglia preserve fear memory by limiting the structural and functional consolidation of extinction, linking molecular recognition at the neuron–microglia interface to competition between fear and extinction at the ensemble level. This framework points to discrete, mechanistically grounded targets—and the retrieval window itself—as opportunities to enhance extinction without broadly disrupting fear learning. Future work should resolve the nanoscale sequence of synapse removal *in vivo*, determine how these mechanisms generalize across circuits, sex, and stress history, and test whether transient, ensemble-targeted modulation of microglial editing can safely promote adaptive memory updating in models of anxiety and PTSD.

## STAR★METHODS

Detailed methods are provided in the online version of this paper and include the following:

### KEY RESOURCES TABLE

### RESOURCE AVAILABILITY

Lead contact

Materials availability

Data and code availability

### EXPERIMENTAL MODEL AND SUBJECT DETAILS

Mice

### METHOD DETAILS

Drug administration

Guide cannula and osmotic pump implantation, and intracerebral microinjection

Auditory fear conditioning

Whole-cell patch-clamp recordings

sgRNA Plasmid Construction

Chronic two-photon cranial window implantation

*In vivo* two-photon imaging of microglia and ensemble neurons

Image processing and quantitative analysis

Immunohistochemistry and imaging

Quantification of c-Fos fluorescence intensity

3D super-resolution analysis of microglia–ensemble interactions

### QUANTIFICATION AND STATISTICAL ANALYSIS

### SUPPLEMENTAL INFORMATION

Supplemental Information includes 19 figures and can be found with this article online.

### ACKNOWLEDGMENTS

We would like to thank Drs. Yu Kong and Xu Wang (Electron Microscopy Facilities of Center for Excellence in Brain Science and Intelligence Technology, Chinese Academy of Sciences) for assistance with EM sample preparation and image analysis. The authors express their gratitude and respect to all animals sacrificed in this study. During the preparation of this manuscript, the authors, as the non-native English speakers, used ChatGPT 5.2 to improve the language and enhance its readability. This study was supported by grants from the STI2030-Major Projects (2021ZD0202800, 2022ZD0204700), the National Natural Science Foundation of China (32371078), the Program of Shanghai Academic/Technology Research Leader (22XD1420700), the Shanghai Municipal Health Commission (2022XD046), Shanghai Pilot Program for Basic Research (21TQ014), Innovative Research Team of High-Level Local Universities in Shanghai, Changping Laboratory (2025B-07-18).

### AUTHOR CONTRIBUTIONS

Y.-L.W., B.P., and W.-G.L. conceptualized the study; Y.-L.W., Y.C., T.L., T.-T.S., Y.Z., Z.J., Xiaoyu Yang, S.-Y.W., J.-Z.R., F.-X.Z., Y.H., S.-J.L., W.X., Xin Yi, Y.-J. W., H.S., J.W., Y.J., Y.-X.L., X.-N.L.,

T.-L.X., B.L., and P.Y. performed experimental research, data analysis and interpretation, Y.-L.W., B.P., and W.-G.L. wrote the manuscript with contributions from all authors. All authors read and approved the final manuscript.

### INCLUSION AND DIVERSITY

We support inclusive, diverse, and equitable conduct of research.

### KEY RESOURCES TABLE

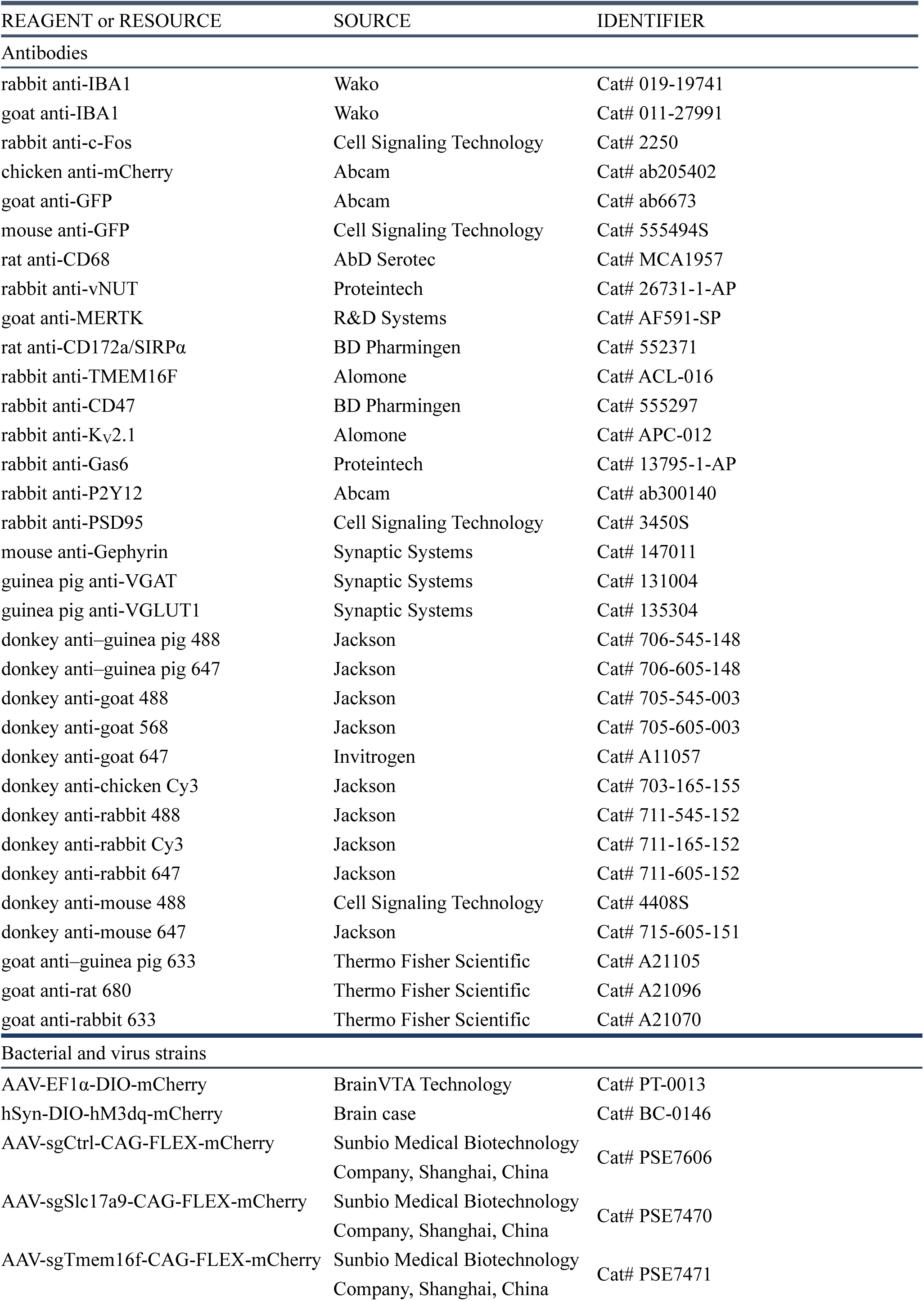
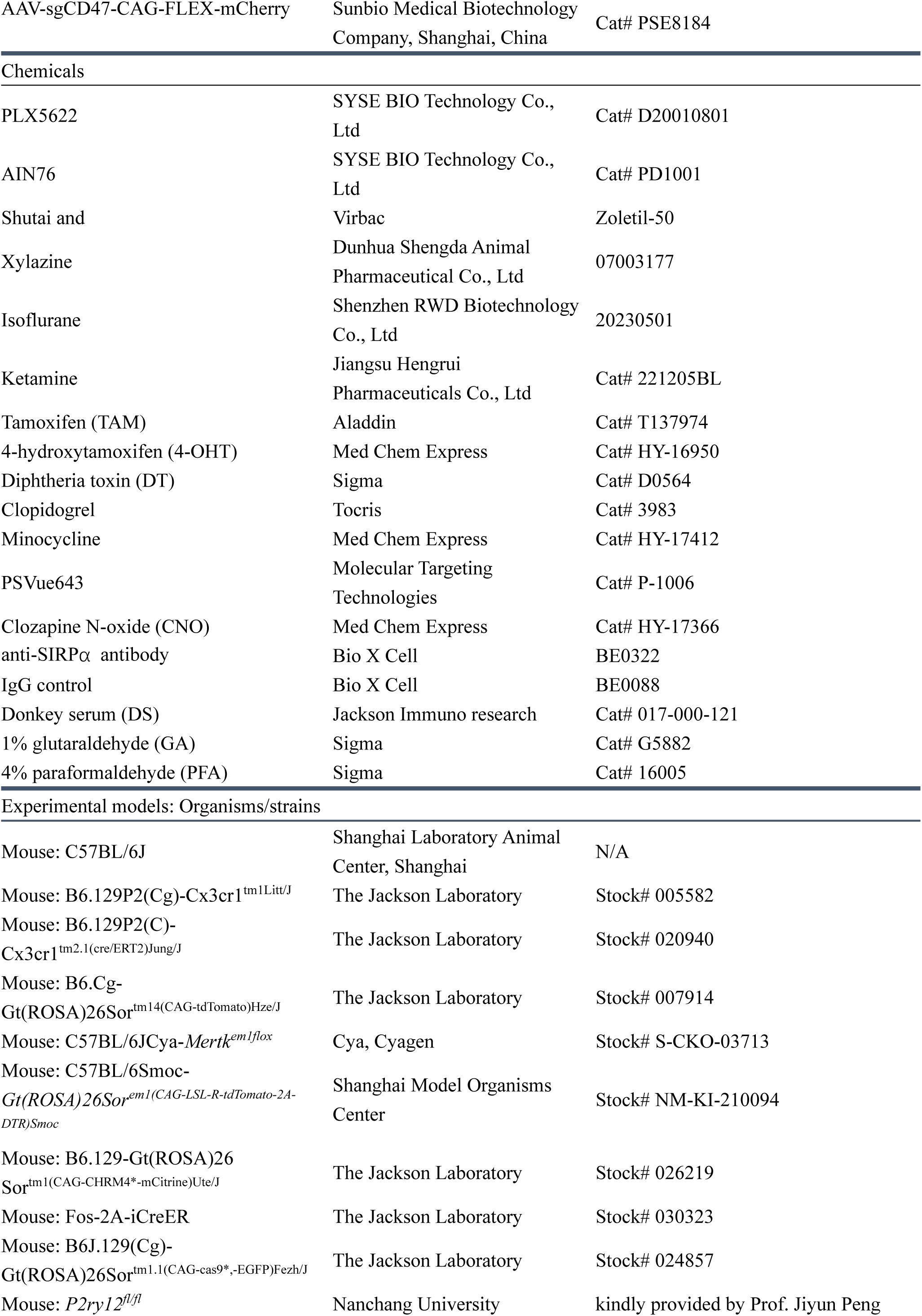
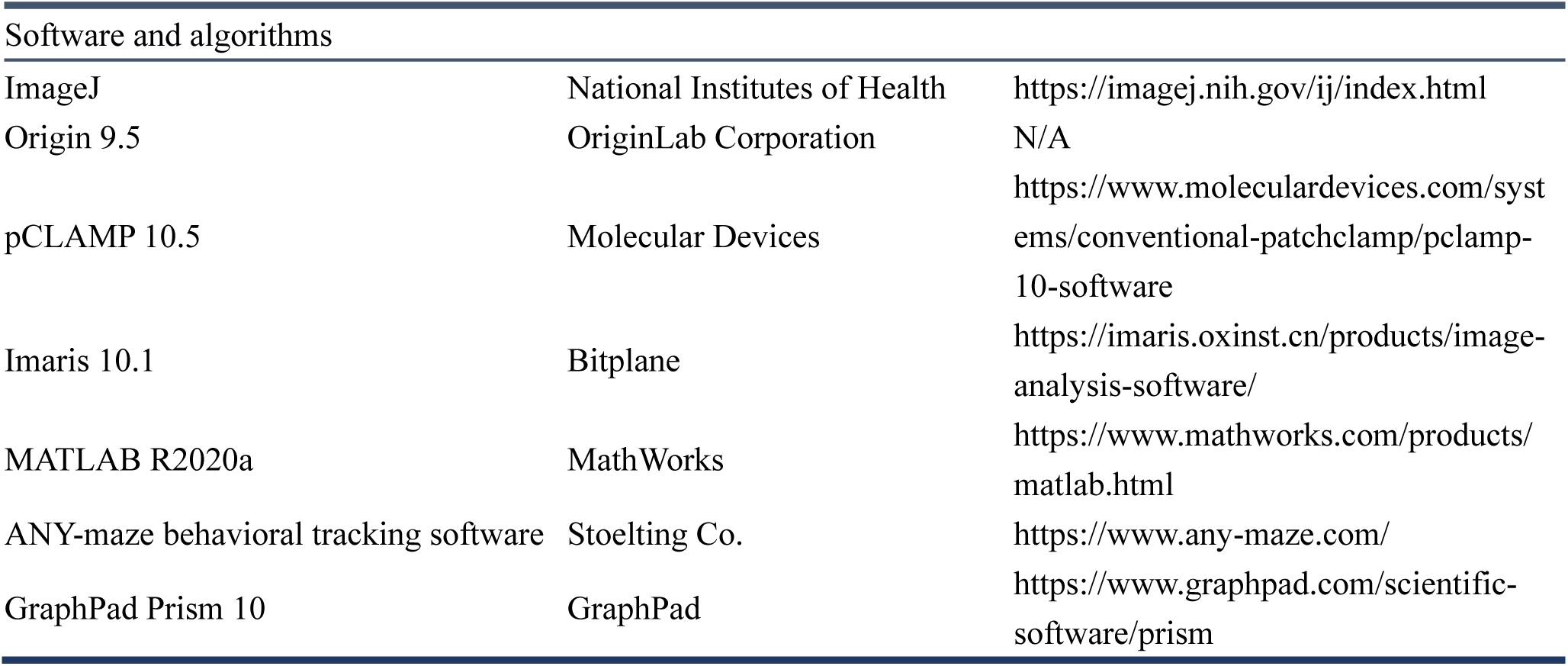

### RESOURCE AVAILABILITY

#### Lead contact

Further information and any related requests should be directed to and will be fulfilled by the lead contact, Prof. Wei-Guang Li.

#### Materials availability

This study did not generate new unique reagents.

#### Data and code availability

All data needed to evaluate the conclusions of the present study are present in the main paper and/or the supplemental information. This paper did not produce original code. Any additional information required for reanalyzing the reported data is available from the lead contact upon request.

### EXPERIMENTAL MODEL AND SUBJECT DETAILS

#### Mice

All animal procedures were approved by the Institutional Animal Care and Use Committee of the Department of Laboratory Animal Science, Fudan University (protocol 2021JS-NZHY-002), and were carried out in accordance with institutional and national guidelines for the care and use of laboratory animals. Mice were maintained under specific pathogen–free (SPF) conditions in the Fudan University Laboratory Animal Center. Unless otherwise stated, adult mice were 8–12 weeks old at the time of experimentation. Animals were housed in groups under a 12 h light/dark cycle (lights on 07:00–19:00) at 21–24 °C and 50–60% relative humidity, with food and water available *ad libitum*. All behavioral testing was performed during the light phase. Wild-type C57BL/6J mice were obtained from Shanghai Jihui Laboratory Animal Co., Ltd. The following C57BL/6-background transgenic lines were used: CX3CR1-GFP (B6.129P2(Cg)-Cx3cr1^tm1Litt/J^; The Jackson Laboratory, stock 005582);^39^ CX3CR1-CreER (B6.129P2(C)-Cx3cr1^tm2.1(cre/ERT2)Jung/J^; The Jackson Laboratory, stock 020940);^53^ Ai14 (B6.Cg-Gt(ROSA)26Sor^tm14(CAG-tdTomato)Hze/J^; The Jackson Laboratory, stock 007914);^41^ *Mertk^fl/fl^* (C57BL/6JCya-*Mertk^em1flox^*/Cya, Cyagen, stock S-CKO-03713);^48^ R26-CAG-LSL-tdTomato-2A-DTR (C57BL/6Smoc-*Gt(ROSA)26Sor^em1(CAG-LSL-R-tdTomato-2A-DTR)Smoc^*, Shanghai Model Organisms Center, stock NM-KI-210094); *P2ry12^fl/fl^*(kindly provided by Prof. Jiyun Peng, Nanchang University); LSL-hM4Di-YFP (B6.129-Gt(ROSA)26Sor^tm1(CAG-CHRM4*-mCitrine)Ute/J^; The Jackson Laboratory, stock 026219);^79^ FosTRAP2 (Fos-2A-iCreER; The Jackson Laboratory, stock 030323);^37^ Rosa26-CAG-LSL-Cas9-GFP (B6J.129(Cg)-Gt(ROSA)26Sor^tm1.1(CAG-cas9*,-EGFP)Fezh/J^; The Jackson Laboratory, stock 024857). Breeding was performed in-house, and experimental cohorts were generated by crossing the lines above as required (for example, TRAP2::Ai14;;CX3CR1-GFP, and CX3CR1-CreER::*P2ry12^fl/fl^*). For PLX5622-mediated microglial depletion experiments, only male mice and their corresponding controls were used, owing to known sex differences in depletion efficiency.^80^

### METHOD DETAILS

#### Drug administration

##### PLX5622 (chow delivery)

PLX5622 is a highly selective, orally bioavailable colony-stimulating factor 1 receptor (CSF1R) inhibitor that crosses the blood–brain barrier. For microglial depletion, PLX5622 was premixed into AIN-76A rodent chow at 1.2 g/kg and provided *ad libitum*.^52^ PLX5622 chow and the matched control AIN-76A diet were purchased from Shuyi Shuer Biotechnology (Changzhou, China).

##### Clozapine-N-oxide (CNO)

Designer receptors exclusively activated by designer drugs (DREADDs; hM3Dq or hM4Di) were manipulated *in vivo* with clozapine-N-oxide (CNO; MedChemExpress, HY-17366A). CNO was dissolved at 10 mg/ml in DMSO to generate a stock solution, diluted in sterile saline immediately before use, and administered intraperitoneally (i.p.) at 1–3 mg/kg.^81^ Behavioral testing, histology, or *ex vivo* whole-cell electrophysiological recordings were performed 30 min after injection.

##### Diphtheria toxin (DT)

For transient genetic ablation of microglia in CX3CR1-CreER::ROSA26iDTR mice, diphtheria toxin (DT; Sigma, D0564) was dissolved in sterile saline and injected i.p. at 0.03 mg/kg once every 24 h, beginning 4 days before behavioral testing and continued thereafter as required by the experimental design.^23,82^

##### Tamoxifen (TAM)

To induce CreER-dependent recombination in CX3CR1-CreER and other CreER lines, tamoxifen (Aladdin, T137974) was dissolved in corn oil at 20 mg/ml. Mice received 150 mg/kg/day by oral gavage for 7 consecutive days. Behavioral and immunohistochemical experiments were performed 28 days after the final dose to allow for recombination and clearance of peripheral myeloid cells.^48^

##### 4-Hydroxytamoxifen (4-OHT)

Activity-dependent tagging of neuronal ensembles was achieved using TRAP2-based 4-OHT induction. 4-OHT (MedChemExpress, HY-16950) was initially dissolved in absolute ethanol at 20 mg/ml and stored at −20 °C for short-term use. Immediately before injection, the stock was re-dissolved in ethanol, mixed with two volumes of corn oil, and ethanol was evaporated under vacuum to yield a 10 mg/ml suspension stored at 4 °C. 4-OHT was administered i.p. at behaviorally defined time points to couple CreERT2 nuclear translocation and recombination to the desired activity window. To minimize transport-induced immediate-early gene expression, mice were moved from the housing room to the behavioral suite at least 1 h before dosing.

##### Minocycline

Minocycline (MedChemExpress, HY-17412), a brain-penetrant tetracycline derivative with anti-inflammatory actions, was delivered in the drinking water at 0.5 mg/ml supplemented with 5% sucrose to improve palatability. This regimen yields an estimated intake of ∼100 mg/kg/day.^23^ Drug-containing water was provided *ad libitum* and refreshed every 2–3 days.

##### SIRPα-blocking antibody

To antagonize microglial SIRPα, mice were implanted with an osmotic minipump (model 1002, RWD Life Science) connected to a cannula targeting the lateral ventricle. Pumps continuously infused either a monoclonal anti-SIRPα antibody (Bio X Cell, BE0322) or isotype-matched IgG control (Bio X Cell, BE0088) at 10 µg/µl for 14 days. Behavioral training began 3 days after pump implantation to allow recovery.

##### P2Y12 receptor antagonist (clopidogrel, intracerebroventricular infusion)

Central P2Y12 signaling was blocked by continuous intracerebroventricular (i.c.v.) infusion of clopidogrel. Mice were stereotaxically implanted with an osmotic minipump (model 1002, RWD Life Science) coupled to a lateral ventricle cannula. Pumps delivered either clopidogrel (4 µg/µl in vehicle) or vehicle alone for 14 days. Behavioral procedures commenced 72 h after implantation, once animals had recovered from surgery.

##### PSVue 643 (in vivo labeling of externalized phosphatidylserine)

Externalized phosphatidylserine (ePS) was visualized *ex vivo* using PSVue 643 (P-1006, Molecular Targeting Technologies), a cell-impermeant PS-binding probe.^60,83^ Under aseptic conditions, PSVue was injected into the lateral ventricle through a guide cannula. Twenty-four hours later, acute mPFC tissue was collected, and sections were imaged immediately using confocal microscopy with excitation/emission settings matched to the probe’s spectral properties.

#### Stereotaxic viral injections into brain regions and ventricles

Mice were anesthetized by intraperitoneal injection of a mixed anesthetic (Shutai and xylazine; 6 ml/kg body weight). After loss of pedal and corneal reflexes was confirmed, the scalp was shaved and the animal was secured in a stereotaxic frame. The skin was disinfected with povidone–iodine, incised under aseptic conditions, and the skull surface was cleaned with sterile saline. The skull was leveled in the anterior–posterior and medial–lateral axes using Bregma and Lambda as reference points. Stereotaxic coordinates were defined relative to Bregma using the Allen Mouse Brain Atlas. Small craniotomies were drilled at the target locations, and a pulled borosilicate glass micropipette preloaded with viral suspension was lowered to the desired depth. Virus was infused at 0.1 μl/min using a microinfusion pump. The pipette was left in place for 10 min after the end of the infusion to allow diffusion and minimize backflow, and then withdrawn slowly. The scalp was sutured and disinfected again with povidone–iodine. Mice were placed on a heating pad and monitored until fully ambulatory before returning to their home cages. Unless indicated otherwise, the following coordinates (relative to Bregma) were used: medial prefrontal cortex (mPFC), anteroposterior (AP) +1.75 mm, mediolateral (ML) ±0.40 mm, dorsoventral (DV) −2.70 mm; lateral ventricle (intracerebroventricular, i.c.v.), AP −0.40 mm, ML ±1.00 mm, DV −2.50 mm (DV measured from the dura).

#### Guide cannula and osmotic pump implantation, and intracerebral microinjection

Anesthesia and stereotaxic positioning were performed as above. The scalp was depilated, disinfected with povidone–iodine, and incised to expose the skull. Residual periosteum and soft tissue were gently removed; a small amount of 30% hydrogen peroxide was applied to the skull to clear remaining tissue, taking care to avoid contact with surrounding skin. Two anchor holes were drilled away from the planned target site and miniature self-tapping screws were inserted to secure the head cap.

For lateral ventricle targeting, coordinates were AP −0.40 mm, ML ±1.00 mm, DV −2.50 mm (from dura). A burr hole matching the outer diameter of the stainless-steel guide cannula was drilled at the target location. The cannula was lowered along the z-axis to the target depth using the stereotaxic manipulator and fixed in place with dental acrylic (dental acrylic powder mixed with denture base resin) to form a head cap that also covered the anchor screws. After the cement had fully cured, the animal was removed from the frame. Postoperative care included routine monitoring and administration of antibiotics for 3 consecutive days.

For intracerebral microinjection via the guide cannula in awake mice, solutions were delivered at 0.5 μl/min using a microinfusion pump. The internal injector extended slightly beyond the guide tip and was left in place for 10 min after infusion to limit reflux along the cannula track.

For continuous intracerebroventricular delivery, osmotic minipumps (model 1002, RWD Life Science) were primed and filled according to the manufacturer’s instructions and connected to the lateral-ventricle cannula via flexible tubing. The pump was implanted subcutaneously on the dorsal flank, and the cannula–pump assembly was secured with dental cement. Pumps provided continuous i.c.v. infusion for the programmed duration as specified in each experiment (e.g., P2Y12 antagonist, SIRPα-blocking antibody).

#### Auditory fear conditioning

The auditory fear conditioning and extinction process were executed using the Ugo Basile Fear Conditioning System (UGO BASILE srl, Model 46250). Mice were hand-habituated for 3 consecutive days and then acclimated to the conditioning chamber (17 × 17 × 25 cm) for 20 min on the following day. Mice were transferred from the housing room to the behavioral suite at least 1 h before testing to minimize transport-induced stress. Fear acquisition was performed in Context A (black-and-white grid walls, stainless-steel grid floor connected to a shock generator, background odor of 75% ethanol). After a 3-min acclimation, mice received five pairings of a conditioned stimulus (CS; 4 kHz tone, 75 dB, 30 s) and an unconditioned stimulus (US; 0.75 mA foot shock, 2 s). The US was delivered 2 s before CS offset (i.e., co-terminating with the CS), and inter-trial intervals (ITIs) were pseudo-randomly varied between 20 and 180 s. For CS-only controls, mice were exposed to the identical CS protocol in Context A without foot shocks.^11^ After the final trial, mice remained in the chamber for 1–3 min before being returned to their home cages. Chambers were wiped with 75% ethanol between animals.

Twenty-four hours after fear (or non-fear) acquisition, extinction training was conducted over 3 consecutive days in a distinct Context B (gray walls, smooth plastic floor, background odor of 4% acetic acid). Following a 3-min acclimation period, mice received 12 CS-alone presentations (same tone parameters, no shock) with ITIs of 20–180 s. After the last CS, mice remained in the chamber for 1–3 min and were then returned to their home cages. Chambers were cleaned with water after each session. After completion of fear or extinction protocols, memory retrieval tests were carried out in Context B. Following a 3-min acclimation, mice received 4 CS presentations (ITI, 20–180 s) to assess expression of fear and extinction memory. All sessions were recorded with an infrared-sensitive CCD camera and analyzed offline using ANY-maze software. Freezing was defined as the absence of all movement except respiration for ≥ 2 s and was used as the primary index of conditioned fear.^10^ Freezing time during each CS was expressed as a percentage of the CS duration. During acquisition, freezing was quantified on a trial-by-trial basis; during extinction and retrieval, freezing was averaged in blocks of four CSs to yield within-session extinction curves and retrieval performance.

#### Whole-cell patch-clamp recordings

Following behavioral testing, mice were deeply anesthetized by intraperitoneal injection of a mixed anesthetic (Shutai and xylazine; 6 ml/kg body weight) and then rapidly killed, and the brain was quickly removed into ice-cold artificial cerebrospinal fluid (ACSF; in mM: 125 NaCl, 2.5 KCl, 12.5 D-glucose, 1 MgCl_2_, 2 CaCl_2_, 1.25 NaH_2_PO_4_, and 25 NaHCO_3_, pH 7.35-7.45) continuously bubbled with 95% O₂/5% CO₂. Coronal slices (300 µm) containing the mPFC were cut in this solution using a vibratome (Leica VT1200S) and transferred to a holding chamber containing ACSF at 34 °C for 45 min. For recordings, slices were placed in a submerged chamber and superfused with carbogen-equilibrated ACSF (∼60 ml/h) maintained at 34 ± 1 °C. Neurons were visualized with infrared differential interference contrast (IR-DIC) optics. Patch pipettes (3–6 MΩ) were pulled from borosilicate glass and recordings were made with a MultiClamp 200B amplifier (Molecular Devices), digitized at ≥10 kHz and filtered at 2–5 kHz (Bessel). After break-in, series resistance (R_s_; 8–25 MΩ) was monitored throughout and compensated by ≤20%; cells were excluded if R_s_ changed by >20% or leak current exceeded 300 pA at −70 mV.

##### Neuronal excitability (current-clamp)

Action potentials were recorded in current-clamp mode using a K^+^-based internal solution (in mM: 145 potassium gluconate, 5 NaCl, 10 HEPES, 2 MgATP, 0.1 Na_3_GTP, 0.2 EGTA, and 1 MgCl_2_, pH 7.2 with KOH, 280–300 mOsm). 500-ms step currents (−25 to +250 pA, 25 pA increments) were injected to assess firing threshold, frequency–current (F–I) relationship, spike amplitude, *dV/dt*, and half-width.

##### Evoked synaptic currents and NMDAR/AMPAR ratio (voltage-clamp)

Electrical stimulation-evoked EPSCs and NMDAR/AMPAR components were recorded in voltage-clamp using a Cs^+^-based internal solution (in mM: 132.5 Cs-gluconate, 17.5 CsCl, 2 MgCl_2_, 0.5 EGTA, 10 HEPES, 2 Na_2_ATP, with pH adjusted to 7.3 using CsOH, and osmolarity set at 280–290 mOsm). The stimulations were delivered through an ISO-Flex stimulus isolator (A.M.P.I.) with bipolar tungsten stimulating electrode placed in the target layer, and EPSCs were evoked with single pulses (0.1-ms duration, 50–200 pA) delivered at 20-s intervals. AMPAR–mediated EPSCs were measured at −70 mV as peak inward currents. NMDAR–mediated EPSCs were recorded at +40 mV. NMDAR currents were quantified as the average EPSC amplitude from 60 to 65 ms after the onset of stimulation. For each mPFC neuron, stimulation of the same intensity and duration was used to record the AMPAR- and NMDAR-mediated oEPSCs.

#### sgRNA Plasmid Construction

To enable efficient CRISPR-based gene disruption in ensemble neurons, we constructed AAV vectors expressing pairs of single-guide RNAs (sgRNAs) targeting each gene of interest (Slc17a9, Tmem16f, and Cd47). For each gene, two distinct sgRNAs were designed and cloned into a single plasmid so that both guides were expressed from independent human U6 (hU6) promoter cassettes. These dual-sgRNA expression cassettes were assembled into a modified pAAV-Flex-CAG-mCherry backbone (a gift from Edward Boyden; Addgene plasmid #28307) using a combination of Golden Gate assembly and restriction enzyme–based cloning. All constructs were verified by Sanger sequencing and then packaged into adeno-associated virus (AAV) for *in vivo* delivery. The sgRNA target sequences used in this study were: Slc17a9: sgSlc17a9-1: 5’-CGGCAGCAGAGGATACACGG-3’, sgSlc17a9-2: 5’-TCAGACCTAGGACAACCCAC-3’; Tmem16f: sg sgTmem16f-1: 5’-GCTGGAGGAGGACGACGATG-3’, sgTmem16f-2: 5’-CCTGCTGAACATGGAGCTGG-3’; Cd47: sgCd47-1: 5’-CACTTCATGCAATGAAACTG-3’, sgCd47-2: 5’-TTGCATCGTCCGTAATGTGG-3’.

#### Chronic two-photon cranial window implantation

All surgeries were performed under isoflurane anesthesia (induction 3%, maintenance 1.2–1.5%) with animals maintained on a feedback-controlled heating pad. Mice were secured in a stereotaxic frame, the scalp was disinfected with povidone–iodine and incised, and the skull was cleaned of periosteum. For chronic access to mPFC, a circular craniotomy (≈3.4 mm diameter) was made over the midline (centered at AP +1.7 mm, ML 0.4 mm relative to Bregma) using a high-speed dental drill, leaving a thin outer bone layer to minimize heat and vibration. The bone flap was gently lifted with fine forceps, taking care to preserve the dura, and the exposed cortex was irrigated with sterile saline; minor bleeding was controlled with absorbable hemostatic material. For 45° microprism implantation,^84^ a right-angle glass prism was affixed to a D263T coverslip (3.5 mm diameter, 150-µm thickness). The dura was carefully opened near the midline, and the prism–coverslip assembly was lowered at 45° to obtain optical access to the mPFC with minimal cortical displacement. The coverslip was seated flush with the skull and sealed at the margins with tissue adhesive (3M Vetbond) and dental acrylic. A custom titanium headplate was then attached to the skull with cyanoacrylate adhesive (ergo 5800) and dental cement to provide long-term stability for repeated imaging. Postoperative care followed standard institutional guidelines. Mice received daily intraperitoneal injections of dexamethasone sodium phosphate and ceftiofur sodium for 7 days, and surgical sites and body weight were monitored until animals fully recovered and exhibited normal behavior.

#### *In vivo* two-photon imaging of microglia and ensemble neurons

*In vivo* imaging began 4–8 weeks after window implantation. Data were acquired on an Evident FVMPE-RS gantry two-photon microscope equipped with a dual-output femtosecond pulsed laser (Newport Insight X3 Dual; one tunable arm, 690–1,300 nm; one fixed at 1,045 nm) using galvanometric scanning.^43,44^ For structural imaging of microglia and dendritic spines, mice were anesthetized with ketamine/xylazine (87 mg/kg i.p.) and imaged once a stable plane of anesthesia was reached.TRAP2::Ai14;;CX3CR1-GFP mice were used to visualize microglia (GFP) and extinction ensemble neurons (tdTomato). Microglial process dynamics were imaged at 920 nm through the microprism using a 16× water-immersion objective (NA ≥ 0.8; Olympus) with a field of view of 127.3 × 127.3 µm^2^ at 512 × 512 pixels. Prior to dual-color imaging, time-lapse Z-stacks (0.5 µm step size) of GFP+ microglia were acquired in a single channel every 5 minutes to record their dynamics. tdTomato-labeled dendrites and spines were imaged at 1,040 nm with a 25×/1.05 NA water-immersion objective (Olympus) with a field of view of 84.9 × 84.9 µm^2^ at 512 × 512 pixels. Excitation power at the brain surface was kept as low as possible and increased only gradually with depth (typically 30–40 mW at the imaging plane) to limit photodamage. The mPFC was sampled at depths of ∼100–150 µm below the pial surface via the 45° prism.

To couple microglial dynamics to extinction behavior, mice underwent standard fear conditioning and 3 days of extinction training. On the third extinction day, 4-OHT was injected intraperitoneally to induce FosTRAP-dependent permanent tdTomato labeling of extinction ensemble neurons. After a 4–8 week expression period, mice were re-exposed to the extinction context in a protocol consisting of 2 baseline days, 1 re-extinction day, and 1 extinction retrieval day. Imaging was performed for 30 min beginning 1 h after each daily behavioral session. Fields of view were matched across days by reference to surface vasculature and stable cell clusters, allowing longitudinal tracking of the same microglia and ensemble dendritic segments. For awake imaging (when performed), mice were habituated to head fixation for ≥ 3 days (30 min per day) prior to data collection to reduce stress and motion.

#### Image processing and quantitative analysis

All imaging data were analyzed using ImageJ (NIH) and Imaris 10.1 (Bitplane).

##### Microglial morphology and dynamics

Z-stacks and time-series were first corrected for motion along the x–y and z axes using the StackReg/TurboReg plugins in ImageJ.^85^ For each microglial cell, a 14-µm-thick stack (20 optical sections) centered on the soma was collapsed into a maximum-intensity projection (MIP), from which total cell area, soma area, and process area were extracted. Surveillance territory was defined by manually outlining a polygon connecting the most distal process tips on the MIP; the enclosed area was taken as the surveillance area. To assess process motility, images acquired at 0 and 30 min were binarized and registered, and “moving area” was defined as the area occupied at either time point but not both; the moving-area fraction was expressed relative to total microglial area at 30 min. The number of distal swellings/bulbous endings was counted manually per cell. Sholl analysis of arbor complexity was performed in Imaris using concentric shells centered on the soma to quantify branch intersections as a function of radius.

##### Dendritic spine dynamics and microglia–spine contacts

tdTomato-positive dendritic branches from identified extinction ensemble neurons were traced in 3D across sessions. For quantitative analysis, dendritic segments of 30 ± 3 µm were selected and matched across time points using fiduciary landmarks. Protrusions extending beyond one-third of the local shaft diameter were considered; filopodia were defined as long, thin protrusions with a head-to-neck diameter ratio < 1.2:1 and a length-to-neck diameter ratio > 3:1 and were excluded from spine counts.^86^ All remaining protrusions were classified as spines. Spine density, formation, and elimination were quantified across days. Microglia–spine interactions were analyzed in Imaris 10.1 using the “Surfaces” module for 3D rendering of microglia and dendrites, combined with the “Coloc” tool to compute absolute contact area and contact area normalized to spine surface area. All analyses were performed single-blind by an experimenter unaware of group identity.

#### Correlative light–electron microscopy (CLEM) with near-infrared branding

To validate the ultrastructural organization of microglia–ensemble contacts, we performed CLEM with near-infrared branding (NIRB) in TRAP2::Ai14;;CX3CR1-GFP mice. Fear- and extinction-ensemble neurons were labeled by TRAP2 after the corresponding behavioral paradigms, and the resulting GFP^+^ microglia and tdTomato^+^ ensemble neurons were examined first by two-photon microscopy and then by serial electron microscopy.

##### Sample preparation and NIRB localization

Mice were deeply anesthetized and transcardially perfused with 4% paraformaldehyde (PFA) and 1% glutaraldehyde (GA) in 0.1 M phosphate buffer (PB, pH 7.4). Brains were removed and sectioned coronally into 50-µm slices on a vibratome. Slices containing mPFC were transferred to a two-photon microscope to locate GFP-labeled microglia and tdTomato-labeled ensemble somata and their contacts.

Sites of microglia–ensemble apposition were identified in z-stacks and marked by NIRB.^87^ Briefly, a widely tunable ultrafast laser was operated at 960 nm in both constant-angular-velocity “tornado” and line-scan modes at ∼90% output (≈400 mW at the focal plane) to burn square fiducial marks adjacent to, but not overlapping, the target contact sites.^70^ For somatic contacts, fields of view of ∼169.7 × 169.7 µm^2^ at 1,024 × 1,024 pixels were acquired, with z-stacks spanning ∼20–30 µm at 1-µm steps to fully capture the microglial process and ensemble soma interface. Fluorescence imaging used 960 nm excitation for GFP and 1,040 nm for tdTomato, with power maintained at 30–40 mW to preserve morphology. After light imaging and NIRB marking, slices were post-fixed in 2.5% GA in PB at 4 °C to prepare for electron microscopy.

##### Electron microscopy processing and imaging

For EM, NIRB-marked regions were dissected into small blocks and rinsed five times in 0.1 M PB (pH 7.4). Blocks were post-fixed in 1% osmium tetroxide for 1 h, washed in double-distilled H_2_O, and stained overnight in 1% aqueous uranyl acetate. The next day, samples were rinsed, dehydrated through a graded ethanol series (30%, 50%, 70%, 80%, 90%, 100%), and then infiltrated with mixtures of propylene oxide and epoxy resin (Embed 812, Electron Microscopy Sciences) before final embedding in 100% resin. Polymerization was carried out in an oven according to the manufacturer’s instructions.

Embedded blocks were trimmed under a stereomicroscope using the two-photon/NIRB fiducials as guides, yielding rectangular blocks ∼0.6–0.8 mm on a side that contained the region of interest. Serial ultrathin sections (∼100 nm) were cut on an ultramicrotome (Leica UC7) and collected on silicon wafers (∼10 × 20 mm) for array-type serial SEM.

Sections were imaged on a ZEISS GeminiSEM 300 in sample-stage deceleration mode at 2 kV accelerating voltage (landing voltage ∼3 kV). For volumetric reconstructions of somatic contacts, images were acquired at a voxel size of ∼10 × 10 × 100 nm (x–y–z) with a dwell time of 2 µs per pixel. Higher-resolution images for display were collected at ∼1 × 1 nm in the contact plane. Image stacks were exported for subsequent registration, segmentation, and 3D reconstruction.

##### CLEM registration and 3D segmentation

CLEM analysis combined two-photon and SEM data using ImageJ (NIH), Dragonfly (Object Research Systems), Photoshop (Adobe), and Imaris 10.1 (Bitplane). Two-photon z-stacks served as references to align the EM series using NIRB fiducials. After registration, serial EM images (∼100-nm section thickness) were inspected and annotated in a layer-by-layer manner.

Microglia were segmented based on established ultrastructural criteria: relatively electron-dense nuclei with clumped chromatin, narrow perinuclear space, electron-dense cytoplasm, and processes containing elongated smooth endoplasmic reticulum profiles.^20^ Ensemble neurons were identified by their large, lightly electron-dense nuclei with dispersed chromatin, prominent nucleoli, abundant rough endoplasmic reticulum and free ribosomes, and clearly defined plasma membranes. Neuronal processes were recognized by parallel microtubules, occasional mitochondria, and synaptic specializations. Using these morphological features and process trajectories, microglial somata, processes, and their apposed ensemble membranes were manually traced and rendered in 3D to quantify the geometry of somatic interfaces and associated bulbous endings.

#### Immunohistochemistry and imaging

Behaviorally tested mice were deeply anesthetized and transcardially perfused with ice-cold PBS followed by 4% PFA in 0.1 M phosphate buffer (PB, pH 7.4) until liver blanching. Brains were removed, post-fixed in 4% PFA containing 0.01% glutaraldehyde for 8–12 h at 4 °C, and sectioned coronally (40–100 µm) on a Leica vibratome. Free-floating sections were stored in cryoprotectant until staining.

For single- and multi-channel immunolabeling, sections were washed in PBS, permeabilized in 0.5% Triton X-100 for 30 min, and blocked in 10% normal donkey serum at room temperature for 2 h. Primary antibodies were applied overnight at 4 °C in PBS containing 0.3% Triton X-100 and 3% donkey serum. The following primary antibodies were used (typical dilution 1:500 unless noted): rabbit anti-IBA1 (Wako, 019-19741), goat anti-IBA1 (Wako, 011-27991), rabbit anti-c-Fos (Cell Signaling Technology, 2250), chicken anti-mCherry (Abcam, ab205402), goat anti-GFP (Abcam, ab6673), mouse anti-GFP (Cell Signaling Technology, 555494S), rat anti-CD68 (AbD Serotec, MCA1957), rabbit anti-vNUT (Proteintech, 26731-1-AP), goat anti-MERTK (R&D Systems, AF591-SP), rat anti-CD172a/SIRPα (BD Pharmingen, 552371), rabbit anti-TMEM16F (Alomone, ACL-016), rabbit anti-CD47 (BD Pharmingen, 555297), rabbit anti-K_V_2.1 (Alomone, APC-012), rabbit anti-Gas6 (Proteintech, 13795-1-AP), rabbit anti-P2Y12 (Abcam, ab300140), rabbit anti-PSD95 (Cell Signaling Technology), mouse anti-Gephyrin (Synaptic Systems, 147011), guinea pig anti-VGAT (Synaptic Systems, 131004), and guinea pig anti-VGLUT1 (Synaptic Systems, 135304).

After three washes in PBST (PBS + 0.1% Tween-20, 10 min each at room temperature), sections were incubated overnight at 4 °C with species-appropriate secondary antibodies (1:500) conjugated to spectrally distinct fluorophores (Alexa Fluor 488/568/633/647/680 or Cy3). Secondary antibodies included: donkey anti–guinea pig 647 and 488 (Jackson, 706-605-148, 706-545-148), donkey anti-goat 488/568/647 (Jackson 705-545-003, 705-605-003; Invitrogen A11057), donkey anti-chicken Cy3 (Jackson, 703-165-155), donkey anti-rabbit 488/Cy3/647 (Jackson, 711-545-152, 711-165-152, 711-605-152), donkey anti-mouse 488/647 (Cell Signaling Technology, 4408S; Jackson, 715-605-151), goat anti–guinea pig 633 (Thermo Fisher Scientific, A21105), goat anti-rat 680 (Thermo Fisher Scientific, A21096), and goat anti-rabbit 633 (Thermo Fisher Scientific, A21070). Sections were then washed three times in 0.01 M PBS and mounted in anti-fade medium.

Low-magnification survey images were acquired on an Olympus VS200 slide scanner with 10× or 20× air objectives. High-resolution imaging was performed on Olympus FV3000 or Leica STELLARIS 8 laser-scanning confocal microscopes equipped with 40×/1.3 NA and 63×/1.4 NA oil-immersion objectives. For quantitative comparisons, acquisition parameters (laser lines, power, detector gain and offset, scan speed, pixel size, z-step, and pinhole size) were held constant across animals within each experiment. Images were typically collected at 1,024 × 1,024 pixels as z-stacks and exported for analysis in Fiji (ImageJ) and Imaris 10.1 (Bitplane) using predefined pipelines for background subtraction, thresholding, and 3D reconstruction.

#### Quantification of c-Fos fluorescence intensity

To assess how microglial manipulations affect neuronal activity, c-Fos expression was quantified in mPFC after fear or extinction retrieval. Within each experimental batch (e.g., PLX5622 vs. control), all brains were processed in parallel for dual IBA1 and c-Fos immunostaining and imaged under identical confocal settings to minimize batch effects. Sections were excluded if overall c-Fos signal was markedly below the batch mean or if strong within-section heterogeneity in staining quality was observed.^88^

Imaging was performed on an Evident FV3000 confocal microscope with a 20× objective. For each mouse, 4–5 anatomically matched mPFC sections were imaged as 3D stacks (1,024 × 1,024 pixels, 1 µm z-step, ∼40 optical sections). All acquisition parameters (laser power, gain, offset, scan speed, pixel size, z-step, and bit depth) were fixed across the batch to ensure comparability.

Volumes were analyzed in Imaris 10.1 using the “Spots” module for automated detection of c-Fos^+^ nuclei (minimum spot diameter typically set to ≥8 µm). Each detected cell was represented as a sphere, and the maximum voxel intensity within that sphere was extracted as the single-cell c-Fos intensity. Using thresholds held constant within each experiment, cells were binned into low, medium, and high expression categories based on the global intensity distribution. For each animal, the proportion of c-Fos^+^ cells falling into each category was calculated relative to the total number of c-Fos^+^ cells per section. All preprocessing, detection, and binning parameters were pre-specified and kept fixed within a batch, and analyses were carried out under semi-blinded conditions.

#### 3D super-resolution analysis of microglia–ensemble interactions

To reconstruct microglia–ensemble interactions at synaptic resolution, we used TRAP2::Ai14;;CX3CR1-GFP mice in which fear and extinction ensemble neurons were tagged via TRAP2-mediated tdTomato expression in defined behavioral windows. Multichannel immunofluorescence was performed to label microglial lysosomes (CD68), synaptic markers (PSD95, Gephyrin, VGAT, VGLUT1), and molecular effectors (MERTK, Gas6, vNUT, K_V_2.1, TMEM16F, SIRPα, CD47, among others), with additional staining to boost endogenous GFP and tdTomato signals. High-resolution 3D datasets were acquired from defined mPFC layers (L2/3 and L5) on Leica STELLARIS 8 or Olympus FV3000 systems using 63× oil objectives (1,024 × 1,024 pixels, zoom ∼3, z-step 0.3–0.5 µm). Acquisition settings were matched across groups for each experiment. The overarching goal was to quantify: (i) ensemble neuron morphology, (ii) microglial morphology, (iii) spatial contacts between microglia and neuronal compartments (bodies, neurites, dendritic spines), and (iv) synapse-level engulfment of ensemble-derived material.

Image preprocessing and 3D reconstruction were performed in Imaris 10.1. Background subtraction and (where appropriate) deconvolution were applied to each channel using the batch-processing module. Microglia and ensemble neuronal bodies were rendered with the “Surfaces” and “Spots” tools. The Spots tool (typical diameter 8–12 µm) was used to detect individual neurons and microglial somata and to compute shortest distances between their centers. The Surfaces tool was used to segment cell bodies, processes, and dendrites and to extract metrics including surface area, volume, branch length and complexity, and the number and size of bulbous endings.^70^

Ensemble-derived puncta and synaptic markers (e.g., PSD95, VGLUT1, Gephyrin, VGAT) were also rendered as Spots or Surfaces to measure density, spatial distribution, and contact with microglial surfaces. Microglia–spine interactions were quantified by combining Surfaces with the “Coloc” tool to determine the absolute and normalized contact area between microglial processes and identified spines or ensemble-labeled puncta.

To quantify phagocytosis, we employed the “Split Spots into Surfaces” workflow together with colocalization analysis.^89^ Microglial Surfaces were generated and used as a 3D mask to restrict CD68 signal to lysosomes within each microglial cell. CD68^+^ volumes were then segmented as Surfaces. Within these CD68-defined compartments, ensemble-derived synaptic markers (e.g., tdTomato^+^PSD95^+^ puncta) were detected and rendered as Spots or small Surfaces. The number and total area of engulfed puncta were recorded for each microglia and expressed as absolute values and as fractions of total microglial or CD68 volume, providing indices of synaptic load and phagolysosomal content.^32^

Throughout all super-resolution analyses, thresholding criteria, smoothing, minimum voxel counts, quality filters, spectral unmixing, and deconvolution parameters were held constant within a given experiment. Acquisition metadata (laser lines, power, detector gain, scan speed, z-step, pixel size, pinhole) and full analysis workflows were documented, and all quantitative analyses were performed with the experimenter blinded to group identity.

### QUANTIFICATION AND STATISTICAL ANALYSIS

Behavioral, histological, electrophysiological, and imaging experiments were repeated across independent biological cohorts (typically ≥ 3), and most behavioral and tissue procedures were carried out by the same experimenter to reduce variability. Where feasible, raw data were independently re-analyzed and plotted under single-blind conditions by additional lab members to minimize operator bias. No formal statistical methods were used to predetermine sample size; group sizes were guided by prior studies using similar paradigms and by practical constraints on *in vivo* imaging and electrophysiology. Unless otherwise stated, *n* in figure legends refers to independent biological units (mice or cells averaged per mouse, as appropriate), and technical replicates were not used as statistical units. Data were organized, analyzed, and visualized in GraphPad Prism v9.4.0 (GraphPad Software Inc.). For each dataset, we first inspected residuals and variance to verify approximate normality and homogeneity of variance. When distributions were grossly skewed or variances were markedly unequal, data were log- or rank-transformed and nonparametric tests were run in parallel as sensitivity analyses; these checks did not alter the main conclusions.

Unless otherwise indicated, statistical tests were two-sided. Pairwise comparisons were performed with unpaired or paired Student’s *t* tests according to the experimental design. For comparisons involving more than two groups or repeated measures (e.g., freezing across sessions, current–frequency curves), we used one-way ANOVA, two-way ANOVA, or two-way repeated-measures ANOVA with factors such as treatment and time/session. When ANOVA indicated a significant main effect or interaction, we applied predefined post hoc tests (e.g., Bonferroni’s, Dunnett’s, or Sidak’s multiple-comparison procedures) as specified in the figure legends.

Outliers were identified only according to predefined technical criteria under conditions blinded to group identity (e.g., loss of cranial window clarity, gross motion artifact during imaging, unstable series resistance during patch clamp, or clear staining failure) and excluded prior to statistical testing; biological variability alone was not considered grounds for exclusion. All summary values are reported as mean ± s.e.m., and individual data points are shown wherever possible to increase transparency.

## Supplemental Figures and Legends

**Figure S1.**
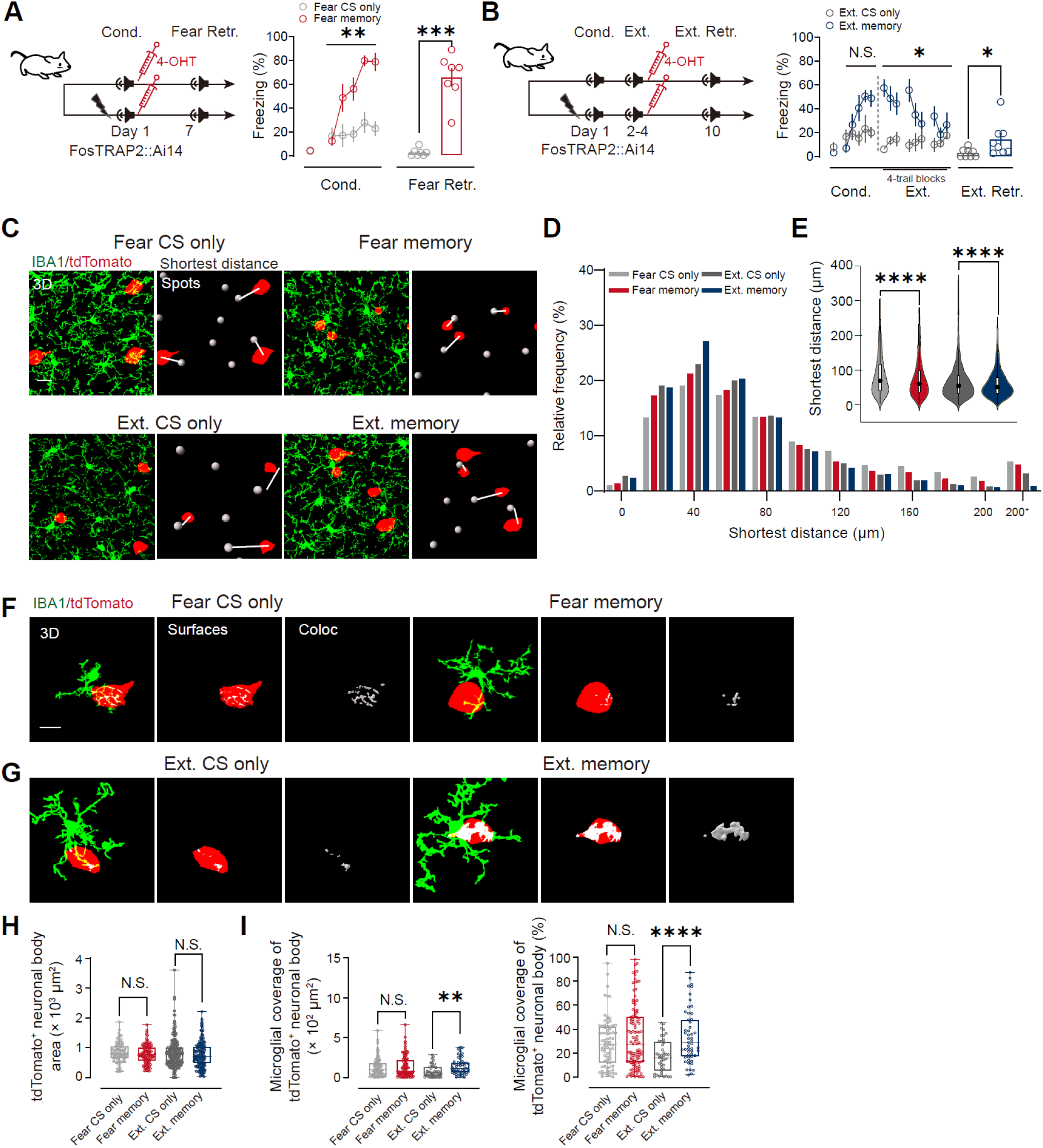
Retrieval recruits microglia toward fear- and extinction-tagged ensemble neuronal bodies, with selective strengthening of coverage of neuronal bodies during extinction retrieval, related to Figure 1. (A) Fear-ensemble tagging in FosTRAP2::Ai14 mice. 4-OHT was administered immediately after auditory fear conditioning (Cond.) to label fear-learning–activated neurons (TRAP^+^; tdTomato^+^), followed by a CS-only test (Fear CS only) or fear retrieval (Fear memory) on day 7. Freezing during CS presentations is shown (mean ± s.e.m.). Fear CS only, *n* = 11 mice; Fear memory, *n* = 7 mice. \*\**P* < 0.01, \*\*\**P* < 0.001, two-way repeated-measures ANOVA for training curves and unpaired Student’s *t* test at retrieval, as appropriate. (B) Extinction-ensemble tagging in a separate cohort of FosTRAP2::Ai14 mice. Mice underwent extinction training (Ext.; days 2–4), received 4-OHT after the third extinction session to label extinction-training–activated neurons, and were tested with CS-only (Ext. CS only) or extinction retrieval (Ext. memory) on day 10. Freezing is shown as in (A). Ext. CS only, *n* = 11 mice; Ext. memory, *n* = 11 mice. N.S., not significant; \**P* < 0.05 (statistics as in A). (C) Representative 3D reconstructions from mPFC sections immunostained for IBA1 to label microglia (green) with tdTomato^+^ ensemble neuronal bodies (tdTomato, red) across the four behavioral groups. Right panels (“shortest distance Spots”) illustrate nearest-neighbor measurements each tdTomato^+^ body to the closest IBA1^+^ microglial soma (gray spheres; white connectors). Scale bar, 20 μm. (D) Relative-frequency distributions of nearest-neighbor distances from tdTomato^+^ bodies to the closest microglial soma for each group. (E) Violin plots of the nearest-neighbor distances (data in D), showing reduced distances in both Fear memory and Ext. memory relative to their respective CS-only controls. Fear CS only, *n* = 1,665 cells (7 mice); Fear memory, *n* = 1,823 cells (7 mice); Ext. CS only, *n* = 3,063 cells (7 mice); Ext. memory, *n* = 3,096 cells (7 mice). \*\*\*\**P* < 0.0001, Mann–Whitney U test. (F and G) Representative single-cell renderings used to quantify somatic microglia–ensemble interfaces. Left, IBA1^+^ microglia (green) and a tdTomato^+^ body (red); middle, 3D surface of the tdTomato^+^ body; right, voxel-based apposition/colocalization at the microglia–soma interface (white). Scale bars, 10 μm. (H) tdTomato^+^ body area is comparable across groups. (I) Somatic interface quantification: microglial coverage of tdTomato^+^ bodies expressed as absolute interface area (left) and percent somatic coverage (right). Extinction retrieval increases somatic coverage relative to Ext. CS only, whereas fear retrieval does not increase somatic coverage relative to Fear CS only. Fear CS only, *n* = 80 cells (7 mice); Fear memory, *n* = 109 cells (7 mice); Ext. CS only, *n* = 41 cells (7 mice); Ext. memory, *n* = 52 cells (8 mice). N.S., not significant; \*\**P* < 0.01, \*\*\*\**P* < 0.0001, unpaired Student’s *t* test.

**Figure S2.**
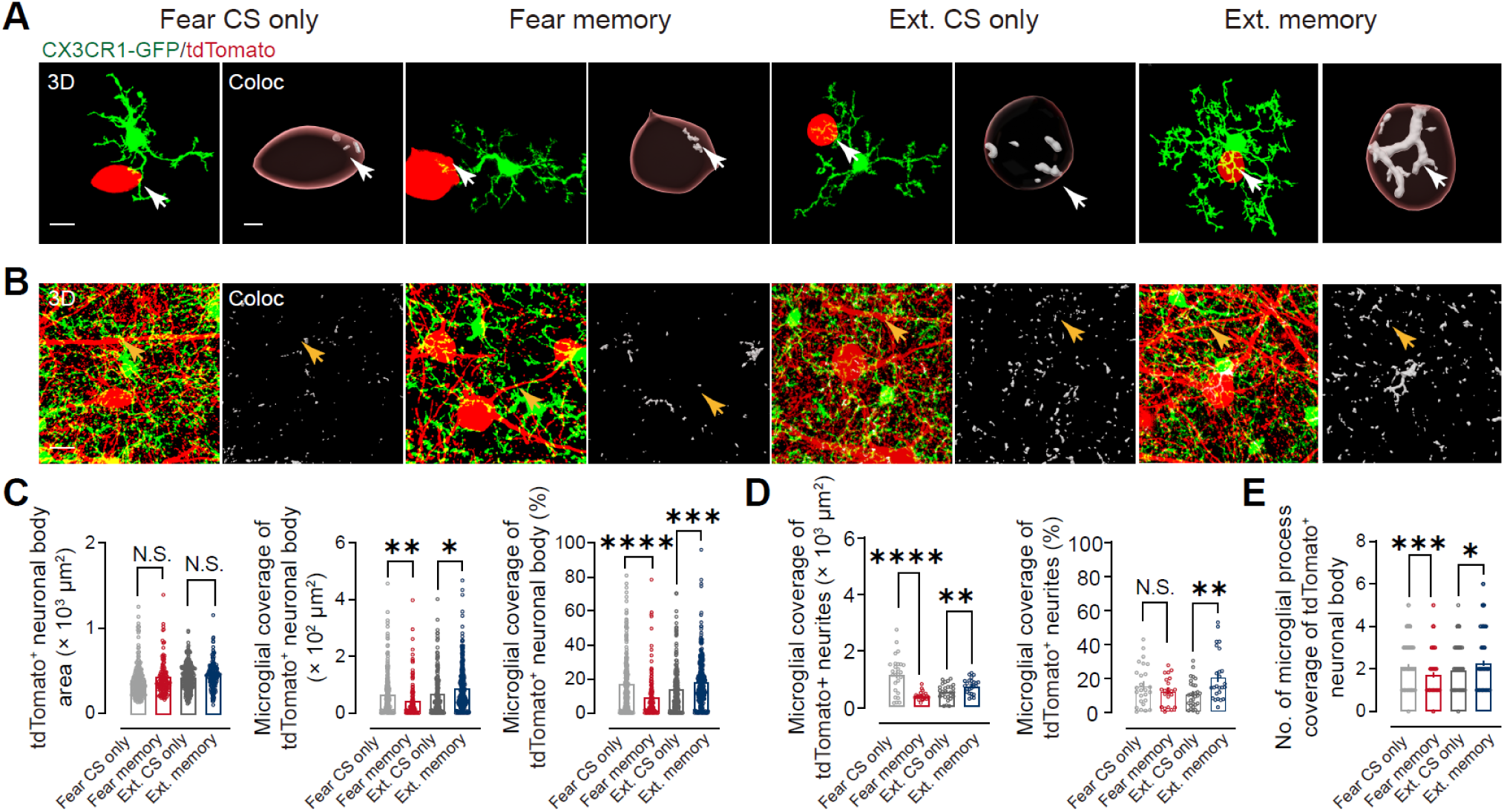
Extinction retrieval strengthens microglia–ensemble interfaces at neuronal bodies and neurites, related to Figure 1. (A) Representative 3D renderings from FosTRAP2::Ai14;;CX3CR1 mPFC showing single CX3CR1-GFP^+^ microglia (green) apposed to tdTomato^+^ neuronal bodies (tdTomato, red) in Fear CS only, Fear memory, Ext. CS only, and Ext. memory groups. *Coloc* panels depict microglia–TRAP interface voxels mapped to the tdTomato^+^ body surface (white; arrows). Scale bars, 10 μm. (B) Representative high-magnification views of microglia (green) and tdTomato^+^ structures (red) with corresponding *Coloc* maps (white; arrows) illustrating group differences in microglia–ensemble apposition. Scale bars, 10 μm. (C) Somatic interface quantification. tdTomato^+^ body area is unchanged across groups (left). Microglial coverage of tdTomato^+^ bodies (absolute interface area, middle; percent coverage, right) is reduced in Fear memory relative to Fear CS only, but increased in Ext. memory relative to Ext. CS only. Fear CS only, *n* = 229 cells (7 mice); Fear memory, *n* = 135 cells (7 mice); Ext. CS only, *n* = 188 cells (7 mice); Ext. memory, *n* = 269 cells (7 mice). (D) Neurite interface quantification. Microglial coverage of tdTomato^+^ neurites (absolute area, left; percent coverage, right) decreases after fear retrieval and increases after extinction retrieval relative to the corresponding CS-only controls. Fear CS only, *n* = 26 sections (7 mice); Fear memory, *n* = 21 sections (7 mice); Ext. CS only, *n* = 27 sections (7 mice); Ext. memory, *n* = 25 sections (7 mice). (E) Number of microglial processes apposed to each tdTomato^+^ neuronal body, showing fewer contacts in Fear memory and more contacts in Ext. memory relative to their respective controls. Same sections as in (D). N.S., not significant; \**P* < 0.05, \*\**P* < 0.01, \*\*\**P* < 0.001, \*\*\*\**P* < 0.0001, unpaired Student’s *t* test (memory versus corresponding CS-only control).

**Figure S3.**
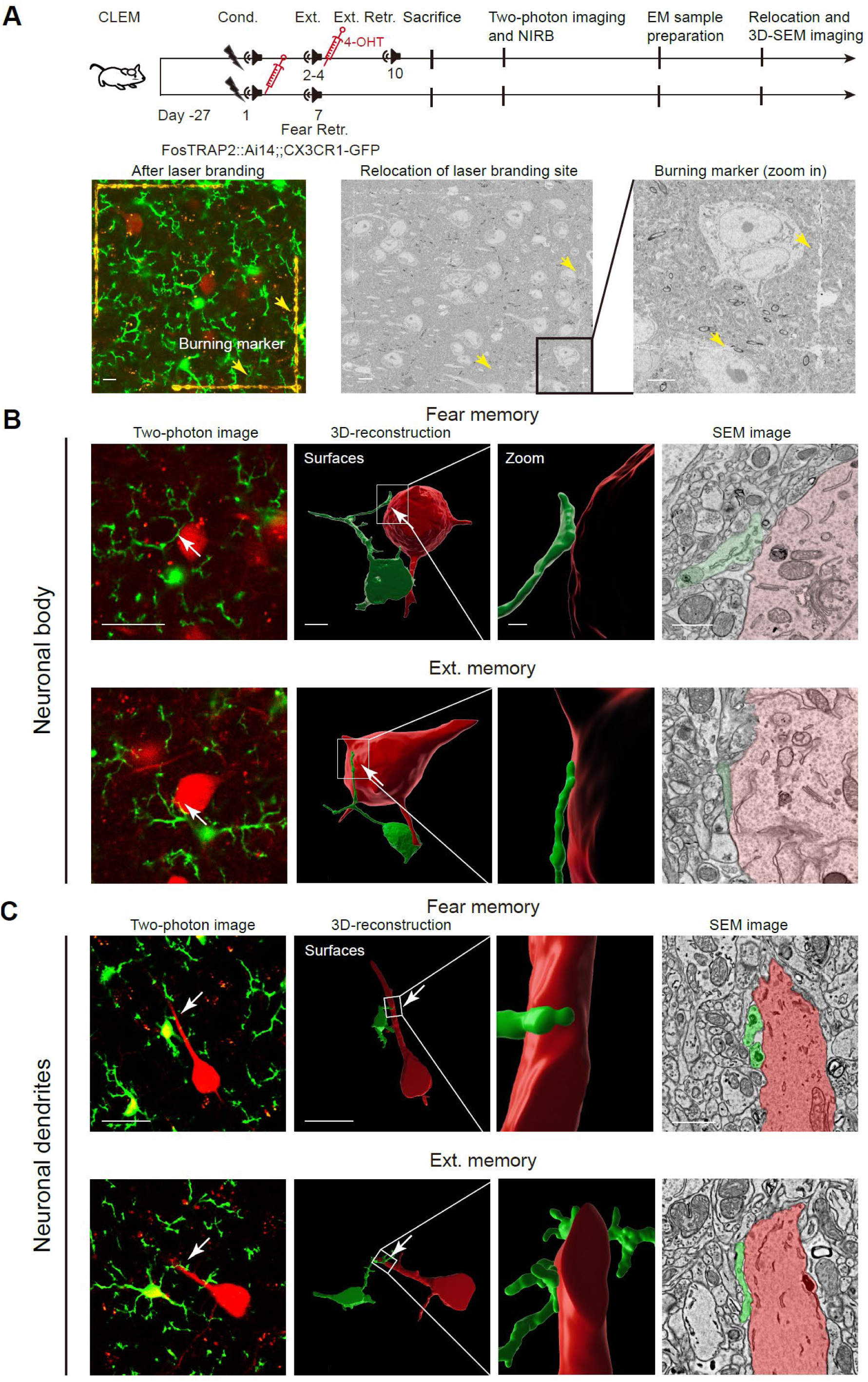
Correlative light–electron microscopy (CLEM) validates ultrastructural microglia–engram contacts, related to Figure 1. (A) Workflow for CLEM with near-infrared branding (NIRB) in FosTRAP2::Ai14;; CX3CR1-GFP mice. Fear engrams were TRAP-labeled by 4-OHT delivery after conditioning (Cond.) and examined after fear retrieval (Fear Retr., day 7). Extinction engrams were TRAP-labeled by 4-OHT delivery after the last extinction session (Ext., days 2–4) and examined after extinction retrieval (Ext. Retr., day 10). After retrieval, acute mPFC slices were imaged by two-photon microscopy to identify candidate CX3CR1-GFP^+^ microglia processes contacting tdTomato^+^ engram bodies/neurites; the region of interest was then marked in situ by NIRB, followed by EM sample preparation and relocation of the branded site for 3D-SEM acquisition and alignment across modalities. Scale bars, 10 μm (two-photon), 10 μm (EM relocation view), and 5 μm (burn mark zoom). (B and C) Representative CLEM examples of microglial processes apposed to fear (top) or extinction (bottom) engram somata or dendrites. For each condition, two-photon images (left) show GFP^+^ microglia (green) contacting tdTomato^+^ engram neurons (red); corresponding 3D surface reconstructions (second panels) and zoomed views (third panels) highlight the contact region (boxed). Serial block-face SEM sections (right) depict the ultrastructural interface between the same microglial process (false-colored green) and ensemble neuronal bodies (B) or dendrites (C) (false-colored red). Scale bars, 10 μm (two-photon), 10 μm (3D-reconstruction Surfaces), 1 μm (3D-reconstruction Zoom) and 1 μm (SEM).

**Figure S4.**
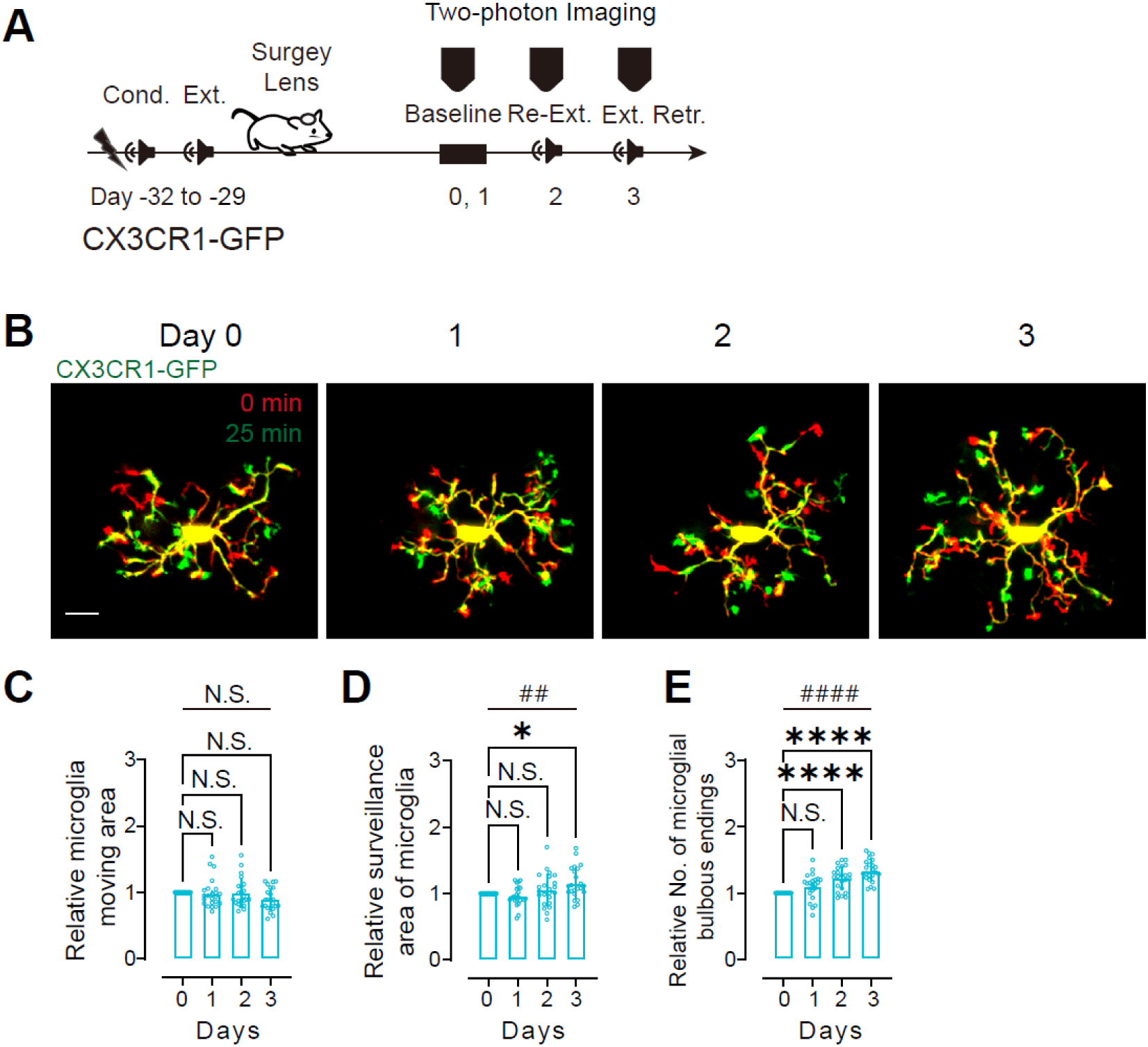
Longitudinal *in vivo* imaging reveals extinction retrieval–evoked expansion of microglial surveillance territories, related to Figure 2. (A) Two-photon imaging schedule in CX3CR1-GFP mice. Animals underwent auditory fear conditioning (Cond.), and extinction training (Ext.), followed by GRIN lens implantation over mPFC. The same fields were imaged at baseline (Days 0–1), during re-extinction (Re-Ext., Day 2), and at extinction retrieval (Ext. Retr., Day 3). (B) Representative microglial cell imaged on each day. Overlays show the same cell at 0 min (red) and 25 min (green) within a session; overlap appears yellow. Scale bar, 10 μm. (C–E) Quantification of within-session process dynamics from paired images (0 and 25 min), expressed relative to Day 0 for each cell. (C) Moving area, area swept by motile processes. (D) Surveillance area, total territory explored by processes over the imaging interval. (E) Bulbous endings, number of distal bulbous process termini. Moving area remains stable across days, whereas surveillance area increases selectively at extinction retrieval and bulbous endings are elevated during re-extinction and retrieval. *n* = 23 cells (4 mice). Bars, mean ± s.e.m.; dots, individual cells. Statistics: one-way repeated-measures ANOVA with Dunnett’s multiple comparisons versus Day 0 and planned paired comparisons between days, as indicated. N.S., not significant; \**P* < 0.05, \*\*\*\**P* < 0.0001, paired Student’s *t* test. ^##^*P* < 0.01, ^####^*P* < 0.0001, one-way ANOVA with Dunnett’s multiple-comparisons test.

**Figure S5.**
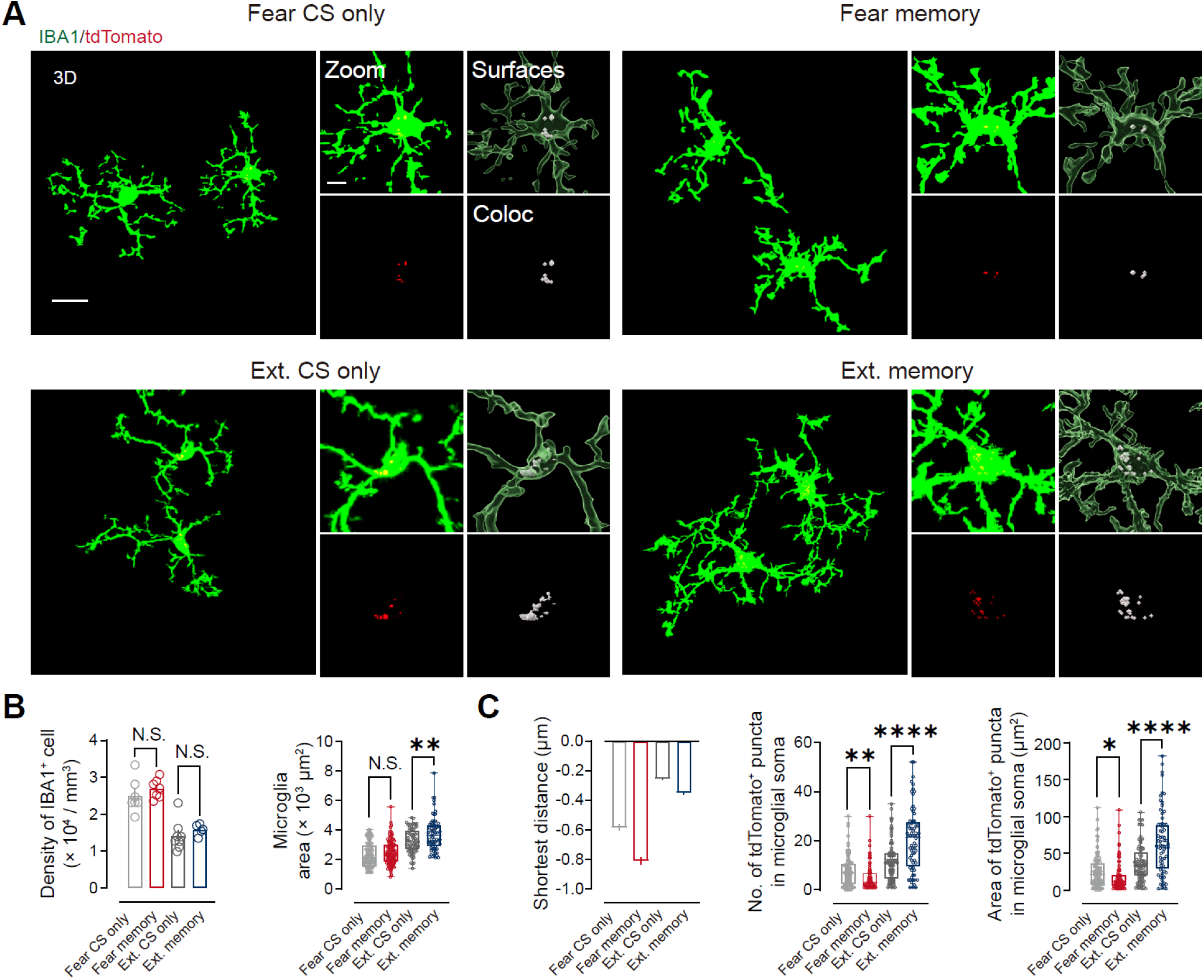
Extinction retrieval expands microglial arbors and increases internalized ensemble-derived puncta in microglial somata, related to Figure 3. (A) Representative 3D reconstructions from mPFC of FosTRAP2::Ai14 mice immunolabeled for IBA1 (microglia, green) with tdTomato (tdTomato^+^ ensemble signal, red) under the indicated conditions (Fear CS only, Fear memory, Ext. CS only, Ext. memory). For each group, right panels show a higher-magnification view (Zoom), 3D surface rendering of the microglial soma/process compartment (Surfaces), and a colocalization map (Coloc) highlighting tdTomato^+^ puncta assigned to the microglial soma volume. Scale bars, 15 μm (3D) and 5 μm (Zoom). (B) Microglial abundance and morphology quantified from IBA1 signal. Left, density of IBA1^+^ cells (×10^4^ cells/mm^3^) is unchanged between each memory group and its CS-only control. Right, microglial area per cell (IBA1-based surface area; ×10^3^ µm^2^) increases selectively in Ext. memory relative to Ext. CS only, with no increase in the fear comparison. Density: Fear CS only, *n* = 7 mice; Fear memory, n = 7 mice; Ext. CS only, *n* = 8 mice; Ext. memory, *n* = 5 mice. Area: Fear CS only, *n* = 110 cells (7 mice); Fear memory, *n* = 112 cells (7 mice); Ext. CS only, *n* = 67 cells (8 mice); Ext. memory, *n* = 68 cells (5 mice). (C) Ensemble-derived puncta within microglial somata. Left, signed shortest distance from tdTomato^+^ puncta to the microglial soma surface (negative values indicate puncta located within the soma volume, consistent with internalization). Middle and right, number and summed area of tdTomato^+^ puncta within microglial somata. Internalized tdTomato^+^ puncta decrease after fear retrieval relative to Fear CS only, but increase after extinction retrieval relative to Ext. CS only. Fear CS only, *n* = 105 cells (7 mice); Fear memory, *n* = 78 cells (7 mice); Ext. CS only, *n* = 84 cells (8 mice); Ext. memory, *n* = 68 cells (5 mice). N.S., not significant; \**P* < 0.05, \*\**P* < 0.01, \*\*\*\**P* < 0.0001, comparisons are between each memory group and its corresponding CS-only control, two-sided unpaired Student’s *t* test.

**Figure S6.**
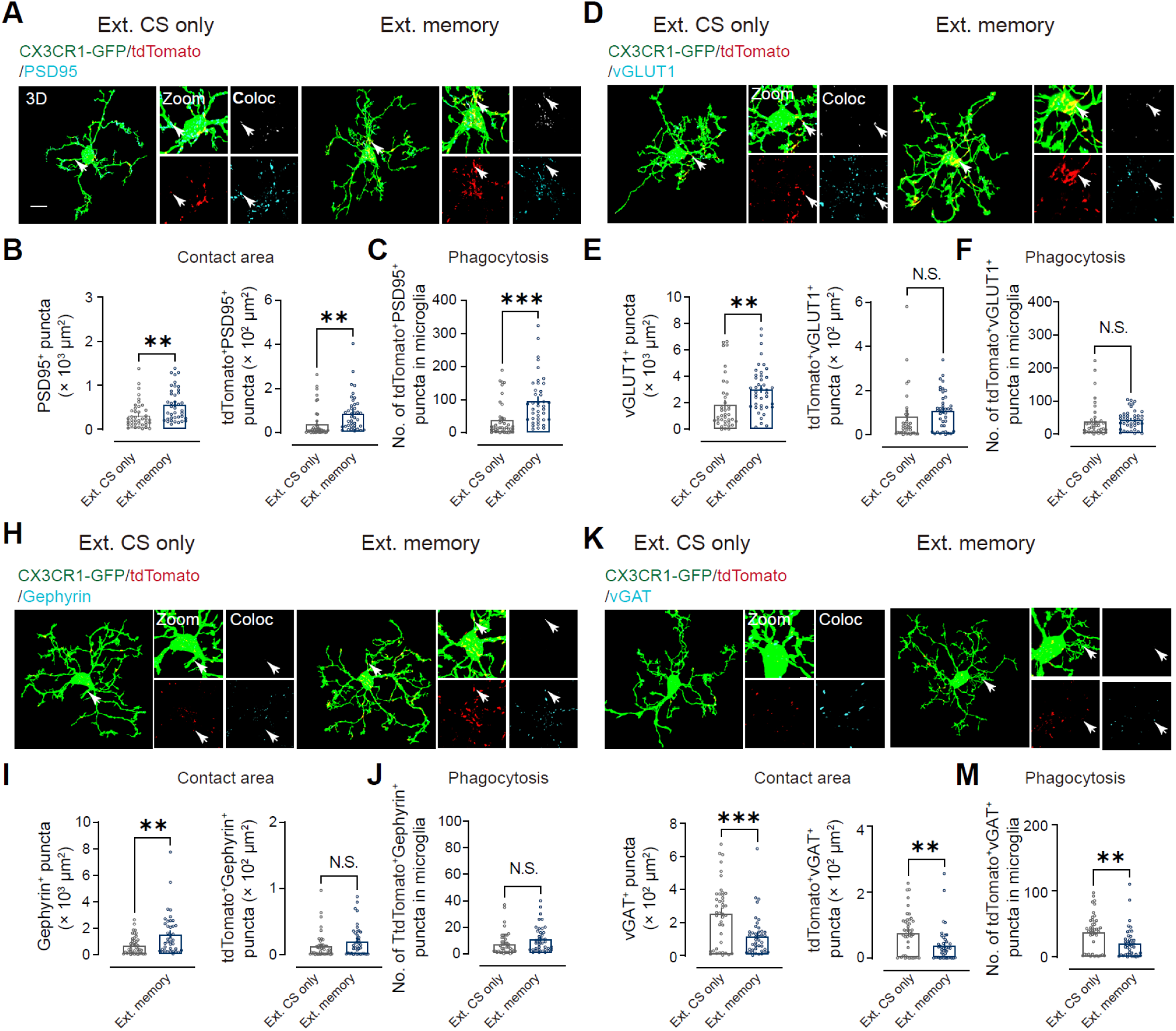
Extinction retrieval bias microglial engulfment toward ensemble-derived excitatory postsynaptic sites, related to Figure 3. (A) Representative 3D confocal reconstructions from mPFC of FosTRAP2::Ai14;;CX3CR1-GFP mice showing microglia (CX3CR1-GFP, green), extinction-tagged ensemble signal (tdTomato, red), and PSD-95 (cyan) in Ext. CS only and Ext memory (extinction retrieval) groups. Insets show a higher-magnification view (Zoom) and a colocalization map (Coloc) highlighting tdTomato^+^ synaptic puncta assigned to microglial contact/engulfment (arrows). Scale bars, 10 μm (3D) and 2 μm (zoom). (B and C) Quantification of PSD-95 engagement. (B) Contact area between microglia and PSD95^+^ puncta (left) and the subset attributable to tdTomato^+^PSD-95^+^ puncta (right). (C) Engulfment quantified as the number of tdTomato^+^PSD-95^+^ puncta enclosed within microglial volumes per cell. Ext. CS only, *n* = 40 cells (7 mice); Ext. memory, *n* = 40 cells (7 mice). (D) As in (A), but for vGLUT1 (cyan). (E and F) vGLUT1 engagement. (E) Contact area with all vGLUT1^+^ puncta (left) and with tdTomato^+^vGLUT1^+^ puncta (right). (F) Number of engulfed tdTomato^+^vGLUT1^+^ puncta per microglia. Ext. CS only, *n* = 38 microglia (7 mice); Ext. memory, *n* = 41 microglia (7 mice). (H) As in (A), but for Gephyrin (cyan). (I and J) Gephyrin engagement. (I) Contact area with all Gephyrin^+^ puncta (left) and with tdTomato^+^Gephyrin^+^ puncta (right). (J) Number of engulfed tdTomato^+^Gephyrin^+^ puncta per microglia. Ext. CS only, *n* = 42 microglia (7 mice); Ext. memory, *n* = 42 microglia (7 mice). (K) As in (A), but for vGAT (cyan). (L and M) vGAT engagement. (L) Contact area with all vGAT^+^ puncta (left) and with tdTomato^+^vGAT^+^puncta (right). (M) Number of engulfed tdTomato^+^vGAT^+^ puncta per microglia. Ext. CS only, *n* = 42 microglia (7 mice); Ext. memory, *n* = 42 microglia (7 mice). Across markers, extinction retrieval increased contact with—and engulfment of—tdTomato^+^PSD-95^+^ postsynaptic puncta, did not increase engulfment of tdTomato^+^vGLUT1^+^ or tdTomato^+^Gephyrin^+^ puncta, and reduced both contact and engulfment of tdTomato^+^VGAT^+^ puncta. N.S., not significant; \*\**P* < 0.01, \*\*\**P* < 0.001, two-sided unpaired Student’s *t* test.

**Figure S7.**
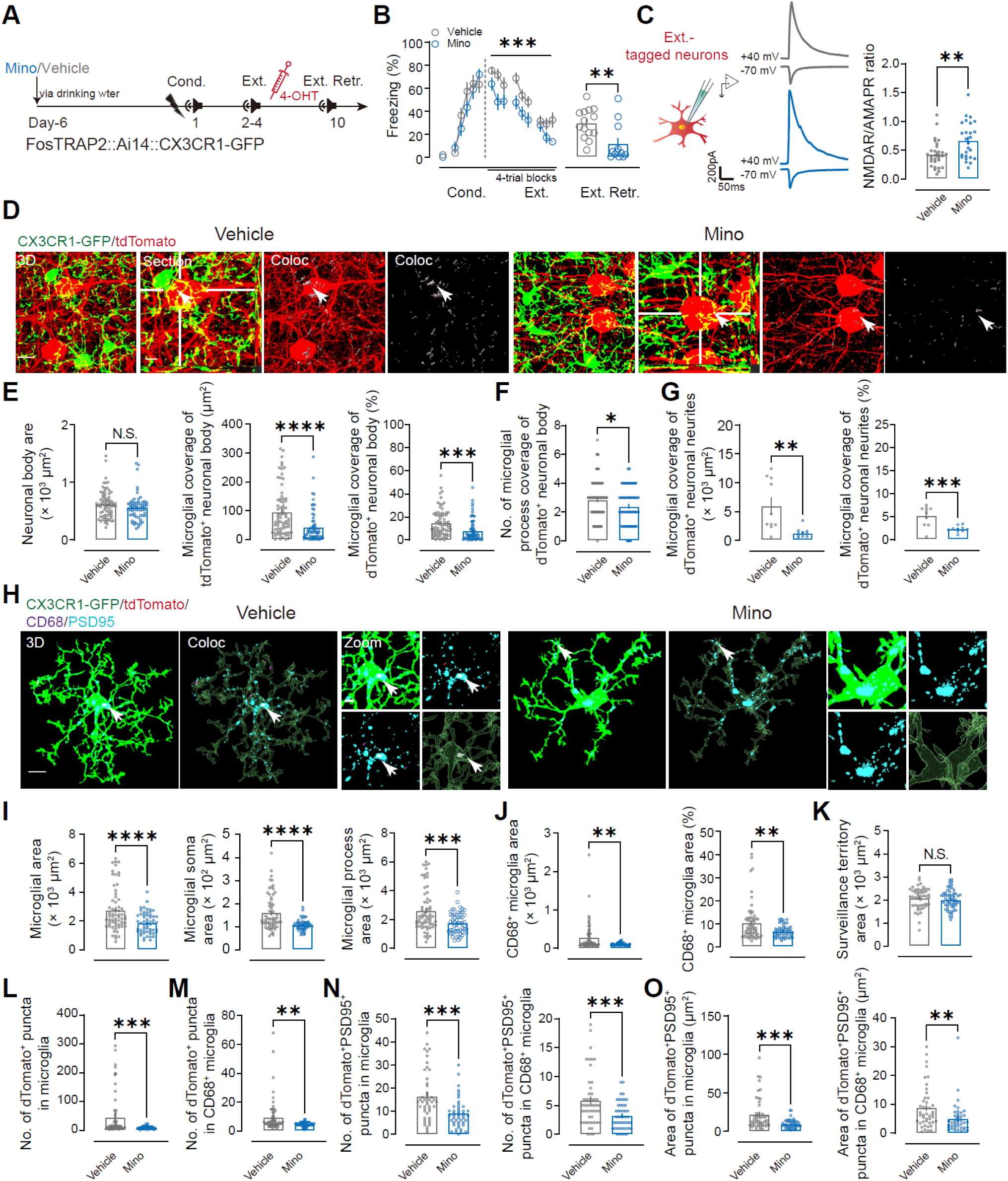
Minocycline facilitates fear extinction while dampening microglial engagement and postsynaptic engulfment at extinction ensembles, related to Figure 3. (A) Experimental timeline in FosTRAP2::Ai14;;CX3CR1-GFP mice. Minocycline (Mino) or vehicle was provided in drinking water starting from day −6. Mice underwent fear conditioning (Cond., day 1) followed by extinction training (Ext., days 2–4); 4-OHT was administered after the third extinction session to TRAP-label extinction-ensemble neurons. Extinction retrieval (Ext. Retr.) was tested on day 10. (B) Freezing during Cond., Ext., and Ext. Retr. Minocycline spared fear acquisition but accelerated extinction and reduced freezing at retrieval. Vehicle, *n* = 14 mice; Mino, *n* = 12 mice. (C) Synaptic physiology in TRAP-labeled extinction-ensemble neurons after Ext. Retr. Left, schematic and representative traces; right, quantification of the NMDAR/AMPAR eEPSC ratio. Vehicle, *n* = 30 cells (6 mice); Mino, *n* = 26 cells (6 mice). (D) Representative confocal images showing CX3CR1-GFP^+^ microglia (green) and tdTomato^+^ extinction-ensemble neurons (red) after Ext. Retr. *Coloc* indicates microglia–ensemble apposition (arrows). Scale bar, 10 μm. (E to G) Minocycline reduces structural coupling between microglia and extinction ensembles. (E) Ensemble body (left), microglial coverage of ensemble body (middle; absolute area); and somatic coverage expressed as a percentage (right). (F) Number of microglial processes contacting each ensemble body. (G) Microglial coverage of ensemble neurites (left; absolute area) and neurite coverage as a percentage (right). Vehicle, *n* = 66 cells from 10 sections (5 mice); Mino, *n* = 74 cells from 10 sections (5 mice). (H) Representative 3D renderings of microglia (green) with lysosomes (CD68, cyan) and postsynaptic puncta (PSD-95, magenta) illustrating reduced microglial lysosomal loading and reduced tdTomato^+^PSD-95^+^ cargo after minocycline. Insets show zoom views; arrows indicate engulfed puncta. Scale bars, 7 μm (overview), 2 μm (zoom). (I–K) Minocycline blunts retrieval-associated microglial state changes. (I) Total microglial area (left), soma area (middle), and process area (right). (J) CD68^+^ lysosomal area per microglia (left) and as a fraction of microglial area (right). (K) Surveillance territory per microglia. Vehicle, *n* = 64 cells (5 mice); Mino, *n* = 55 cells (5 mice); surveillance territory: vehicle, *n* = 48 cells (5 mice); Mino, *n* = 59 cells (5 mice). (L–O) Minocycline suppresses internalization of extinction-ensemble material and excitatory postsynaptic cargo. (L) Total tdTomato^+^ puncta per microglia. (M) tdTomato^+^ puncta within CD68^+^ compartments. (N) tdTomato^+^PSD-95^+^ puncta per microglia (left) and within CD68^+^ compartments (right). Vehicle, *n* = 48 cells (5 mice); Mino, *n* = 47 cells (5 mice). Data are mean ± s.e.m. (B and C) or shown as individual cells with summary statistics as indicated. N.S., not significant; \**P* < 0.05, \*\**P* < 0.01, \*\*\**P* < 0.001, \*\*\*\**P* < 0.0001, unpaired Student’s *t* test.

**Figure S8.**
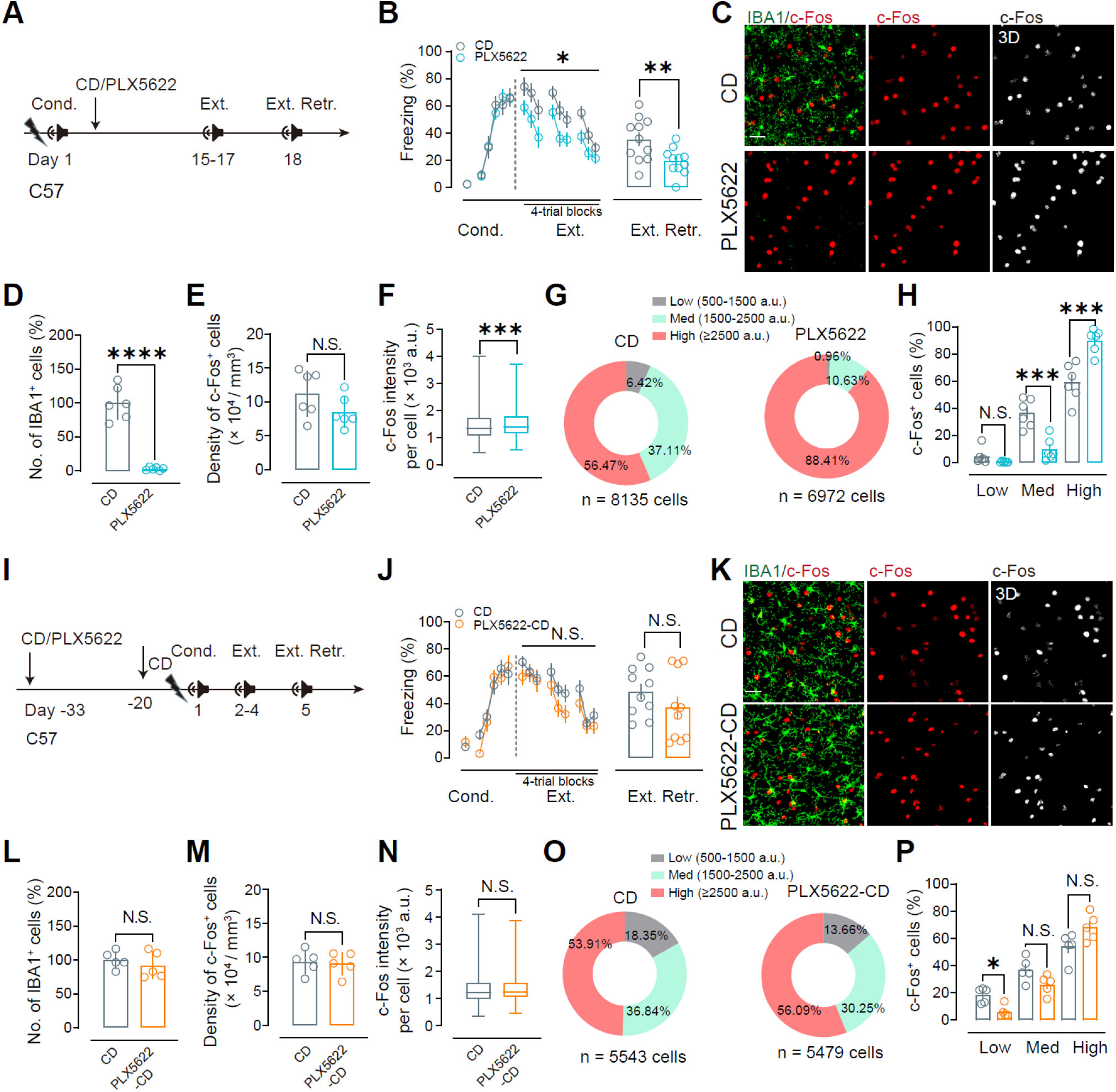
Microglial depletion restricted to the extinction phase facilitates extinction and elevates retrieval-time neuronal activation, and these effects normalize after microglial repopulation, related to Figure 4. (A) Experimental timeline for extinction-phase depletion. C57BL/6J mice were maintained on control diet (CD) or PLX5622 beginning after conditioning (Cond.; Day 1) and throughout extinction training (Ext.; Days 15–17) and extinction retrieval (Ext. Retr.; Day 1). (B) Freezing during CS presentations across Cond., Ext., and Ext. Retr. PLX5622 treatment accelerated extinction and reduces freezing at Ext. Retr. (mean ± s.e.m.). CD, *n* = 11 mice; PLX5622, *n* = 11 mice. (C) Representative mPFC sections at Ext. Retr. immunolabeled for microglia (IBA1, green) and activity (c-Fos, red); right panels show 3D segmentation of c-Fos signal used for per-cell quantification. Scale bar, 50 μm. (D to H) Quantification of depletion and retrieval-time activation in mPFC at Ext. Retr. (D) IBA1^+^ cell counts (expressed as % of CD) confirm robust depletion. (E) Density of c-Fos^+^ cells. (F) Per-cell c-Fos fluorescence intensity. (G) Distribution of c-Fos^+^ cells across intensity bins (Low, 500–1500 a.u.; Med, 1500–2500 a.u.; High, ≥2500a.u.). CD, *n* = 8135 c-Fos^+^ cells; PLX5622, *n* = 6972 c-Fos^+^ cells; pooled from 6 mice per group. (H) Fraction of c-Fos^+^ cells in each intensity bin, showing a depletion-associated shift toward higher-intensity activation. (I) Repopulation design. Mice were fed PLX5622 from Day –33 to –20 and then returned to CD (PLX5622→CD) to allow microglial repopulation before Cond. (Day 1), Ext. (Days 2–4), and Ext. Retr. (Day 5). (J) Freezing across Cond., Ext., and Ext. Retr. after repopulation, showing no detectable differences between CD and PLX5622→CD groups (mean ± s.e.m.). CD, *n* = 10 mice; PLX5622→CD, *n* = 10 mice. (K) Representative mPFC sections after repopulation, stained as in (C). Scale bar, 50 μm. (L to P) Quantification after repopulation. (L) IBA1^+^ cell counts are restored. (M) Density of c-Fos^+^ cells. (N) Per-cell c-Fos intensity. (O) Distribution of c-Fos⁺ cells across intensity bins. CD, *n* = 5543 c-Fos^+^ cells; PLX5622→CD, *n* = 5479 c-Fos^+^ cells; pooled from 5 mice per group. (P) Fraction of c-Fos^+^ cells in each bin, showing overall normalization of the activation profile (with high-intensity fractions indistinguishable from controls). Statistics: two-way repeated-measures ANOVA (behavioral time courses) with post hoc tests as indicated; unpaired two-tailed Student’s *t* tests for between-group comparisons at retrieval and for image-based quantifications. N.S., not significant; \**P* < 0.05, \*\**P* < 0.01, \*\*\**P* < 0.001, \*\*\*\**P* < 0.0001.

**Figure S9.**
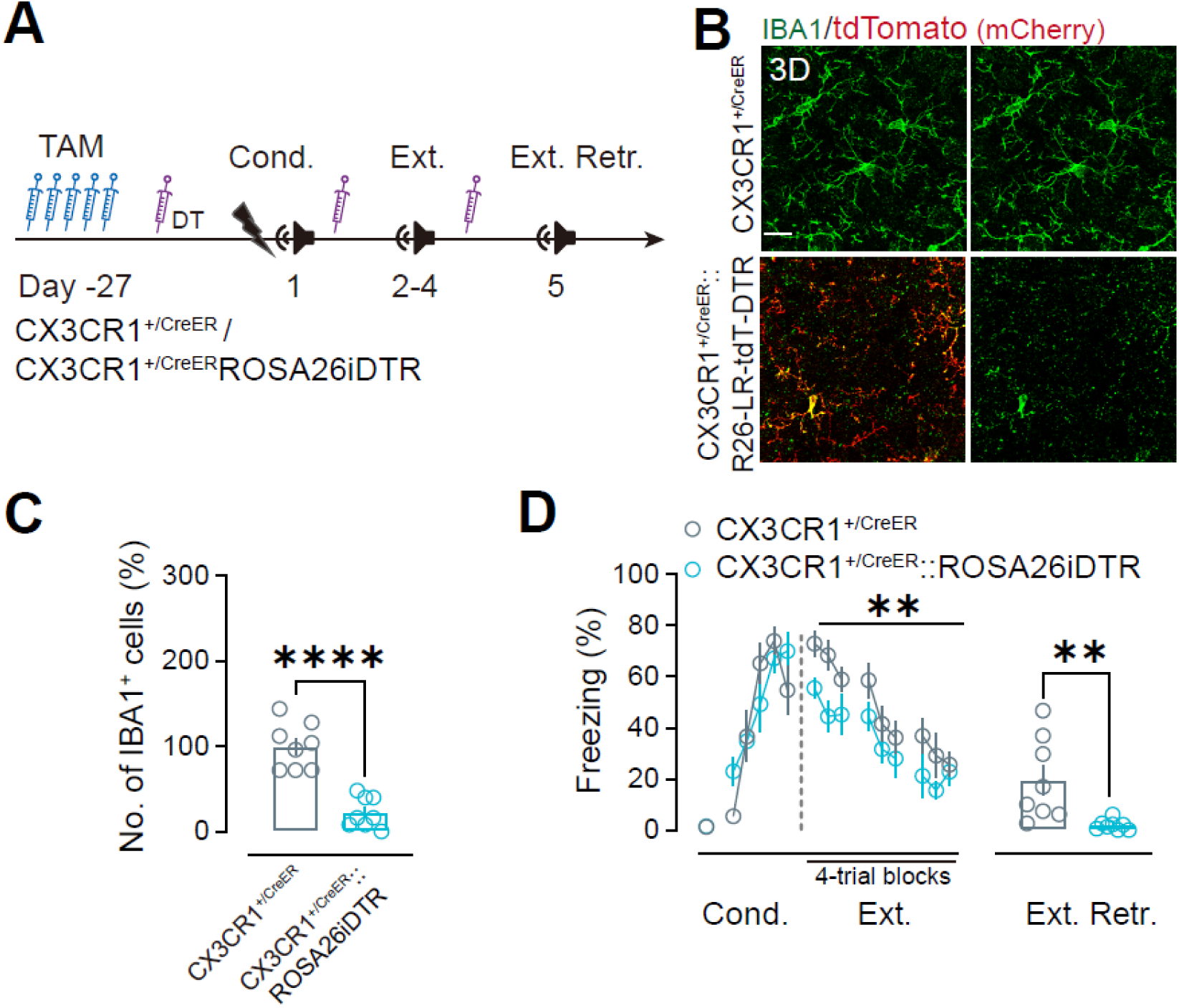
Inducible DTR-mediated microglial ablation facilitates fear extinction, related to Figure 4. (A) Experimental design in CX3CR1^+/CreER^ control and CX3CR1^+/CreER^::ROSA26iDTR mice. Tamoxifen (TAM) was administered on day −27 to induce tdTomato/DTR expression in CX3CR1^+^ microglia, and diphtheria toxin (DT) was delivered at the indicated time points during conditioning (Cond.), extinction training (Ext.), and extinction retrieval (Ext. Retr.) to ablate DTR-expressing cells. (B) Representative mPFC images at extinction retrieval showing IBA1 (green) and tdTomato/DTR reporter signal (red), illustrating marked loss of IBA1^+^ microglia in CX3CR1^+/CreER^::ROSA26iDTR mice. Scale bar, 15 μm. (C) Quantification of IBA1^+^ microglia (control set to 100%), confirming efficient ablation. (D) Freezing during conditioning, extinction training (4-trial blocks), and extinction retrieval. Microglial ablation leaves fear acquisition intact but accelerates extinction and reduces freezing at extinction retrieval. *n* = 8 mice per group. Data are mean ± s.e.m. \*\**P* < 0.01, \*\*\*\**P* < 0.0001. Statistics: two-tailed unpaired Student’s *t* test for (C) and retrieval comparison in (D); two-way repeated-measures ANOVA for the extinction time course in (D), with post hoc testing as indicated in the panel and described in Methods.

**Figure S10.**
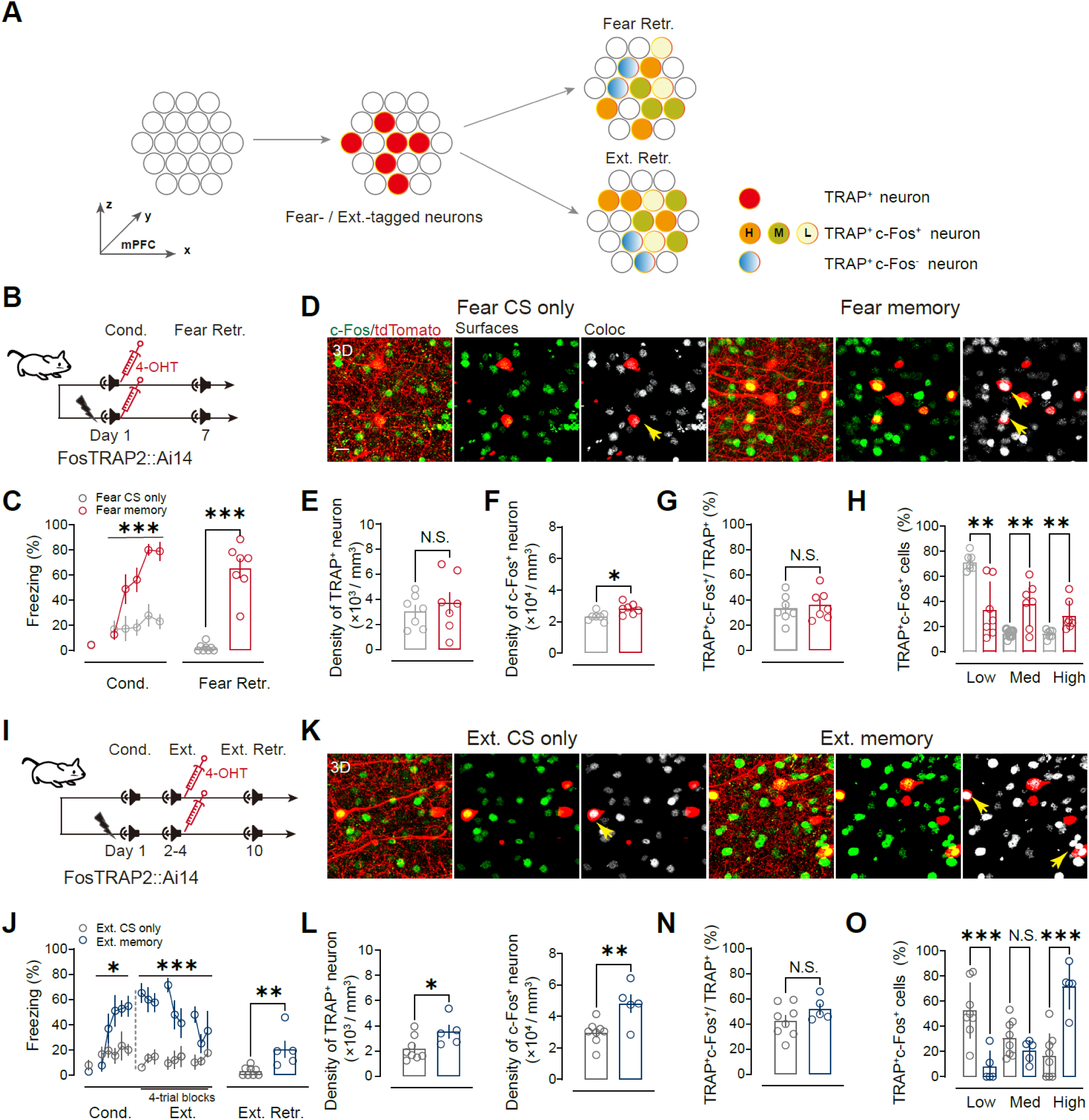
Highly active fear and extinction ensemble neurons are preferentially reactivated at retrieval, related to Figure 4. (A) Conceptual schematic of activity-dependent ensemble tagging and retrieval-time reactivation analysis in mPFC. Neurons tagged by FosTRAP (TRAP^+^; tdTomato^+^) during learning are reassessed at retrieval by c-Fos immunostaining; tdTomato^+^c-Fos^+^ neurons are further binned by c-Fos intensity (Low/Med/High). (B) Fear-ensemble tagging and retrieval schedule in FosTRAP2::Ai14 mice. 4-OHT was administered immediately after fear conditioning (Cond., Day 1) to label fear-learning–activated neurons, followed by fear retrieval (Fear Retr., Day 7). (C) Freezing during fear conditioning and retrieval in Fear CS-only and Fear memory groups (mean ± s.e.m.). *n* = 7 mice per group. (D) Representative confocal images of mPFC showing tdTomato^+^ neurons (tdTomato, red), retrieval-induced c-Fos neurons (green), segmented *Surfaces*, and tdTomato^+^c-Fos^+^ colocalization (*Coloc*, white). Arrowheads indicate reactivated TRAP⁺c-Fos⁺ cells. Scale bar, 20 µm. (E to H) Quantification of fear-ensemble recruitment at retrieval. (E) Density of tdTomato^+^ neurons. (F) Density of c-Fos^+^ neurons. (G) Reactivation fraction (tdTomato^+^c-Fos^+^ / tdTomato^+^, %). (H) Distribution of tdTomato^+^c-Fos^+^ neurons across c-Fos intensity bins (Low/Med/High), showing a retrieval-associated shift toward higher-intensity reactivated ensemble cells in the fear-memory group. (I) Extinction-ensemble tagging and retrieval schedule. Mice were conditioned (Day 1), underwent extinction training (Ext., days 2–4), received 4-OHT after the final extinction session to label extinction-learning–activated neurons, and were tested at extinction retrieval (Ext. Retr., Day 10). (J) Freezing during extinction training and extinction retrieval in Ext. CS-only and Ext. memory groups (mean ± s.e.m.). Ext. CS-only, *n* = 8 mice; Ext. memory, *n* = 5 mice. (K) Representative confocal images and segmentation as in (D) for extinction retrieval. Arrowheads indicate tdTomato^+^c-Fos^+^ cells. Scale bar, 20 µm. (L to O) Quantification of extinction-ensemble recruitment at retrieval. (L) Density of tdTomato^+^ neurons. (M) Density of c-Fos^+^ neurons. (N) Reactivation fraction (tdTomato^+^c-Fos⁺ / tdTomato⁺, %). (O) Distribution of tdTomato^+^c-Fos^+^ neurons across c-Fos intensity bins, indicating preferential engagement of the high-activity subset during extinction retrieval. Statistics: two-way repeated-measures ANOVA for freezing across sessions with post hoc comparisons; unpaired two-tailed Student’s t tests for between-group comparisons, as appropriate. N.S., not significant; \**P* < 0.05, \*\**P* < 0.01, \*\*\**P* < 0.001.

**Figure S11.**
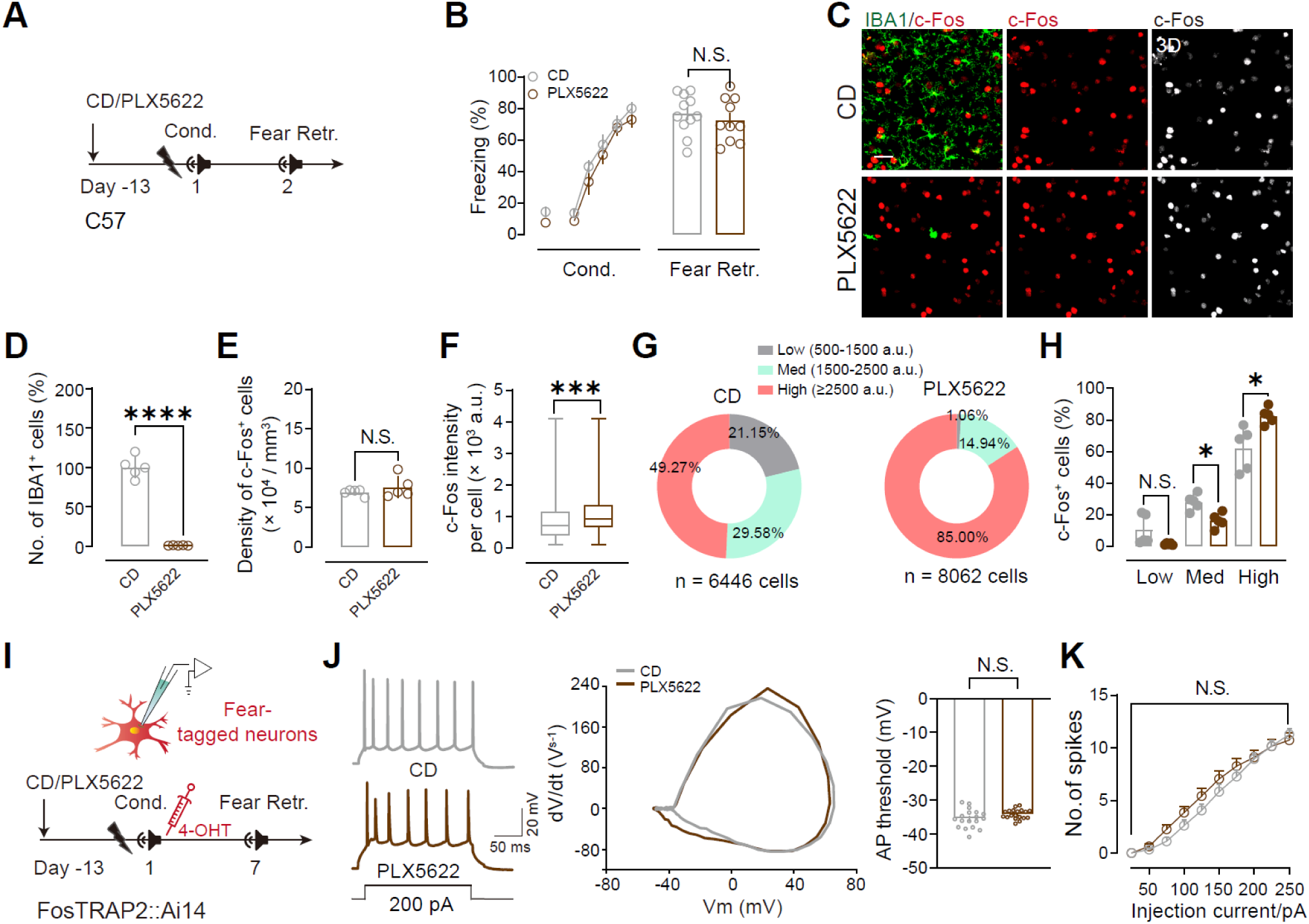
Microglial depletion spars fear acquisition and retrieval and does not increase excitability of fear ensemble neurons, related to Figure 4. (A) Timeline for diet-based microglial depletion. C57BL/6J mice were maintained on control diet (CD) or PLX5622 chow (CSF1R inhibitor) beginning 13 days before auditory fear conditioning (Cond.) and tested for fear retrieval (Fear Retr.) 24 h later. (B) Freezing during conditioning and at fear retrieval (mean ± s.e.m.). *n* = 11 mice per group. (C) Representative mPFC sections collected after fear retrieval, immunolabeled for IBA1 (green) and c-Fos (red); right, 3D rendering of the c-Fos signal. Scale bar, 50 μm. (D–H) Quantification of depletion and retrieval -evoked activation in mPFC. (D) IBA1^+^ microglia remaining (percent of CD). (E) Density of c-Fos⁺ cells. (F) Per-cell c-Fos fluorescence intensity. (G) Distribution of c-Fos^+^ cells across intensity bins (Low, 500–1500 a.u.; Med, 1500–2500; High, ≥2500 a.u.). CD, *n* = 6466 cells; PLX5622, *n* = 8062 cells; 5 mice per group. (H) Summary of the fraction of c-Fos^+^ cells in each bin, showing a shift toward Fos-high neurons with microglial depletion. (I) Timeline for TRAP labeling and *ex vivo* recording. FosTRAP2::Ai14 mice were fed CD or PLX5622 from Day –13; 4-OHT was administered immediately after conditioning to tag fear ensembles, and tdTomato^+^ neurons were targeted for whole-cell current-clamp recording in acute mPFC slices prepared after fear retrieval (Day 7). (J) Representative current-evoked firing (200 pA), phase plots, and action-potential threshold measurements from fear-tagged neurons in CD and PLX5622 groups. (K) Frequency–current relationship (spike number versus injected current) for fear-tagged neurons. CD, *n* = 13 neurons from 6 mice; PLX5622, *n* = 13 neurons from 6 mice. Data are mean ± s.e.m. N.S., not significant; \**P* < 0.05, \*\*\**P* < 0.001, two-way repeated-measures ANOVA or unpaired two-tailed Student’s *t* test, as appropriate.

**Figure S12.**
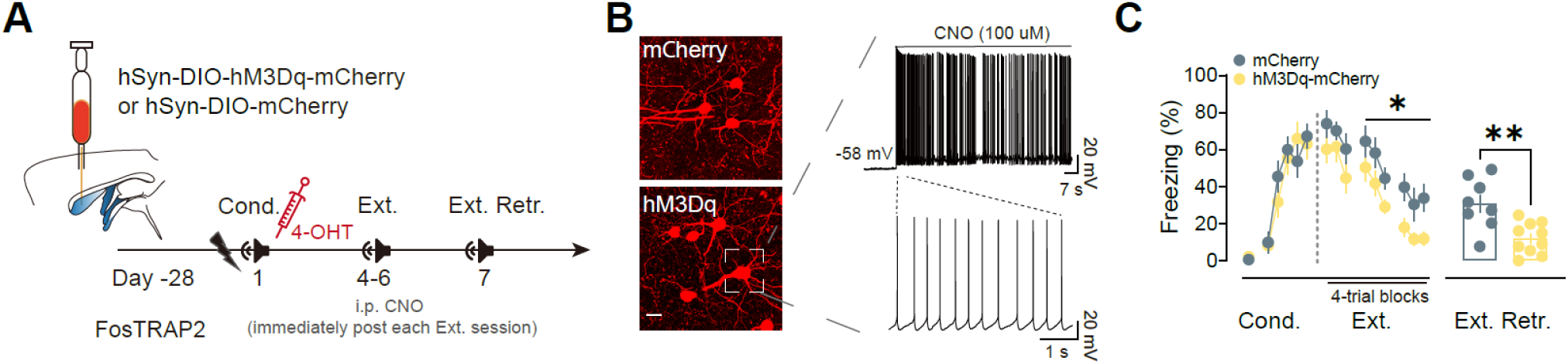
Chemogenetic activation of fear-ensemble neurons facilitates extinction, related to Figure 4. (A) Experimental design. FosTRAP2 mice received intracerebroventricular (ICV) injection of AAV-hSyn-DIO-hM3Dq-mCherry or control AAV-hSyn-DIO-mCherry (Day –28). Fear ensemble neurons were TRAP-labeled by 4-OHT immediately after conditioning (Cond., Day 1), restricting hM3Dq-mCherry or mCherry expression to conditioning-activated neurons. Mice then underwent extinction training (Ext., Days 4–6); clozapine-N-oxide (CNO) was delivered intraperitoneally immediately after each extinction session to engage hM3Dq during the post-session window, followed by an extinction retrieval test (Ext. Retr., Day 7). (B) Chemogenetic validation. Representative mPFC images showing mCherry control expression (top) or hM3Dq-mCherry expression (bottom). Right, representative whole-cell current-clamp recordings from an hM3Dq^+^ neuron showing robust depolarization and high-frequency firing in response to bath-applied CNO (100 μM). Scale bar, 20 μm. (C) Behavioral effects. Freezing during CS presentations across Cond., Ext., and Ext. Retr. in mCherry controls (*n* = 8 mice) and hM3Dq-mCherry mice (*n* = 10 mice). Post-session chemogenetic activation of fear-ensemble neurons accelerated extinction learning and reduced freezing at extinction retrieval. Data are mean ± s.e.m. \**P* < 0.05, \*\**P* < 0.01, two-way repeated-measures ANOVA for Ext. and unpaired Student’s *t* test at Ext. Retr.

**Figure S13.**
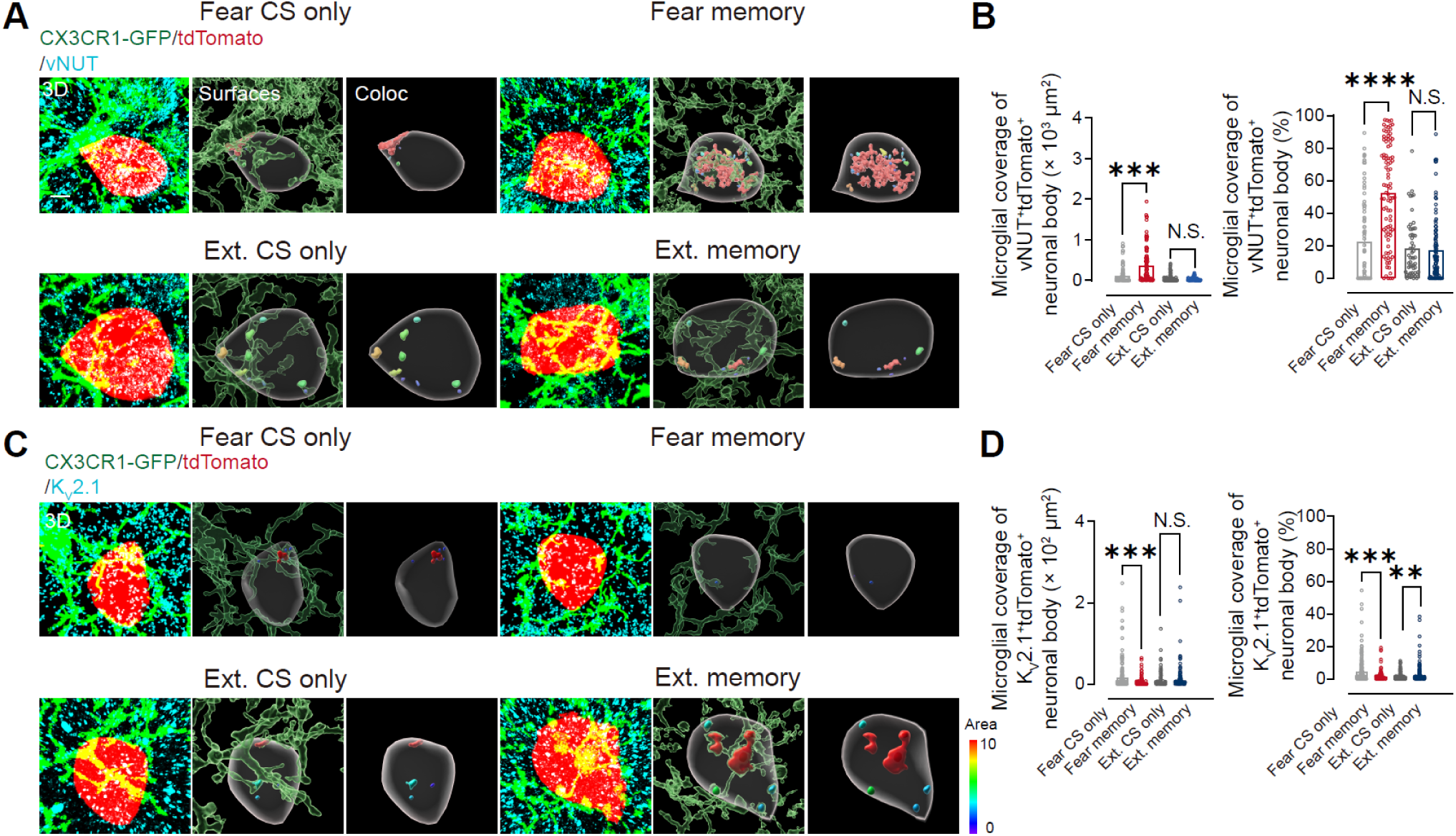
vNUT and K_V_2.1 are differentially enriched at microglia–ensemble somatic junctions during fear and extinction retrieval, related to Figure 5. (A) Representative 3D reconstructions of FosTRAP2::Ai14;;CX3CR1-GFP mice immunolabeled for vesicular nucleotide transporter vNUT (cyan). Microglia (CX3CR1-GFP, green) contact TRAP-labeled ensemble bodies (tdTomato, red). *Surfaces*, 3D surface renderings of microglia and TRAP⁺ somata; *Coloc*, vNUT signal restricted to the microglia-apposed somatic surface (puncta color-coded by area as indicated). Scale bar, 3 μm. (B) Microglia coverage of vNUT^+^ somatic microdomains on tdTomato^+^ ensemble neurons, quantified as absolute coverage area (left) and fractional coverage (right). Fear retrieval increases microglial apposition to vNUT^+^ domains, whereas extinction retrieval does not. Fear CS only, *n* = 82 cells (6 mice); fear memory, *n* = 91 cells (6 mice); Ext. CS only, *n* = 54 cells (7 mice); Ext. memory, *n* = 102 cells (7 mice). (C) As in (A), but immunolabeled for K_V_2.1 (cyan). Scale bar, 3 μm. (D) Microglial coverage of K_V_2.1^+^ somatic microdomains on tdTomato^+^ ensemble neurons (absolute coverage area, left; fractional coverage, right). Fear retrieval reduces microglial engagement of Kv2.1⁺ domains, whereas extinction retrieval selectively increases fractional coverage (with no significant change in absolute covered area). Fear CS only, *n* = 203 cells (7 mice); fear memory, *n* = 111 cells (7 mice); Ext. CS only, *n* = 238 cells (7 mice); Ext. memory, *n* = 224 cells (7 mice). N.S., not significant; \*\**P* < 0.01, \*\*\**P* < 0.001, \*\*\*\**P* < 0.0001, unpaired Student’s *t* test (memory versus corresponding CS-only control).

**Figure S14.**
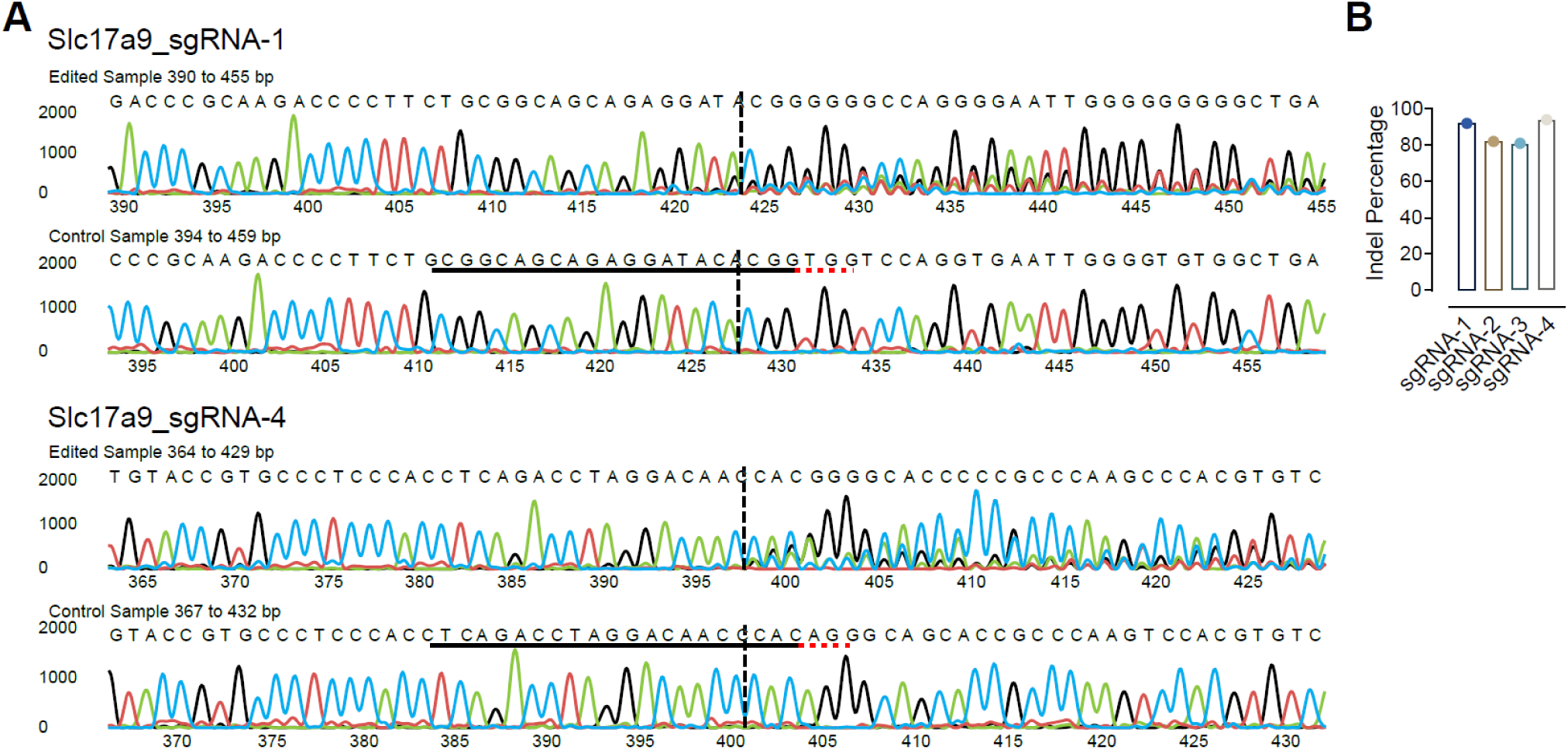
Sanger sequencing validates CRISPR–Cas9 editing of *Slc17a9* (vNUT), related to Figure 5. (A) Representative Sanger chromatograms of PCR amplicons spanning the *Slc17a9* sgRNA-1 (top) and sgRNA-4 (bottom) target sites from edited and control samples. The predicted Cas9 cleavage site is indicated (dashed line). Edited samples show mixed base calls downstream of the cut site, consistent with indel generation; sgRNA target sequences are underlined. (B) Indel frequencies for four *Slc17a9* sgRNAs (sgRNA-1 to sgRNA-4), estimated from Sanger trace decomposition.

**Figure S15.**
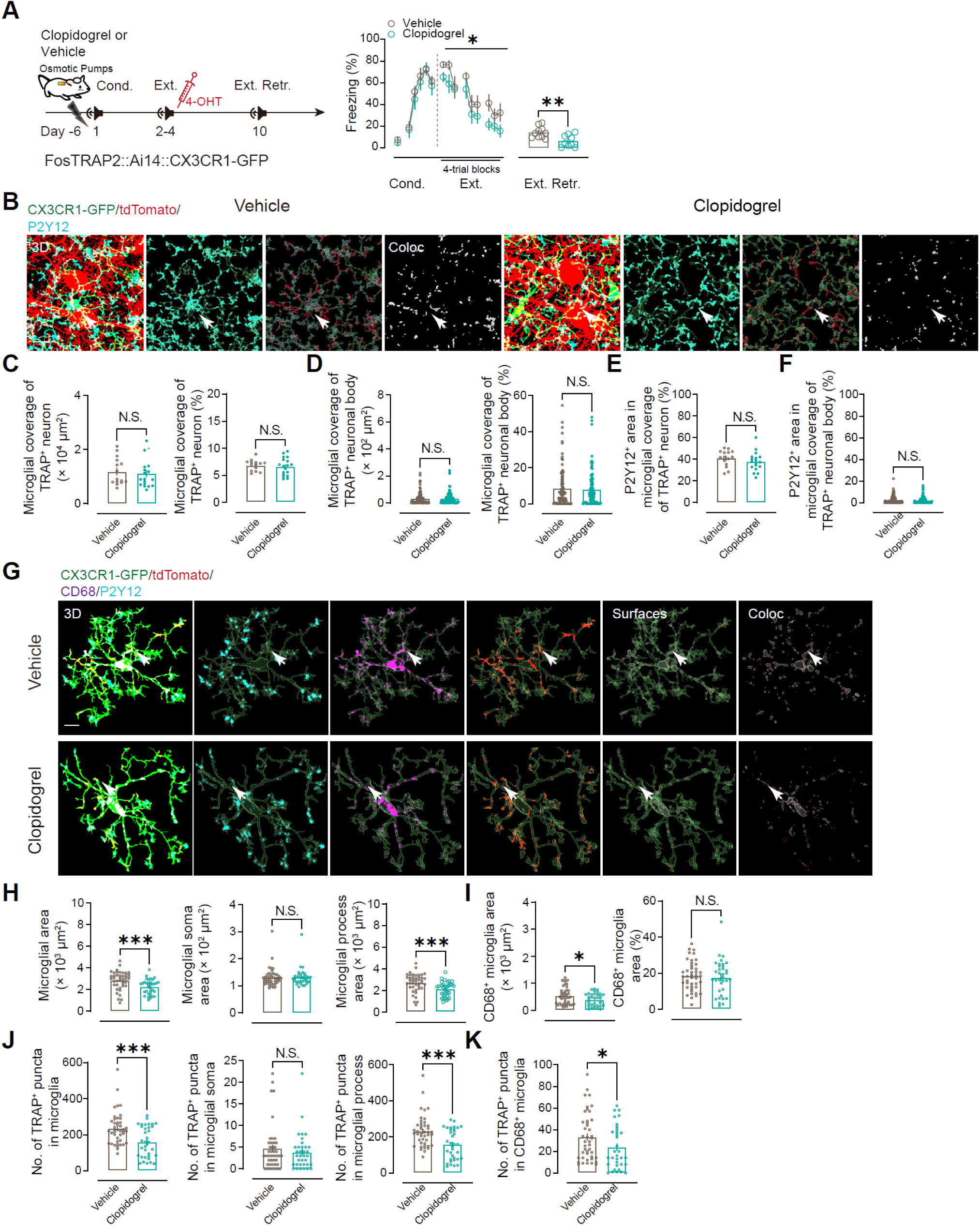
P2Y12 antagonism promotes extinction and selectively reduces process-associated microglial engulfment without altering somatic apposition to extinction ensembles, related to Figure 5. (A) Continuous intracerebroventricular (ICV) infusion of the P2Y12 antagonist clopidogrel or vehicle control via osmotic minipumps in FosTRAP2::Ai14;;CX3CR1-GFP mice (pump implantation, Day – 6; conditioning, Cond., Day 1; extinction training, Ext., Days 2–4, with 4-OHT after the third session; extinction retrieval, Ext. Retr., Day 10). Right, freezing during conditioning, extinction (4-trial blocks), and extinction retrieval (mean ± s.e.m.). Vehicle, *n* = 10 mice; clopidogrel, *n* = 9 mice. (B) Representative mPFC reconstructions at extinction retrieval showing CX3CR1-GFP^+^ microglia (green), extinction-tagged neurons (tdTomato, red), and P2Y12 (cyan). *Coloc*, P2Y12 signal within the microglia–TRAP contact shell (arrows). Scale bar, 10 μm. (C and D) Gross microglia–ensemble apposition is preserved after clopidogrel treatment. Microglial coverage of tdTomato^+^ neurons (C; tdTomato^+^ surface) and tdTomato^+^ bodies (D) quantified as absolute contact area (left) and percent surface coverage (right). (E and F) P2Y12^+^ area within microglia-apposed tdTomato^+^ surfaces (E) and within microglia-apposed tdTomato^+^ neuronal bodies (F). Vehicle, *n* = 99 cells from 15 sections (5 mice); clopidogrel, *n* = 96 cells from 18 sections (5 mice). (G) Representative single-cell 3D reconstructions illustrating microglial morphology and intracellular cargo after vehicle- and clopidogrel. Channels show GFP (microglia, green), P2Y12 (cyan), CD68 (magenta), and TRAP (red). *Coloc*, tdTomato^+^ puncta within microglia (arrows). Scale bar, 10 μm. (H and I) Microglial state metrics. Total area, soma area, and process area (H), and CD68⁺ lysosomal area per microglia and fraction of CD68^+^ microglia (I). Vehicle, *n* = 43 cells (5 mice); clopidogrel, *n* = 35 cells (5 mice). (J and K) Engulfment of extinction-ensemble material. Total tdTomato^+^ puncta within microglia, partitioned into somatic versus process compartments (J), and tdTomato^+^ puncta within CD68^+^ compartments (K). Vehicle, *n* = 48 cells (5 mice); clopidogrel, *n* = 47 cells (5 mice). N.S., not significant; \**P* < 0.05, \*\*\**P* < 0.001, two-way repeated-measures ANOVA or two-tailed Student’s *t* test as appropriate.

**Figure S16.**
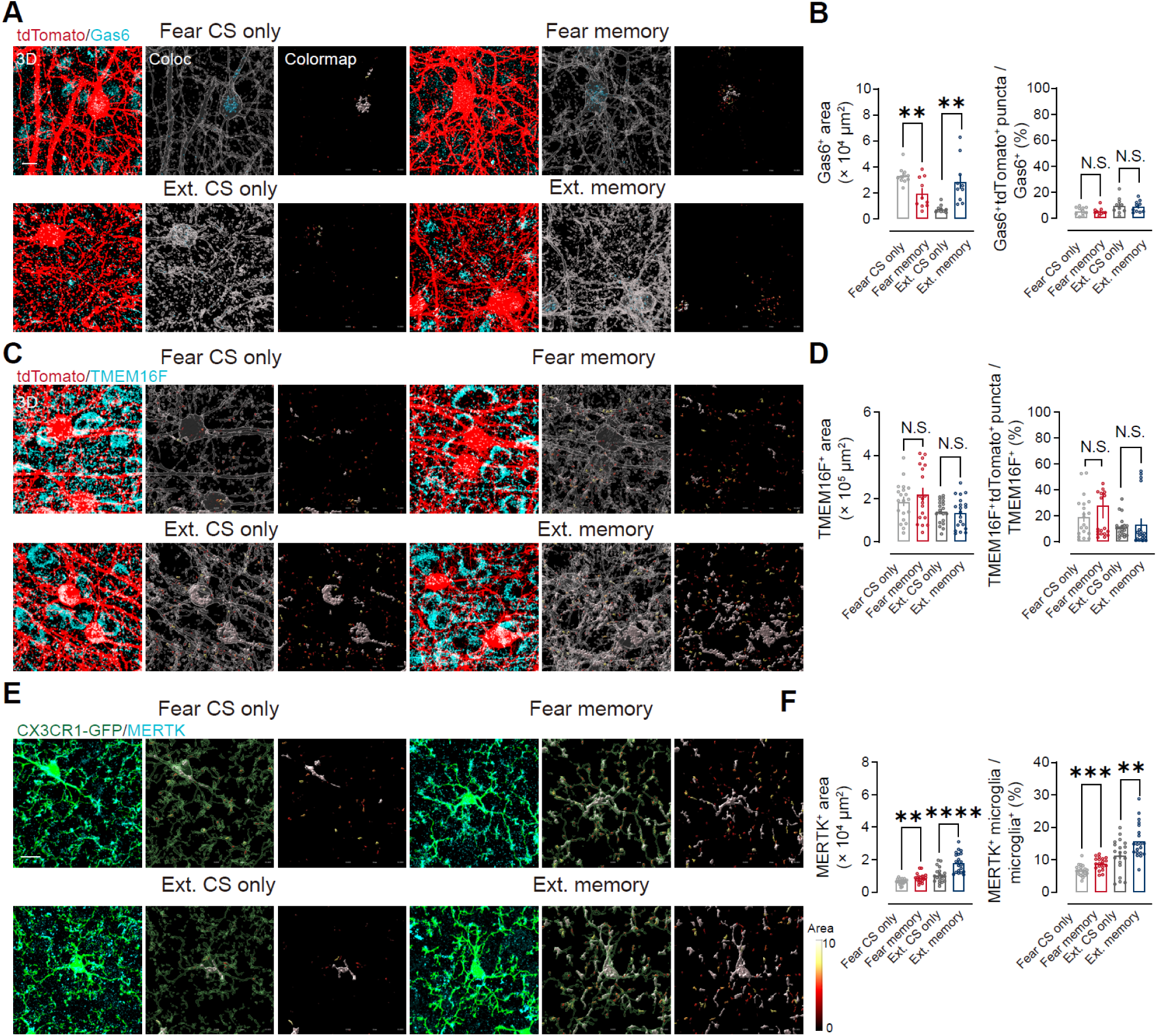
TMEM16F abundance is stable, whereas Gas6 and microglial MERTK are retrieval regulated, related to Figure 6. (A) Representative confocal images of TMEM16F immunoreactivity (cyan) and TRAP-labeled ensemble neurons (tdTomato, red) in mPFC from fear CS-only, fear retrieval (Fear memory), extinction CS-only, and extinction retrieval (Ext. memory) groups. Left, merged view; middle, TMEM16F channel; right, TMEM16F puncta overlapping TRAP signal (*Coloc*; heat map). Scale bar, 10 μm. (B) Quantification of TMEM16F signal. Left, total TMEM16F^+^ area per field. Right, fraction of TMEM16F^+^ puncta colocalized with TRAP⁺ elements. TMEM16F levels and TRAP association were unchanged by fear or extinction retrieval relative to the corresponding CS-only controls. Fear CS only, *n* = 20 sections (5 mice); Fear memory, *n* = 19 sections (5 mice); Ext. CS only, *n* = 20 sections (5 mice); Ext. memory, *n* = 20 sections (5 mice). (C) Representative confocal images of Gas6 immunoreactivity (cyan) and TRAP labeling (tdTomato, red) across the same behavioral conditions. Left, merged view; middle, 3D surface rendering of Gas6 signal (*Surfaces*); right, Gas6 puncta overlapping tdTomato^+^ elements (*Coloc*; heat map). Scale bar, 10 μm. (D) Quantification of Gas6 signal. Left, total Gas6^+^ area per field. Right, fraction of Gas6^+^ puncta colocalized with tdTomato⁺ elements. Gas6 immunoreactivity decreased after fear retrieval but increased after extinction retrieval relative to the corresponding CS-only controls, whereas Gas6-TRAP overlap remained unchanged. Fear CS only, *n* = 11 sections (5 mice); Fear memory, *n* = 10 sections (5 mice); Ext. CS only, *n* = 10 sections (5 mice); Ext. memory, *n* = 10 sections (5 mice). (E) Representative confocal images of microglia (CX3CR1-GFP, green) and MERTK (cyan; heat map) in mPFC across conditions. Left, merged view; middle, microglial surface reconstructions (*Surfaces*); right, MERTK signal within microglial volumes (*Coloc*; heat map). Scale bar, 5 μm. (F) Quantification of MERTK signal. Left, total MERTK^+^ area per field. Right, fraction of MERTK signal contained within microglia (MERTK colocalized with CX3CR1-GFP microglial volume / total MERTK). Both fear and extinction retrieval increased MERTK immunoreactivity and its enrichment in microglia relative to the corresponding CS-only controls. Fear CS only, *n* = 18 sections (7 mice); Fear memory, *n* = 20 sections (7 mice); Ext. CS only, *n* = 19 sections (7 mice); Ext. memory, *n* = 21 sections (7 mice). \*\**P* < 0.01, \*\*\**P* < 0.001, \*\*\*\**P* < 0.0001, unpaired Student’s *t* test (retrieval versus the corresponding CS-only control).

**Figure S17.**
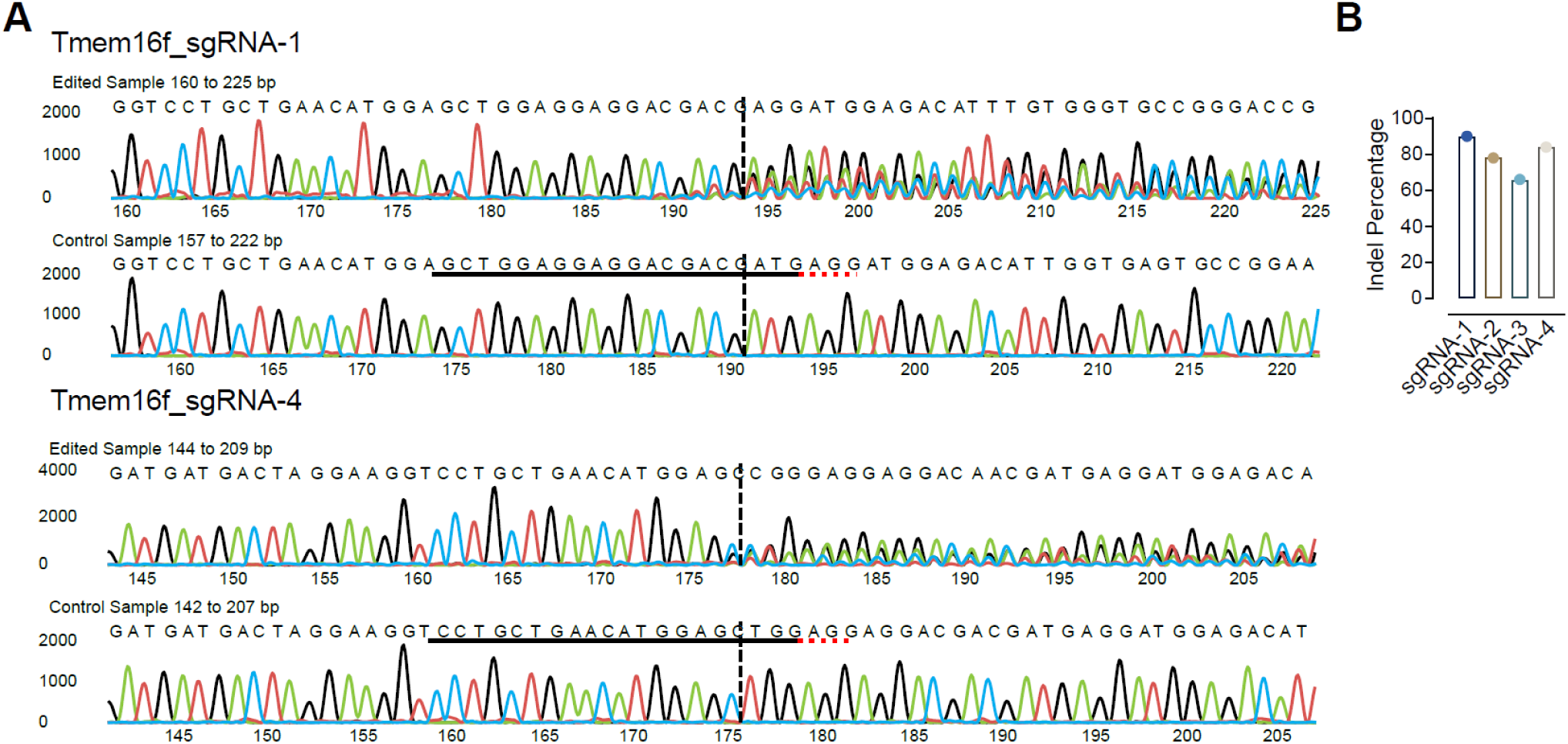
Validation of sgRNAs targeting *Tmem16f*, related to Figure 6. (A) Representative Sanger sequencing chromatograms of PCR amplicons spanning the *Tmem16f* target sites for sgRNA-1 (top) and sgRNA-4 (bottom) from edited and control samples. The sgRNA protospacer is underlined, the PAM is shown in red, and the dashed line marks the predicted Cas9 cut site. Mixed peaks downstream of the cut site in edited samples indicate indel formation. (B) Indel frequencies for four candidate *Tmem16f* sgRNAs (sgRNA-1 to sgRNA-4), estimated from Sanger-trace decomposition.

**Figure S18.**
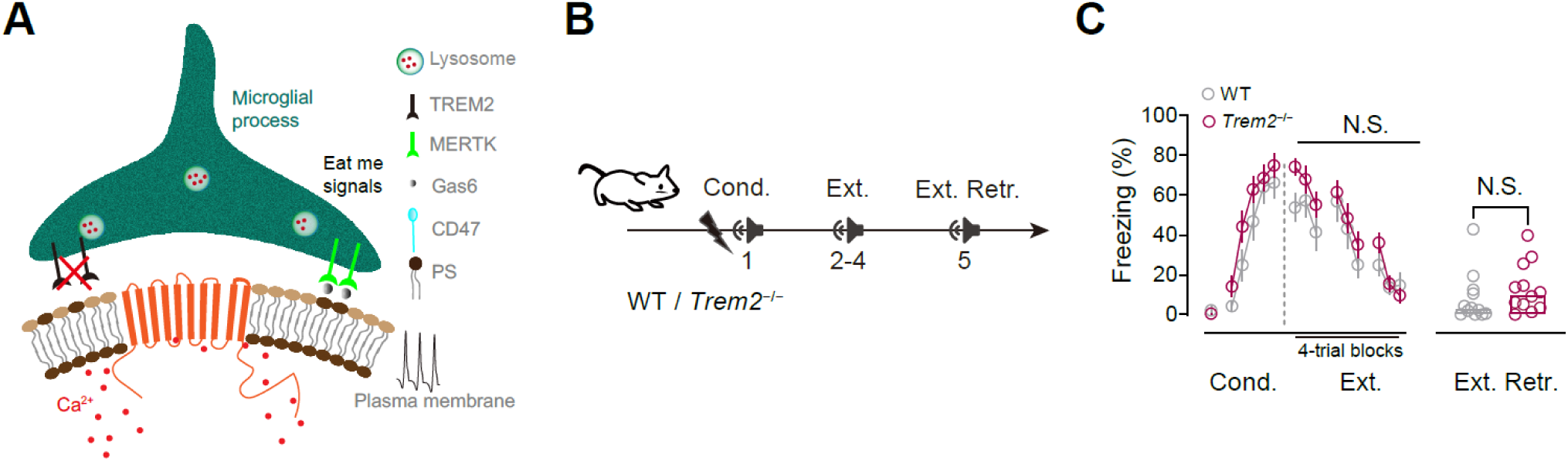
TREM2 is dispensable for fear extinction, related to Figure 7. (A) Schematic of candidate microglial phagocytosis pathways at the neuron–microglia interface. Externalized phosphatidylserine (PS) on neuronal membranes can be bridged by Gas6 to microglial MERTK (“eat-me” signaling); triggering receptor expressed on myeloid cells 2 (TREM2) is shown as an additional microglial receptor linked to phagocytic activation that was tested here. (B) Behavioral timeline for wild-type (WT) and *Trem2* knockout (*Trem2*^−/−^) mice across conditioning (Cond.; Day 1), extinction training (Ext.; Days 2–4), and extinction retrieval (Ext. Retr.; Day 5). (C) Freezing during extinction (4-trial blocks) and at extinction retrieval is indistinguishable between WT and *Trem2*^−/−^ mice. WT, *n* = 9; *Trem2*^−/−^, *n* = 11. Data are mean ± s.e.m.; N.S., not significant, two-way repeated-measures ANOVA for extinction curves; unpaired two-tailed Student’s *t* test for Ext. Retr.

**Figure S19.**
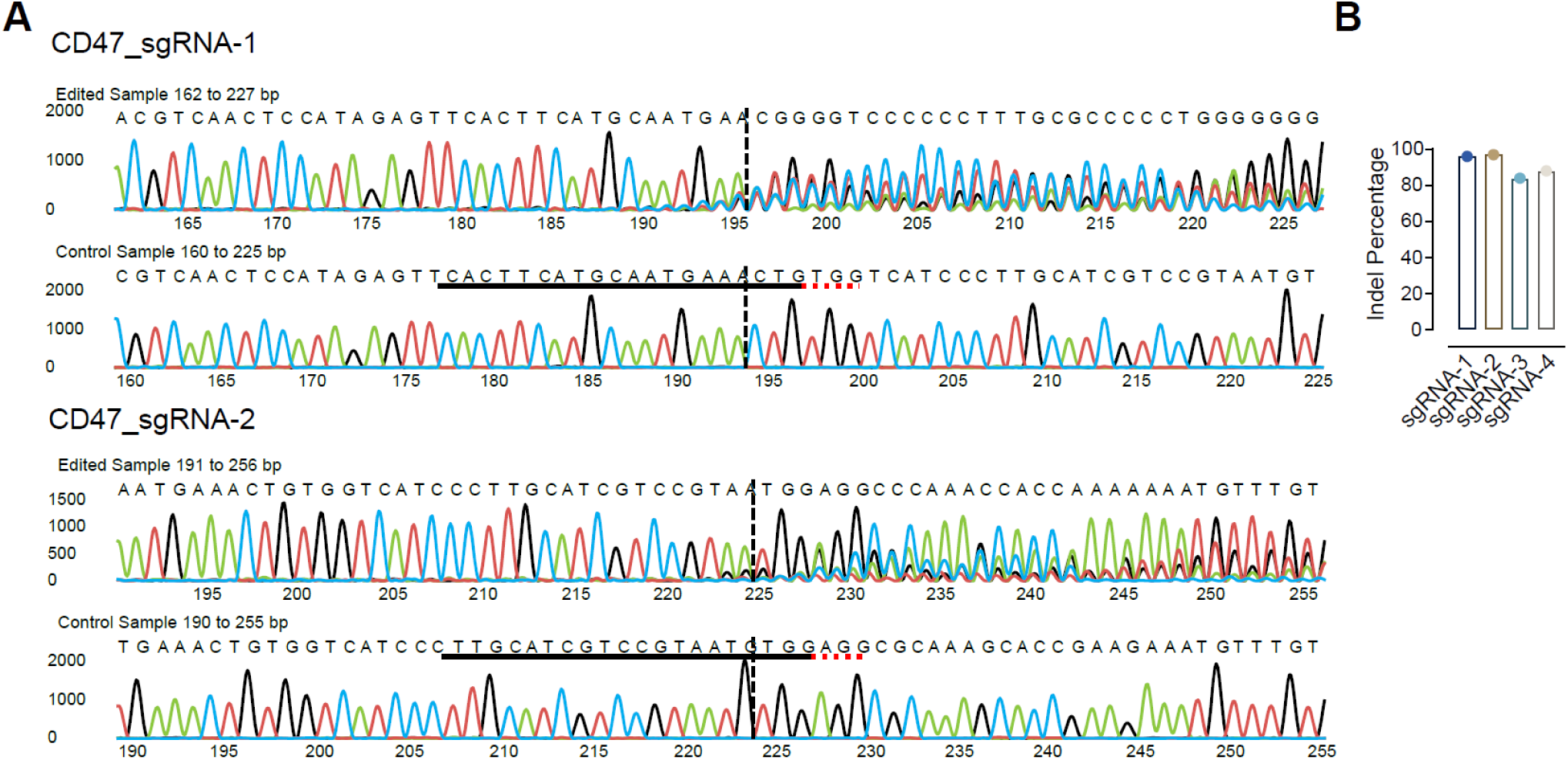
CRISPR–Cas9 editing efficiency for *Cd47* sgRNAs, related to Figure 7. (A) Representative Sanger sequencing chromatograms spanning the *Cd47* target region for sgRNA-1 and sgRNA-2, shown for edited and control samples. The sgRNA protospacer is underlined, the PAM is indicated in red, and the dashed line marks the predicted Cas9 cleavage site. Edited samples show mixed sequencing peaks downstream of the cut site, consistent with indel generation (read coordinates: sgRNA-1, 162–227 bp edited vs. 160–225 bp control; sgRNA-2, 191–256 bp edited vs. 190–255 bp control). (B) Quantification of indel frequency for four *Cd47* sgRNAs (sgRNA-1 to sgRNA-4).

